# Internal Phase Separation in Synthetic DNA Condensates

**DOI:** 10.1101/2025.03.19.644034

**Authors:** Diana A. Tanase, Dino Osmanović, Roger Rubio-Sánchez, Layla Malouf, Elisa Franco, Lorenzo Di Michele

## Abstract

Biomolecular condensates regulate cellular biochemistry by organizing enzymes, substrates and metabolites, and often acquire partially de-mixed states whereby distinct internal domains play functional roles. Despite their physiological relevance, questions remain about the principles underpinning the emergence of multi-phase condensates. Here, we present a model system of synthetic DNA nanostructures able to form monophasic or biphasic condensates. Key condensate features, including the degree of interphase mixing and the relative size and spatial arrangement of domains, can be controlled by altering nanostructure stoichiometries. The modular nature of the system facilitates an intuitive understanding of phase behavior, and enables mapping of the experimental phenomenology onto a predictive Flory-Huggins model. The experimental and theoretical framework we introduce will help address open questions on multiphase condensation in biology and aid the design of functional biomolecular condensates *in vitro*, in synthetic cells, and in living cells.

## 1 Introduction

Cellular condensates and membrane-less organelles invariably comprise multiple macromolecular species, and often display distinct sub-compartments, as observed for nucleoli [1], P-granules, and stress granules [2–4]. These internal domains are known to play functional roles, *e*.*g*. in the sequential processing of ribosomal RNA occurring in nucleoli [1, 5, 6]. Yet, the mechanisms underpinning their formation and characteristics remain poorly understood due to the complexities of the intracellular environments.

Synthetic condensate models, whereby macromolecular makeup and interactions can be precisely controlled, constitute an ideal tool to unravel the biophysics of multi-phase condensation. Several examples of synthetic condensate-forming constructs have been proposed, primarily reliant on engineered proteins condensing thanks to low complexity domains or multivalent binders [7–14]. While model multiphase condensates can be constructed with protein-based systems [15], difficulties with programming protein-protein interactions, particularly if mediated by disordered domains, make it challenging to perform systematic studies and draw general conclusions.

Alongside proteins, nucleic acids are common constituents of biological condensates, often recruited as clients but sometimes playing a structural role [16–19]. In contrast to proteins, however, nucleic acids benefit from well understood and selective base-pairing, which has led to powerful techniques for designing synthetic DNA and RNA constructs with near-arbitrary properties [20–23]. Leveraging the capabilities of DNA and RNA nanotechnology, multiple platforms have emerged to engineer “designer” condensates either using long DNA chains [24, 25] or small, branched, “nanostar” constructs [26–39]. DNA nanostars (NSs), in particular, constitute a valuable model to investigate condensation and liquid-liquid phase separation (LLPS), owing to the precise control they afford over valency and interaction strength. These constructs typically feature three to six double-stranded (ds)DNA “arms”, with programmable length and flexibility [31,40,41]. Arms terminate with adhesive moieties such as hydrophobic groups [40, 42] or, more commonly, single-stranded (ss)DNA “sticky ends” (SE). The phase behaviour of single-component NS mixtures has been thoroughly investigated, first with Biffi *et al*., who explored LLPS in trivalent and tetravalent DNA NSs [43], and more recently with studies on penta- and hexavalent NSs [44], the effect of ionic strength [45], and that of altering arm length [31].

Thanks to base-pairing selectivity, binary (or multi-component) NS mixtures with orthogonal SEs assemble into distinct condensates that do not mix [30, 34, 46]. Multiphase condensates can then be produced by adding “linker” NSs, with sticky ends complementary to two other NS species [30,39,46,47]. Disrupting the linkers *via* strand displacement [35,48,49] or enzymatic digestion [46, 47] can then trigger condensate de-mixing, resulting in division-like events valuable as readouts for molecular computation or biosensing [48]. Despite these remarkable demonstrations of functionality, the potential of multicomponent NS mixtures as means of systematically investigating multiphase condensation remains largely untapped. Progress has been hindered by the nature of the design solutions reported to date, which often combined direct NS-NS interactions with those mediated by (multivalent) linkers, making fine-tuning of the relative affinities challenging [30, 37, 39].

Here we introduce a modular system designed to facilitate the systematic exploration and rationalization of LLPS in binary DNA NS mixtures. The system comprises two non-interacting NS species and three divalent linkers, mediating both intra- and inter-species interactions between the NSs species. NS-NS interactions can be modulated by changing linker stoichiometry, leading to monophasic or biphasic DNA condensates with diverse morphologies and compositions. By systematically exploring the compositional parameter space, we locate the boundary between monophasic and biphasic behaviors and identify an order parameter governing the transition. We further demonstrate that experimental phase behavior can be mapped onto a ternary Flory-Huggins model [50–52], in which the action of linkers as the mediators of NS-NS affinity is described implicitly, providing a predictive tool for navigating the multi-component parameter space. Finally, we explore the effect of changing thermal annealing protocols on phase behavior, highlighting intrinsic challenges with achieving thermodynamic equilibration in slowly-relaxing NS phases [53].

The quantitative and systematic understanding afforded by our experimental model system, and the associated theoretical framework, will facilitate the rationalization of multicomponent biomolecular condensation in biology. Additionally, the improved programmability enabled by our platform will pave the way for deploying multiphase DNA condensates in bioprocessing [8], biosensing [48], and synthetic cell engineering [54–56].

## 2 Results

### 2.1 System Overview

**Figure** 1**A** summarizes the design and self-assembly protocol of the condensate-forming DNA nanostructures. The system comprises five building blocks: two four-arm NSs, dubbed A and B, and three linear linkers, dubbed aa, bb, ab. Each NS self-assembles from four unique strands, resulting in 20 base-pair (bp) dsDNA arms, each terminated by a 6 nucleotide (nt) ssDNA SE. An unbound thymine separating neighbouring arms enhances junction flexibility [28, 41, 43] and helps to destabilize stacked “X” conformations [57]. One in ten NSs is fluorescently labelled with either ATTO 488 (NS A) or Alexa 647 (NS B), to enable fluorescent imaging. SE sequences – *α* on NS A and *β* on NS B – are designed to be neither mutually- nor self-complementary. Linkers assemble from two strands, producing a 20 bp duplex with two SEs. Linkers aa and bb feature two identical *α*^*^ and *β*^*^ SEs, respectively, while ab includes one *α*^*^ SE and one *β*^*^ SE, with “* “indicating complementarity to SEs on the NSs. Hence, in our system, linker aa mediates attractive interactions between otherwise non-interacting A NSs, bb mediates attractive interactions between B NSs, and ab mediates cross-species binding. SE sequences were designed to ensure similar hybridization free energy, estimated as − 8.98 kcal mol^−1^ and − 10.25 kcal mol^−1^ for *α*/*α*^*^ and *β*/*β*^*^, respectively (NUPACK [58], 25°C, 2 µM, 0.3 M Na^+^). The SEs match affinities previously used in the literature [28, 30, 43, 45, 46], ensuring sufficient stability to enable condensation at room temperature while retaining a liquid-like droplet behavior. We expect that variations in SE affinity within these broad constraints would produce qualitatively similar outcomes but, as discussed later, condensate morphology and phase composition may be quantitatively different. All oligonucleotide sequences are provided in Table S1.

**Figure 1:**
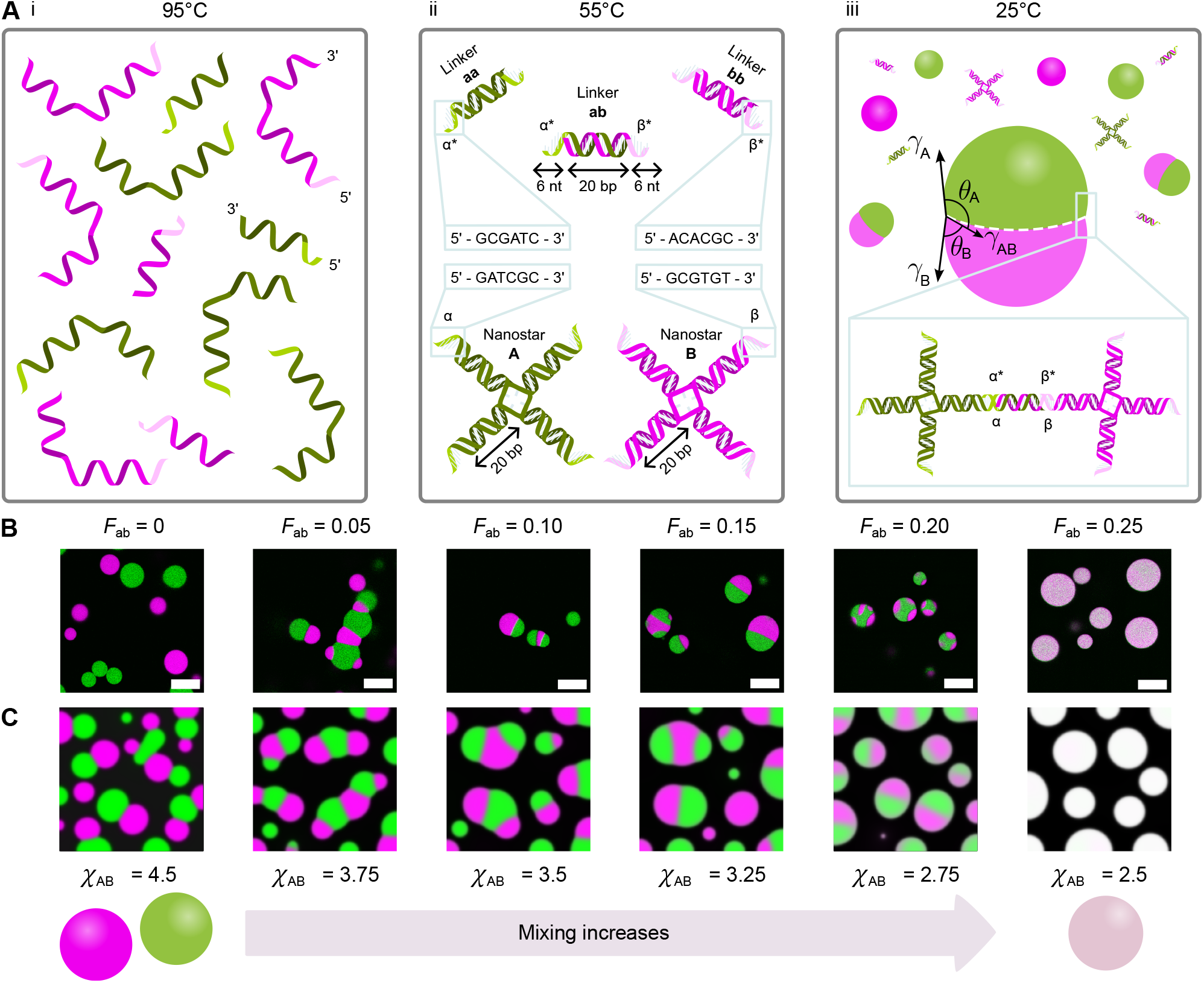
Synthetic DNA condensates with programmable internal phase separation can be engineered using five DNA building blocks: two orthogonal nanostars and three linkers. The interactions between four-arm DNA nanostars (NSs) – A (shown in green) and B (shown in magenta) – are controlled by three types of duplex linkers (aa, bb, ab) that mediate A-A, B-B, and A-B interactions through complementary sticky ends (SEs), denoted as *α/α*^*^ and *β/β*^*^. (**A**) Summary of the two-stage annealing process. i) At high temperatures (above 80 °C), all strands are fully dehybridized. ii) As the temperature decreases below the melting temperature of the individual NSs and linkers, the five building blocks (A, B, aa, bb, and ab) self-assemble. Both SE pairs are designed to have the same %GC content and are 6 nt in length. iii) Condensates are prepared by mixing the five components and annealing from 55 °C – above the melting temperature of the condensates. As the temperature is decreased, complementary SEs on NSs and linkers bind, leading to condensation. In de-mixed condensates, an A-rich phase and a B-rich phase can be identified along with corresponding contact angles, *θ*_*A*_ and *θ*_*B*_. (**B**) Confocal micrographs (scale bars are 20 µm) show how the affinity between the A and B populations can be tuned by controlling the fraction of ab linkers, *F*_ab_, to yield pure condensates (*F*_ab_ = 0), increasingly mixed biphasic droplets (0.05 ≤ *F*_ab_ ≤ 0.20), and fully mixed condensates (*F*_ab_ = 0.25). Confocal micrographs are rescaled between minimum and maximum pixel intensity to aid visualization. (**C**) Simulation snapshots produced with an effective Flory-Huggins model with three components – A NSs, B NSs and the surrounding solvent – whereby the effect of linkers is implicitly encoded. Changing the effective interaction parameter *χ*_*AB*_, describing the affinity between A and B NSs, produces morphologies similar to those found experimentally. Equivalent parameters for A-A and B-B interactions are fixed to *χ*_*AA*_ = 4 and *χ*_*BB*_ = 4. The total packing fraction of A and B components is set to *ϕ*_*A*_ = *ϕ*_*B*_ = 0.23 [see Supplementary Note I, Supplementary Information].

Assembly of individual NSs and linkers was induced by annealing separately the relevant constituent strands from 95°C, ensuring initial denaturation of the dsDNA domains (Figure S1, Table S2, and Experimental Section). The constructs, whose correct assembly was verified by agarose gel electrophoresis (Figure S2), were then mixed with the desired concentration ratios. Finally, condensate formation was induced through a second, slow thermal annealing starting at 55°C – a temperature chosen to ensure melting of any NS-linker aggregates (as discussed below) but insufficient to disassemble the individual constructs (Figure S1, Table S2).

As sketched in Figure 1**A**, progressively lowering the temperature from 55°C causes the linkers to cross-link the NSs, triggering LLPS and the formation of DNA condensates. The monophasic or biphasic character of the condensates is controllable through their NS and linker composition. A transition from fully de-mixed A-type (green) and B-type (magenta) droplets, to partially-mixed branched or “Janus” morphologies, and then fully mixed droplets, is observed upon increasing the fraction of ab linkers (*F*_ab_), while keeping constant the total linker concentration (Figure 1**B**).

### 2.2 Flory-Huggins Modeling

The experimental results in Figure 1**B** can be captured by a simple computational model based on the Flory-Huggins (F-H) theory of phase separation, which only requires three components.

A full theoretical description of the experimental system, including two NSs, three linkers, and the surrounding solvent, is challenging, and would include details such as NS valency, the number and orientation of linkers connected to the NS arms, and excluded volume effects. Even a phenomenological F-H model describing all six components is rather complex, as we discuss in Supplementary Note I (Supplementary Information). In Supplementary Note I, we show that our model leads to the familiar phases that are observed in the experiments: a phase rich in A and B NSs or two different phases richer in A or B, respectively. Of particular interest, however, is whether there are any redundant degrees of freedom which are included in more complete descriptions of field theories of this type, and that can be coarse-grained while conserving a phase behavior similar to experiments. To explore this possibility, we show that F-H theories in multiple fields can be shown to be identical to models with fewer fields when two components are sufficiently attractive (Supplementary Note I). In other words, we find that strongly attractive interactions are suggestive of redundant degrees of freedom. The experiments are also indicative of the fact that, despite the larger number of components, A and B droplets behave as a binary mixture of NSs with tunable interactions. We thus pose the following question: To what extent, in the context of a F-H theory, can linkers be modeled as effective modifiers of interaction parameters between A and B NSs, without having to take into account the greater degree of complexity of introducing a full phase description of linker behavior? To address this question, we propose a ternary free energy density, f, describing the A and B NSs and the solvent, with the effect of linkers as the mediators of NS-NS interactions being encoded by three interaction parameters (χ_AA_, χ_BB_, and χ_AB_) [50, 52]:

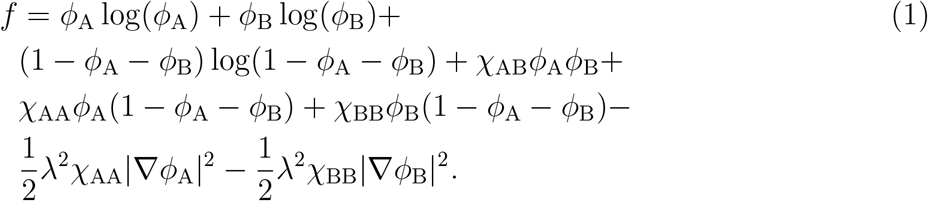

In Equation 1, *ϕ*_A_ and *ϕ*_B_ are the volume fractions of components A and B, respectively, while λ defines the width of the interface between the A-rich and B-rich phases and the solvent, and is set to 1 in our simulations. In Supplementary Note I, we show that we can derive approximate relationships between χ_AA_, χ_BB_, and χ_AB_ and linker concentrations. For simplicity, we ignore the contribution to the free energy resulting from the interfaces between the A-rich and B-rich phases.

We use the Cahn-Hilliard equation [51] to model the time evolution of the field theory in Equation 1, and solve numerically for different values of χ_AB_ and fixed χ_AA_ and χ_BB_, as outlined in Supplementary Note I. In Figure 1**B** and **C** we show that the F-H description qualitatively replicates experimental patterns. Additional simulations snapshots are shown in Figure S3. A more systematic comparison is provided in the following sections, aimed at verifying whether our simplified theory is capable of quantitatively describing the experimental phase behaviour. We note that, recently, Cappa *et al*. modelled monophasic and biphasic DNA-NS condensation in monophasic and biphasic by combining the Cahn-Hilliard equation with a free energy functional derived from the Wertheim theory [39], obtaining qualitative agreement with experimental observations in literature [30, 39].

### 2.3 Mapping Phase Behavior

The experimental system has five free parameters, namely the concentrations of all NSs and linkers. Mapping phase behavior across this high-dimensionality parameter space is experimentally unfeasible, and beyond the scope of this work. To prevent the combinatorial explosion of the parameter space, we impose a stoichiometric constraint on the concentrations of SEs, namely that the overall concentration of one SE type on NSs is equal to that of the complementary SE on linkers: [α] = [α^*^] and [β] = [β^*^]. The constraint, equivalent to imposing charge neutrality in an electrostatic system, can be expressed in terms of the concentrations of the individual constituents as

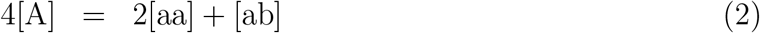

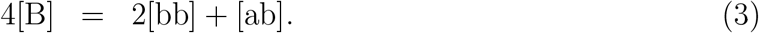

Equations 2 and 3 imply [linker] : [NS] = 2 : 1, where [linker] = [aa] + [bb] + [ab] and [NS] = [A] + [B]. We further constrain the system by working at constant overall NS (and linker) concentration, choosing [NS] = 0.5 µM and [linker] = 1 µM. Once these constrains are imposed, the system is left with two free parameters, which we chose as the molar fraction of NS A, F_A_ = [A]/[NS], and the molar fraction of the ab linker, *F*_ab_ = [ab]/[linker]. The molar fractions of the remaining components are then determined by [see Supplementary Note II, Supplementary Information, for derivation]

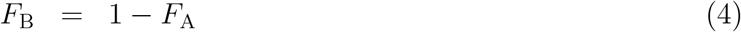

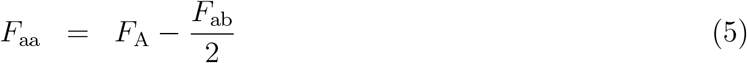

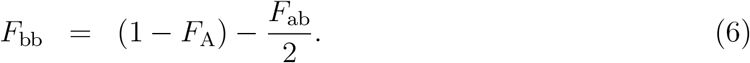

**Figure** 2**A** maps the phase behavior of the mixture with 63 different compositions spanning the *F*_ab_ × *F*_A_ space, with higher magnification confocal micrographs of the condensates and associated scale bars shown in Figure S4. Representative full fields of view are shown in Figures S5-S13. Figure S14 graphically depicts the linker composition of all samples in Figure 2**A**, as determined by Equations 5 and 6.

**Figure 2:**
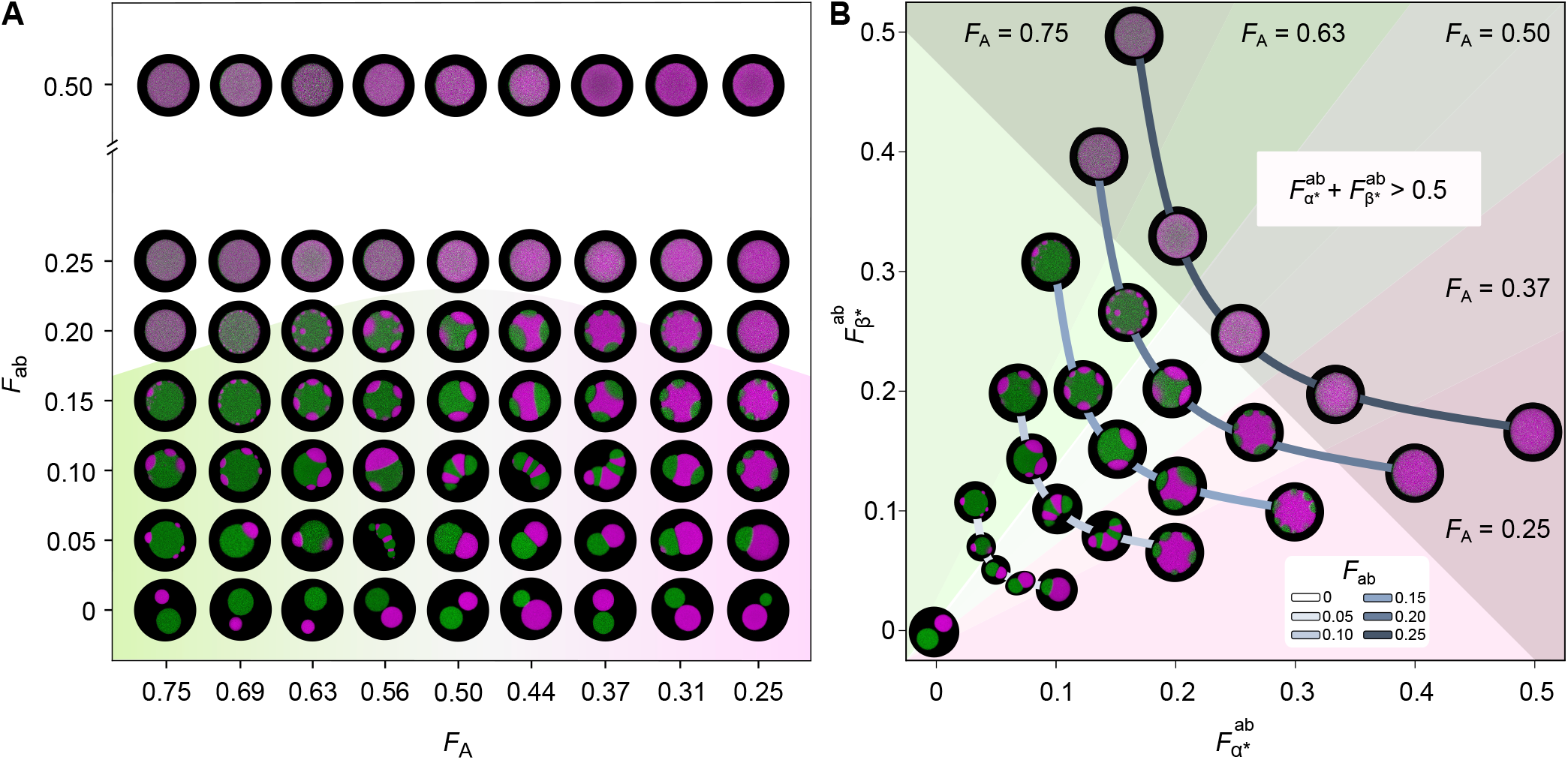
Phase diagrams with selected confocal micrographs showing the variety of morphologies accessible with the system of DNA NSs and linkers. (**A**) Phase diagram mapping the *F*_ab_ × *F*_A_ space, where *F*_ab_ and *F*_A_ = 1 − *F*_B_ are the fractions of ab linkers and A NSs, respectively [see main text]. The transition from biphasic to monophasic condensates occurs at lower *F*_ab_ when *F*_A_ ≠ *F*_B_ (*F*_A_ ≠ 0.5). The shaded region highlights biphasic condensates. (**B**) Alternative phase diagram with a subset of the micrographs shown in panel **A** for five different values of *F*_***A***_, in the 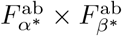 space, where 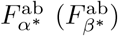 is defined as the fraction of *α*^*^ (*β*^*^) SEs present on ab linkers, as opposed to aa (bb) linkers. The monophasic to biphasic transitions occurs when 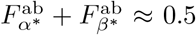. Shaded regions mark monophasic/biphasic domains and different *F*_A_ values. Grey lines connect samples with the same *F*_ab_. For both panels, micrographs were segmented using a mask to remove background. Accompanying scale bars can be found in the Supplementary Information, Figure S4 while Figures S5-S13 show larger microscopy fields of view.

Expectedly, we observe that *F*_A_ controls the relative abundance of the A-rich and B-rich phases in biphasic condensates, while *F*_ab_ largely governs the transition between biphasic condensates, found at low *F*_ab_, and single-phase morphologies (Figure 2**A**). We note, however, that the monophasic-to-biphasic transition occurs at different values of *F*_ab_ depending on *F*_A_: samples asymmetric in their NS composition require a lower relative concentration of the ab linker to produce fully mixed condensates. To rationalize this trend, we note that as *F*_A_ approaches 0 or 1, the proportion of the minority ‘cis’ linker (aa or bb, respectively) decreases relative to that of the ab linker (Figure S14). In turn, this implies that the minority NS component becomes increasingly more likely to bind ab linkers compared to aa or bb, which facilitates its solubilization in the majority NS phase and the emergence of monophasic condensates.

Following this observation, we introduce two different system parameters:

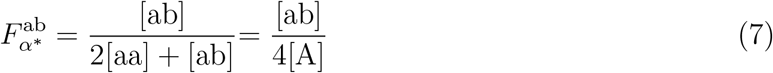

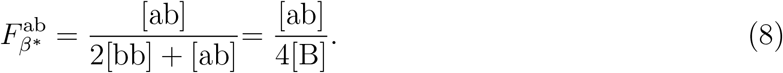

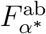 and 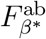 indicate, respectively, the fraction of *α*^*^ or *β*^*^ SEs present on ab linkers compared to aa or bb linkers. More specifically, 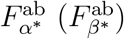 can be interpreted as the fraction of sticky ends in type A (B) NSs involved in bonds with B (A) NSs, as opposed to binding NSs of the same kind.

Figure 2**B** shows a sub-set of the samples in Figure 2**A** distributed on the 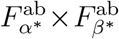 plane, highlighting that the biphasic to monophasic transition occurs at constant 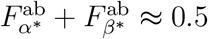 [see Figure S4 for larger micrographs and scale bars]. To explain this behavior, let us consider the case of an asymmetric sample with *F*_A_ = 0.75, for which the biphasic to monophasic transition occurs when 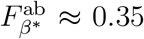 and 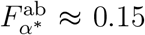 (Figure 2**B**). Here, because A NSs are more abundant than B NSs, a relatively low likelihood that A NSs form trans bonds with B NSs – approximated by 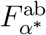 – is sufficient to solubilize B NSs, while a higher 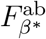 can be tolerated while retaining biphasic character. This reasoning becomes more intuitive if one considers the role of A-B bonds in stabilizing domain interfaces in biphasic condensates. When *F*_A_ (*F*_B_) decreases, the fraction of A NSs (B NSs) located at the interface between A- and B-rich domains increases, and so must do the fraction of A-B bonds relative to all bonds formed by A (B) NSs, namely 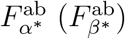.

We thus hypothesize that, compared to the fraction of linkers *F*_AB_, the heuristic order parameter 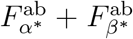 represents a better predictor for the tendency of the mixture to acquire monophasic or biphasic morphologies, which accounts for the effect of changing NS stoichiometry. It should further be noted that, for symmetric samples with *F*_A_ = *F*_B_ = 0.5, and therefore 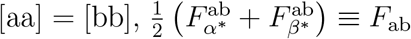. This equivalence places the threshold for the monophasic-to-biphasic transition at *F*_AB_ ≈ 0.25, implying that, for complete mixing to occur, a NS of type A (B) needs to form, on average, at least one bond with a NS of type B (A).

We then proceed to analyze the morphological characteristics of the condensates, including phase composition and interfacial tensions, through an image segmentation pipeline applied to confocal micrographs (**Figure** 3**A**), presented in detail in Experimental Section and Figure S15-S18. To assess the degree of mixing in biphasic condensates, we use the partition coefficients, *ρ*_A_ and *ρ*_B_. These are computed as the ratio between the fluorescence intensity of a given NS type (A or B) sampled in the phase depleted of that NS type (B-rich or A-rich phase, respectively) and the intensity sampled in the phase enriched in the NS type (A-rich or B-rich phase, respectively). As a result, *ρ*_A_ and *ρ*_B_ tend to 0 for fully de-mixed condensates and approach 1 for fully mixed (monophasic) condensates [30,33,37,38,46]. Figure 3**B** shows *ρ*_A_ and *ρ*_B_ as a function of *F*_ab_, demonstrating a high degree of de-mixing at low fractions of ab linker and a sharp rise as full mixing is approached, as reported in previous studies with multivalent linkers [37, 38, 46]. Remarkably, plotting the data as a function of 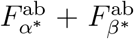 collapses the curves, causing them to approach the biphasic-to-monophasic phase boundary 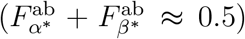 with similar trends, and confirming the relevance of this expression as a meaningful order parameter for the de-mixing transition (Figure 3**C**). The distributions of *ρ*_A_ and *ρ*_B_ values for all samples can be inspected in Figure S19, while information on the numbers of condensates analyzed are collated in Table S3.

**Figure 3:**
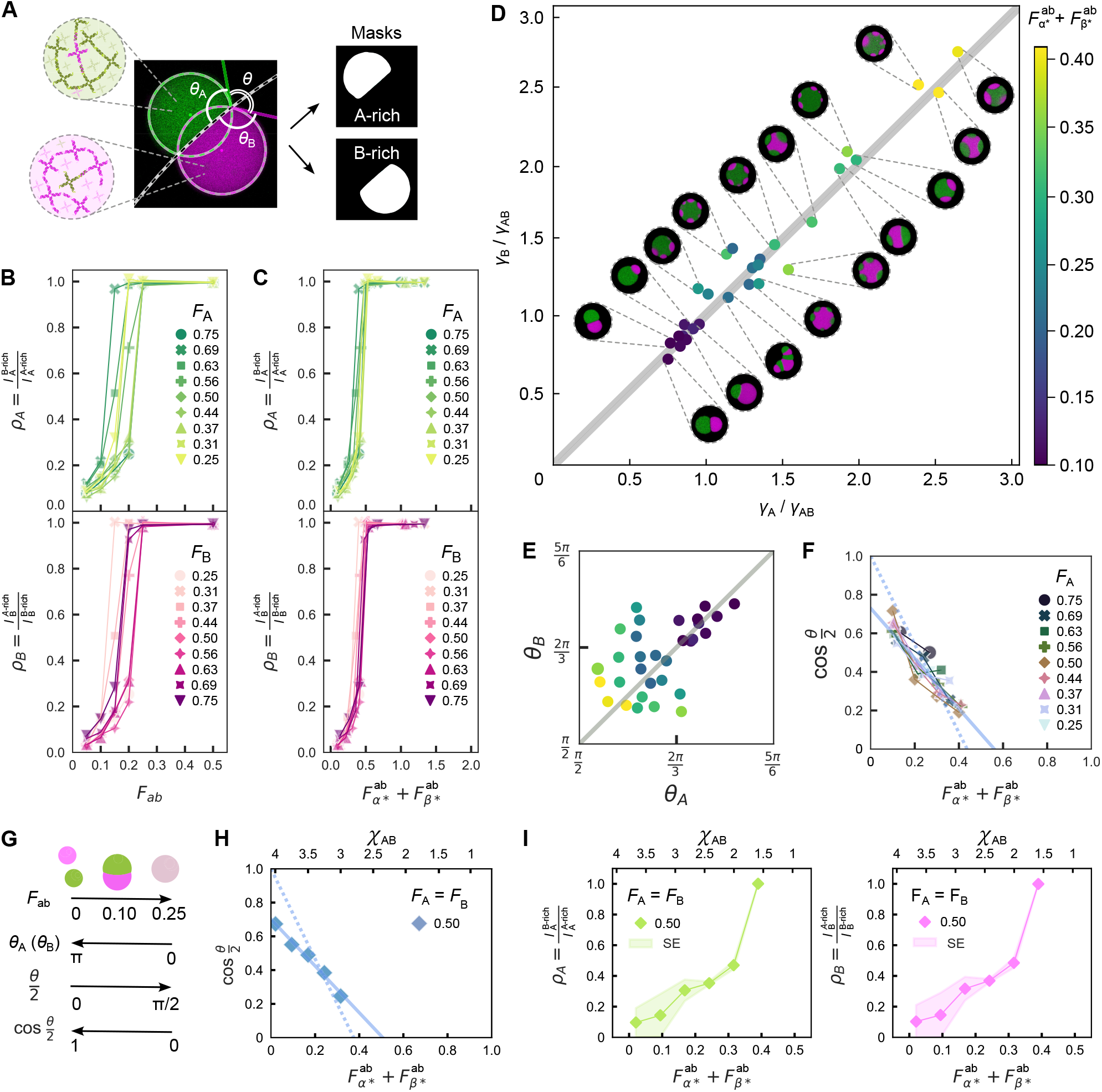
Quantitative analysis of confocal micrographs and simulations snapshots shows how partition coefficients, contact angles, and interfacial tensions change with nanostructure stoichiometry. (**A**) Example of the segmentation routine used to identify the A-rich and B-rich domains in a Janus-like droplet. Masks are used to determine relative intensities in each of the two channels and compute the partition coefficients, *ρ*_A_ and *ρ*_B_. *ρ*_A_ (*ρ*_B_) describes the relative abundance of NS A (or B) in the B-rich (A-rich) phase over the A-rich (B-rich) phase [see main text and Experimental Section]. (**B**) Both *ρ*_A_ (top, green) and *ρ*_B_ (bottom, magenta) increase with increasing *F*_ab_ for all *F*_A_ and *F*_B_, indicating a progression from highly de-mixed biphasic condensates (low *ρ*_A_, *ρ*_B_) to fully mixed monophasic droplets (*ρ*_A_ = *ρ*_B_ = 1). (**C**) Partition coefficient data collapse when plotted as a function of the empiricial order parameter 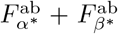. (**D**) Scatter plot showing the correlation between the interfacial intension ratios *γ*_A_*/γ*_AB_ and *γ*_B_*/γ*_AB_, computed as discussed in the main text. Representative confocal micrographs are also shown in Figure 2. (**E**) Analogous scatter plot of the contact angles *θ*_A_ *vs θ*_B_ [see **A**]. In **D** and **E**, marker color reflects the empirical parameter 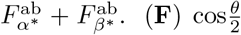 [see **A**], proportional to *γ*_AB_, decreases with increasing 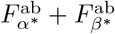, independently of *F*. Solid and dashed lines are linear fits, the latter constrained to 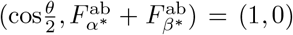. (**G**) Graphical summary of the relationship between condensate morphology, composition, and contact angles. (**H**) Simulations results show 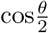 decreasing linearly with decreasing *χ*_*AB*_. Comparing a linear fit to that of experimental 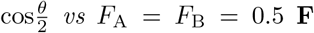, enables mapping of *χ*_AB_ onto 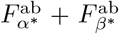. (**I**) Simulated partition coefficients *vs χ*_AB_ and *F*_A_ = *F*_B_ = 0.5 (calibrated as in **H**). While values are very similar, statistical variations lead to marginally different *ρ*_A_ and *ρ*_B_ values. Shaded regions indicate standard errors across condensates in a given simulation snapshot. In **H** and **I**, contact angles and partition coefficients are obtained from simulated snapshots (as shown in Figure 1 for de-mixed samples with *χ*_AA_ = *χ*_BB_ = 4, *ϕ*_A_ = *ϕ*_B_ = 0.23) using the same segmentation pipeline used for experiments [see Experimental Section]. All experimental data points shown in panels **B**-**F** represent the median of the condensate populations for a given sample composition, imaged in 2-3 fields of view. The distributions of partition coefficients, contact angles, and interfacial tensions are provided in Figure S19-21. The number of condensates analyzed from multiple fields of view for each composition are reported in Tables S4 and S5.

For each A-B interface in biphasic structures, fitting the segmented contours with circles allows us to determine two values for the contact angles of the A-rich and B-rich phases, which are averaged to compute *θ*_A_ and *θ*_B_ (Figure 3**A**). Figure S20 shows the distributions of *θ*_A_ and *θ*_B_ for all sample compositions in Figure 2, expectedly demonstrating a decrease with increasing *F*_ab_, which becomes steeper for *F*_A_ approaching 0 or 1.

In a system of three interacting fluids, here the A-rich condensate phase, the B-rich condensate phase, and the surrounding water-rich phase, the contact angles are linked to interfacial tensions through the Neumann stability criterion [30, 59]:

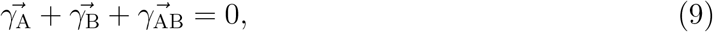

where the orientations of the three vectors are tangential to the interfaces, as depicted in Figure 1**A** *iii*, and their magnitudes are equal to the interface tension between the A-rich phase and the solvent (*γ*_A_), the B-rich phase and the solvent (*γ*_B_), and the two condensate phases (*γ*_AB_). Using Equation 9, one can calculate interface-tension ratios from the contact angles, specifically [59, 60]:

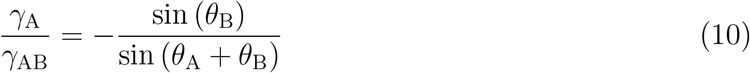

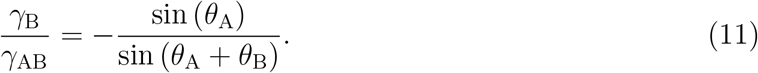

Distributions of the measured *γ*_A_/*γ*_AB_ and *γ*_B_/*γ*_AB_ are shown in Figure S21, while information on the number of sampled interfaces is provided in Table S4. In Figure 3**D**, we present a scatter plot of *γ*_A_/*γ*_AB_ *vs γ*_B_/*γ*_AB_, demonstrating that the interfacial tensions between the two dense phases and the solvent are similar across the range of tested compositions, as expected given the similar hybridization free energy of α/α^*^ and β/β^*^ SEs. Deviations from the *γ*_A_ ~ *γ*_B_ symmetry line, more evident when plotting *θ*_A_ *vs θ*_B_ (Figure 3**E**), are largely ascribed to segmentation inaccuracies, expected to be particularly prominent in compositionally asymmetric samples where one of the two phases forms domains with dimensions approaching the diffraction limit. Both *γ*_A_/*γ*_AB_ and *γ*_B_/*γ*_AB_ increase with increasing 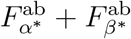 and the system approaching the single-condensed-phase regime. Note that the condition *γ*_A_ ~ *γ*_B_ prevents the emergence of core-shell architectures with an A-rich (B-rich) core being encased in a B-rich (A-rich) shell, which would require *γ*_B_ + *γ*_AB_ < *γ*_A_ (*γ*_A_ + *γ*_AB_ < *γ*_B_) [51]. These morphologies may be accessed by re-engineering NS to break the surface tension symmetry, *e*.*g*. by changing SE length or NS arm length [33].

Under the simplifying assumption that *γ*_A_ = *γ*_B_ ≡ *γ*_0_, Equation 9 yields

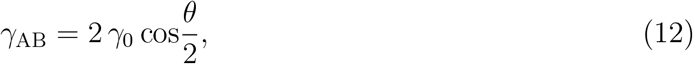

where *θ* = 2π − *θ*_A_ − *θ*_B_ (Figure 3**A**). Figure 3**F** shows that, when plotted against the heuristic order parameter 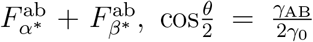 data for all tested *F*_A_ values collapse onto a decreasing linear trend [30]. Fitting and extrapolation indicate that 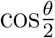 should approach zero for 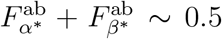, consistent with the interfacial tension *γ*_AB_ becoming vanishingly small as the mixing transition is approached. To facilitate data interpretation, Figure 3**G** summarizes the mutual relationships between contact angles, interface tensions and condensate morphology.

To quantitatively compare numerical results with experiments, we processed simulated snapshots shown in Fig 1**C** (time point 5) with the same image segmentation pipeline used for the experiments, extracting 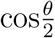 (Figure 3**H**), *ρ*_A_ and *ρ*_B_ (Figure 3**I**) as a function of the F-H interaction parameter χ_AB_. Consistent with experiments, 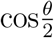 linearly tends to zero as χ_AB_ approaches the critical value for mixing. Comparison between the linear fit of 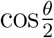 in experimental and simulated data, allows us to map χ onto 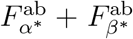. Specifically, a linear relationship between the parameters can be identified:

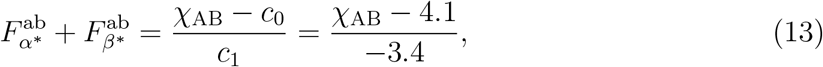

where values for the constants c_0_ and c_1_ have been pinpointed by equating the linear fits to experimental and simulated data. Importantly, while fitting was conducted on contact angle data, this relationship produces good agreement between the dependence of the partition coefficients *ρ* and *ρ* on 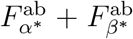 observed in simulations (Figure 3**I**) and experiments (Figure 3**B** and **C**), confirming the predictive value of the Flory-Huggins model.

We acknowledge that slight deviations from the intended sample stoichiometry may occur due to experimental error. We can roughly estimate these errors to be smaller than the incremental 5% steps used when varying *F*_ab_, given the observation of robust trends as a function of this variable.

### 2.4 Effect of Mixture Composition on Condensate Melting Temperature

To facilitate equilibration, samples discussed in Figure 2 and 3 were slowly annealed from *T* = 55 ^°^C to 25 ^°^C. This temperature range was chosen to enclose the melting temperature, *T*_m_, of the condensates, *i*.*e*. 25 ^°^C < *T*_m_ < 55 ^°^C. The condensate melting temperatures were measured by performing cooling and heating ramps while conducting epifluroescence imaging using a thermal microscope stage, as discussed in Experimental Section and shown in **Figure** 4**A** and Figure S22**a**. Detailed snapshots capturing condensates on a cooling ramp are shown in Figure S23. We then computed the ratio between the standard deviation and the mean of the pixel values for images collected in both the A NS and B NS fluorescent channels. The resulting traces, shown for selected sample compositions in Figure 4**B** and for all tested compositions in Figure S22**b**, exhibit sharp slope changes coinciding with condensate formation and melting, which were used to evaluate *T*_m_.

**Figure 4:**
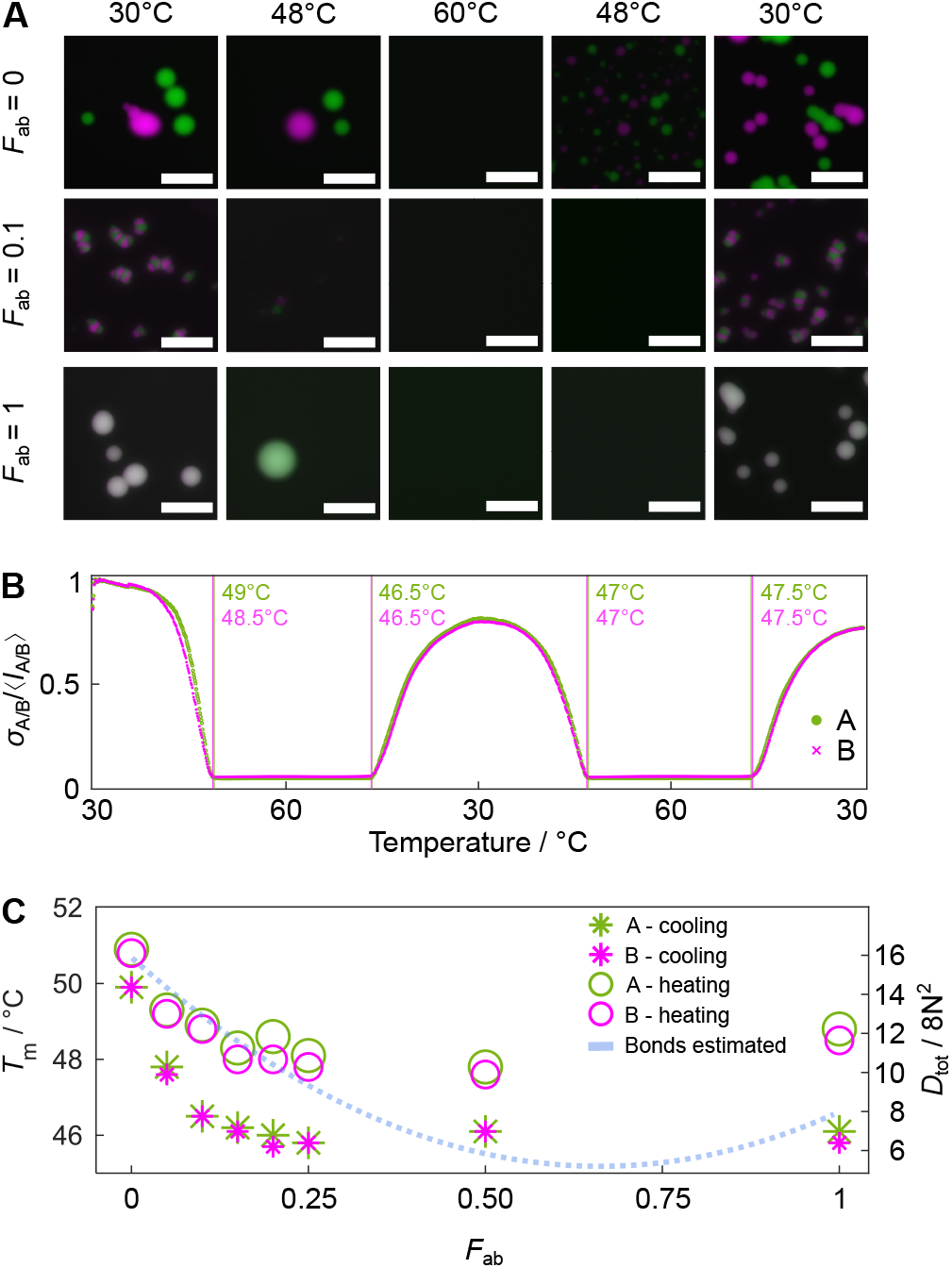
Condensate melting temperature depends on composition. (**A**) Selected epi-fluorescence snapshots of samples with *F*_ab_ = 0, 0.1, and 1 showing DNA condensates at specified temperatures on a heating ramp between 30°C - 60°C followed by a cooling ramp between 60°C - 30°C (± 0.5°C *per* 15 minutes). For each sample, all micrographs were scaled from 0 to maximum pixel intensity as identified in the first time point at 30°C. Scale bars, 50 µm. (**B**) The temperature at which condensates melt on heating and reform on cooling can be identified from sharp changes in *σ*_A*/*B_*/* ⟨*I*_A*/*B_⟩, where *σ*_A*/*B_ and ⟨*I*_A*/*B_⟩ are, respectively, the standard deviation and the mean of the pixel values evaluated from epifluorescence images of the ATTO 488 (NS A) and Alexa 647 (NS B) fluorescent channels within a field of view. The example shows data for *F*_ab_ = 0.10 over two heating and two cooling ramps. (**C**) Transition temperatures depend on *F*_ab_ and can be understood through an analysis of nanostar interaction combinatorics (Supplementary Note III). Hysteresis between cooling and heating ramps reflects the effects of slow melting or nucleation kinetics, or delays in heat propagation. Data are averages of at least six measurements performed on heating (circles) and cooling (stars) ramps. Standard deviation values of *T*_m_ are provided in Table S5. All samples have *F*_A_ = *F*_B_ = 0.5.

Data are presented in Figure 4**C** and Table S5 for symmetric samples with *F*_A_ = *F*_B_ and variable 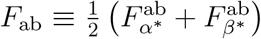. Expectedly, we observe a small degree of thermal hysteresis, with *T*_m_ values measured on heating ramps being slightly higher than those measured on cooling. This effect is likely the combined result of the finite timescales of condensate formation and dissolution and heat propagation across the sample cell. More interesting is the trend in *T*_m_ observed as a function of *F*_ab_, showing a maximum at *F*_ab_ = 0, followed by a decrease at intermediate *F*_ab_ values and a slight increase for *F*_ab_ = 1. We argue that this trend emerges from combinatorial effects dependent on linker composition. As discussed in Supplementary Note III (Supplementary Information), one can estimate the total number of bonds that can form between NSs as a function of *F*_ab_, under the simplified assumption that linkers act as mediators of NS-NS interactions, rather than particles in their own right. For a system with equal numbers of A and B NSs (*N*_A_ = *N*_B_ = *N*), and a stoichiometrically matched total number of linkers *N*_L_ = 4 *N*, the total number of bonds possible between nanostars A-A (*D*_A→A_), B-B (*D*_B→B_), and A-B (*D*_A⇌B_) can be computed with a simple combinatorial calculation of pairwise interactions (Supplementary Note III). The total numebr of bonds, *D*_tot_ can thus be expressed as a function of *F*_ab_, yielding

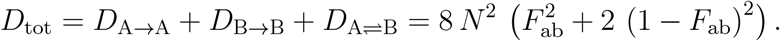

As shown in Figure 4**C**, this expression follows a trend remarkably similar to that observed for *T*_m_, with a minimum for *F*_ab_ ~ 2/3 and a maximum at *F*_ab_ = 0.

### 2.5 Phase Coexistence Depends on the Annealing Protocol

In an attempt to ensure thermodynamic equilibration, all experimental data presented so far were obtained from condensates annealed extremely slowly (− 0.1°C per hour) between 55°C - 25°C. The higher temperature extreme was chosen to be above the highest measured *T*_m_ (Figure 4), ensuring that the nanostructures were initially dispersed and that condensates could nucleate and relax over the slow cooling ramp. In general, the composition of co-existing phases can be temperature-dependent, implying that redistribution of nanostructures between co-existing domains may have to take place during the cooling transient, if thermodynamic equilibrium at 25°C is to be ultimately reached. The nucleation and growth behavior of the condensate droplets may also favour the emergence of metastable configurations. For instance, critical nuclei and small condensates may have compositions that differ from those of the bulk phases observed in large condensates, either due to kinetic or finite size effects, requiring substantial restructuring for the growing condensates to reach equilibrium.

It has been demonstrated that the molecular rearrangement timescales of DNA nanostar phases slow down exponentially upon cooling, according to an Arrhenius scaling [61]. Similarly, rough estimates from Fluorescence Recovery After Photobleaching (FRAP) experiments conducted at *T* ≥ 28°C [46, 62–64] place diffusion constants of NSs within condensates at ~ 10^−1^ to ~ 10^−2^ µm^2^ s^−1^ and relaxation times over the relevant lengthscales of 10-100 µm in the order of minutes to hours (Supplementary Note IV). Combined with the potential need for multicomponent condensates to undergo significant compositional changes, slow relaxation kinetics may make equilibration difficult to achieve. Furthermore, temperature-dependent relaxation kinetics and compositional constraints can impact mixtures differently depending on composition, given the dependence of *T*_m_ upon system stoichiometry (Figure 4).

To verify the occurrence of incomplete equilibration, we tested three different types of annealing protocols, assessing their impact on condensate morphology and phase composition. In a first set of protocols, samples were initially held at 55°C to ensure full melting, before performing a rapid quench to different incubation temperatures *T*_inc_=35°C, 37°C or 39°C, maintained constant for *t*_hold_ = 72 hours. A second protocol testing a much longer isothermal incubation – *t*_hold_ = 3 weeks – was also performed at *T*_inc_ =39°C. In the third protocol, samples were quenched from 55°C to 45°C, followed by a slow annealing ramp (− 0.1°C *per* 30 min) to 35°C. In all cases, after incubation at *T*_inc_ or after the annealing ramp, samples were cooled to 25°C at a cooling rate of −0.05°C min^−1^ [see Supplementary Information for full details on sample preparation and annealing protocols].

The confocal micrographs shown in **Figure** 5 enable comparison between condensate morphologies observed at room temperature, following each annealing protocol, for mixtures with symmetric composition (*F*_A_ = *F*_B_) but variable *F*_ab_. Samples at a given *F*_ab_ were prepared in sealed capillary tubes and sequentially subjected to the various annealing protocols, ensuring that observed effects are not caused by stoichiometric differences. Micrographs of samples discussed previously – annealed in well plates between 55°C and 25°C – are included for comparison.

**Figure 5:**
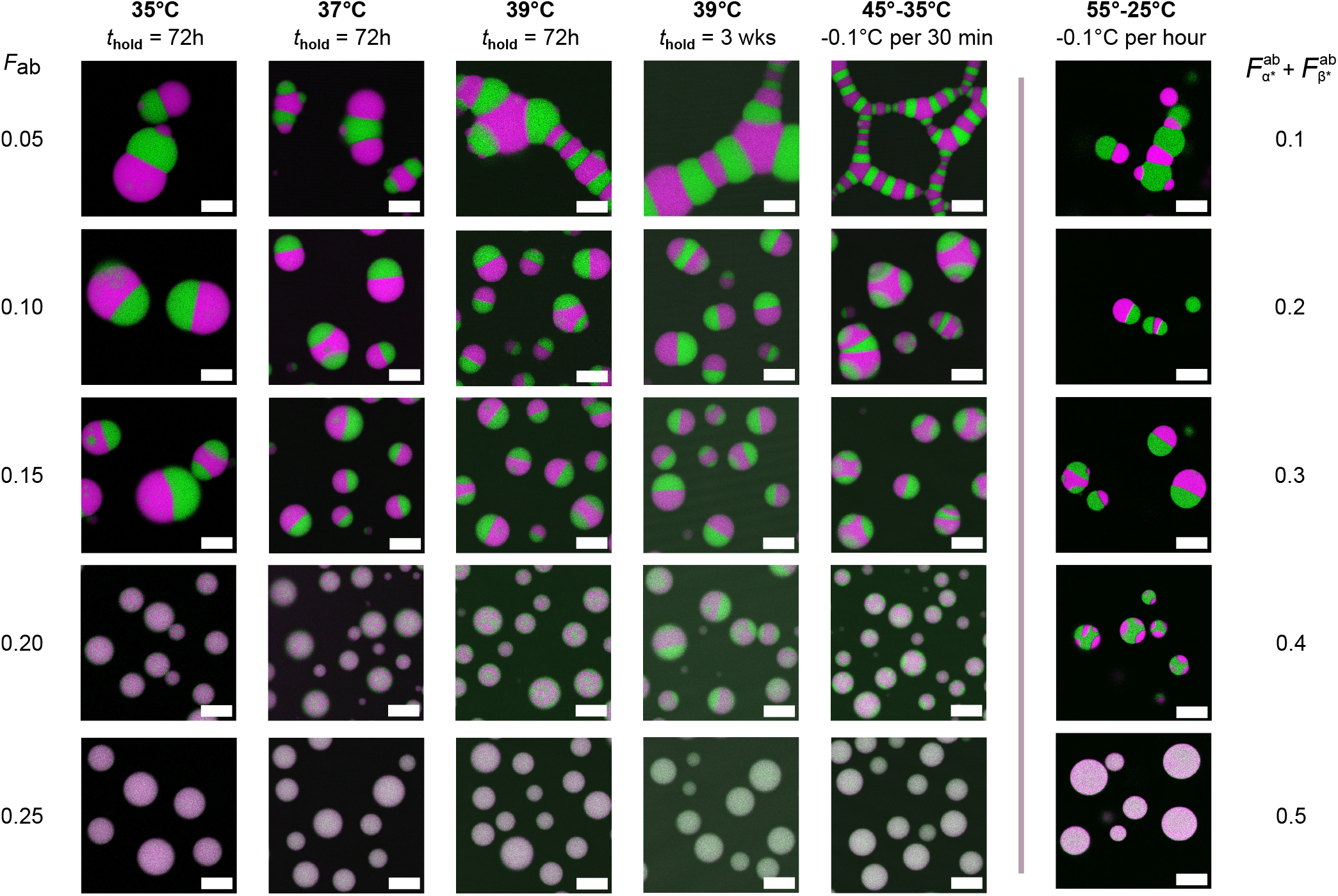
Phase coexistence in binary DNA condensates depends on thermal annealing protocol. As previously shown, mixing increases with increasing *F*_ab_, but the extent of mixing and and condensate morphology depend on the annealing conditions chosen, indicating significant non-equilibrium effects. Rows show images of samples with varying *F*_ab_, while columns correspond to different annealing protocols. For all tested conditions except those in the two right-most columns, samples were initially held at 55°C and then subject to a rapid quench and incubated at the temperature indicated for a time equal to *t*_hold_, before a further rapid cooling step to 25°C. For the second-to-right column, isothermal incubation was replaced by a cooling ramp between 45°C and 35°C. For comparison, the right-most column shows images from samples subject to a very slow anneal from 55°C to 25°C, from which data in Figure 2 and 3 were obtained. All samples have *F*_A_ = *F*_B_ = 0.5. All micrographs were scaled from 0 to maximum pixel intensity to aid visualization. Scale bars, 20 µm.

It is evident that the annealing protocol influences condensate size, morphology and the degree of de-mixing. For *F*_ab_ = 0.05, discrete condensates are observed for isothermal annealing at 35°C and 37°C, while extended branched networks with a distinct candy cane pattern are seen when incubating at 39°C or applying the 45°C - 35°C cooling ramp, similar to those observed by Jeon *et al*. [30]. The discrete two-phase condensates forming at *F*_ab_ = 0.1 and 0.15 are larger when samples are held at 35°C, but do not substantially differ in size across other annealing protocols. Most notably, the occurrence of macroscopic A-B demixing at *F*_ab_ = 0.2 depends on the annealing protocol. The very slow cooling ramp used in previous sections yields clear de-mixing, while near-uniform condensates are observed upon isothermal incubation at 35°C. Small, phase separated domains are visible after a 72-hour incubation at 39°C, which coarsen into faint macroscopic domains after 3 weeks. Blurred, small domains are seen with the 45°C - 35°C cooling ramp. In all cases, condensates are round, indicating vanishingly small interfacial tension between A- and B-rich domains.

We used image segmentation, as above, to assess the impact of the annealing protocol on the degree of interphase mixing through partition coefficients (**Figure** 6 and Figure S24) and interfacial tension ratios (Figure S25). The partition coefficients, *ρ*_A_ and *ρ*_B_, place the onset of de-mixing below 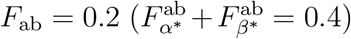 for all alternative annealing protocols, as the segmentation algorithm is unable to identify the small or nearly-mixed domains present under these conditions. At lower *F*_ab_, a trend emerges among isothermal incubation protocols, whereby the degree of mixing decreases with decreasing incubation temperature, *T*_inc_. The sample incubated at 35°C shows the least mixing, similar to samples prepared with the very slow annealing ramp discussed in previous sections. The three-week isothermal incubation at 39°C yields the most mixed condensates. These trends could be thermodynamically justified under the hypothesis that the A- and B-rich phases become more compositionally distinct at lower temperature, and the observed *ρ*_A_ and *ρ*_B_ reflect the equilibrium values at the respective incubation temperature. Upon rapid quenching from *T*_inc_ to 25°C, slow relaxation could then be preventing changes in domain composition. Because, as outlined in Table S8, SE affinity depends on temperature, one may argue that the morphological and compositional changes observed upon isothermally annealing at different temperatures reflect, at least partially, variations in binding strength. The differences observed between the 72 hour and 3 week incubation protocols at *T*_inc_ = 39°C, however, indicate that kinetics influences de-mixing, with domains progressively relaxing towards more mixed states. To rationalize this observation, we inspected image sequences collected *vs* temperature for 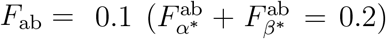 and *F*_A_ = *F*_B_, shown in Figure S23. We note that, upon cooling, condensates apparently containing only the A-rich or B-rich phase initially emerge, and then partially coalesce to form Janus-like droplets. It is plausible that the initial single-phase condensates have a composition different from equilibrium, meaning that reshuffling of the nanostructures may be required after coalescence events, explaining the slow evolution of *ρ*_A_ and *ρ*_B_ after incubation at 39°C. The 45°C - 35°C cooling ramp produces *ρ*_A_ and *ρ*_B_ trends as a function of *F*_ab_ that differ from other protocols, arguably as a result of *T*_m_ depending on sample composition (Figure 4).

**Figure 6:**
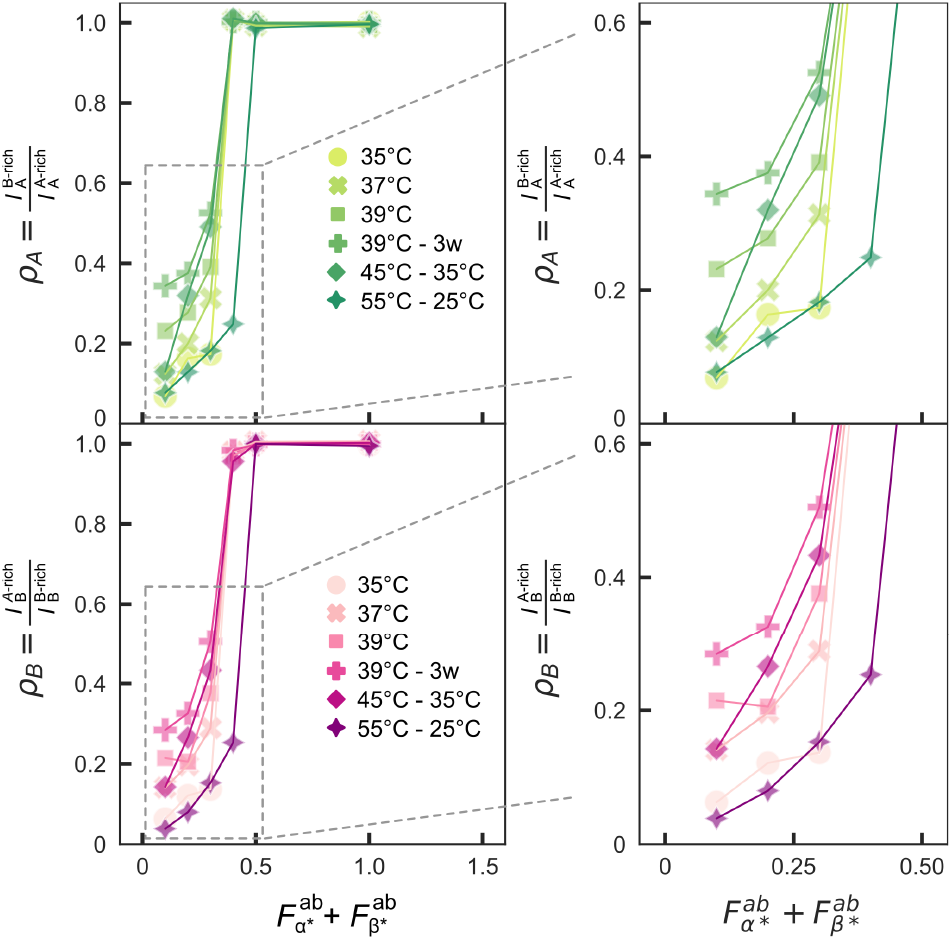
Interphase mixing in biphasic condensates depends on the annealing protocol. Partition coefficient data for the A-rich phase (*ρ*_A_, top) and B-rich phase (*ρ*_B_, bottom) are shown for all samples and annealing protocols discussed in Figure 5, as a function of the heuristic parameter 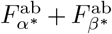. Zoomed-in insets (right) highlight trends in *ρ*_A_ and *ρ*_B_ as a function of annealing protocol. Datapoints represent the median of the condensate populations for a given sample composition, imaged in a field of view. The distributions of partition coefficients is provided in Figure S25 and the number of condensates analyzed for each composition and annealing condition are collated in Table S6.

The data in Figure 5 and 6 thus paint a complex picture in which the annealing protocol may influence the observed morphology and phase composition through both thermodynamic and kinetic factors. Thermodynamically, samples slowly annealed from 55°C to 25°C (close to room temperature) may be the closest to equilibrium at the end of the process. In turn, samples incubated isothermally before being quenched more rapidly may be metastable at room temperature but closer to equilibrium at their respective incubation temperature, although differences noted with changing incubation time indicate that equilibrium at the incubation temperature may not be reached. On the one hand, these findings highlight the need to carefully consider the possibility of incomplete equilibration when describing the behavior of nucleic acid liquids and condensates. On the other hand, similar factors are likely to influence the structure of isothermally assembling multiphase biological condensates, and may warrant detailed investigation. It is further worth noting that the complexities associated with coalescence and fusion in biphasic condensates, which are bound to be highly dependent on the exact morphology of individual multi-domain condensates, are likely to impact equilibration kinetics.

## 3 Conclusion

In summary, we present a biomimetic system of condensate-forming DNA nanostructures enabling the quantitative study of internal phase separation and transitions between single and two-phase condensates. The system consists of two non-interacting DNA nanostars – A and B – and three divalent linkers mediating A-A, B-B and A-B interactions. Nanostar-nanostar affinity can be continuously controlled through linker stoichiometry, generating an array of monophasic and biphasic condensates with programmable degree of mixing and morphology.

We identify an order parameter, 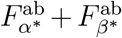, that depends uniquely on relative linker concentration and can describe the monophasic to biphasic transition regardless of the relative abundance of the two nanostar populations. 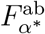 and 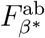 and can be intuitively interpreted as the frequency of A-B bonds, relative to the total number of bonds formed by A and B nanostars, respectively. To test the generality of our findings, we suggest that an analogous order parameter could be extracted for other relevant binary mixtures of condensate-forming macromolecules where intra- and inter-species affinities can be estimated. A similar exercise could help rationalize monophasic-to-biphasic transitions seen in complex cellular environments, *e*.*g*. those occurring in condensates of the tumor-suppressing transcription factor p53, observed to undergo internal phase separation in the presence of non-target nucleic acids while remaining uniform when interacting with p53-target DNA [65].

We show that the experimental phenomenology can be replicated by an effective three-component Flory-Huggins model, describing the two nanostar types and surrounding solvent. The Flory-Huggins interaction parameter modulating A-B interactions, χ_AB_, can be mapped onto 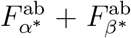, making the model able to quantitatively predict monophasic to biphasic transitions and morphological observables, including the degree of interphase mixing. The model is agnostic to the structural details of the DNA nanostructures including their size, valency and flexibility, and describes interaction-mediating linkers implicitly. While this lack of detail may be seen as a limitation, we argue that the model’s simplicity adds to its generality and speculate that analogous theoretical descriptions could be robust against design variations, including changes in NS valency or interaction schemes in which NS SEs bind directly rather than through the action of linkers. It is however expected that, in these scenarios, quantitative mapping between experimental and simulated phase behaviour would have to be recalibrated.

We further observe that changing the annealing protocol influences condensate morphology, highlighting the significance of slow coarsening kinetics on the structure and relative domain composition in multiphase condensates.

Our model is fully modular and enables exploration of the phase space by simply changing the stoichiometry of its constituents, without the need of introducing new nanostructure designs. The concept of modular condensate design, demonstrated here, could be extended to larger ensembles of nanostructures, expanding the available design space.

Thanks to this modularity and the associated predictive theory, our system represents a valuable tool to unravel the biophysics of internal phase separation in biomolecular condensates, known to play a prominent role in regulating their biological functions [66–68]. Furthermore, the design rules we uncovered will enable the design of synthetic condensates with tailored characteristics, highly sought after in biomanufacturing [69–71], therapeutics [72], synthetic biology [73], and synthetic cell engineering, whereby encapsulated multiphase droplets could function as synthetic organelles [33, 74, 75]. The well understood response of DNA-nanostar liquids to changes in ionic conditions [45] adds to the versatility of the platform, and can further be exploited to generate programmable responses and functionality [76], as done for other classes of liquid-liquid phase separated materials [77,78]. While our DNA-based system was studied *in vitro*, similar design principles can be extended to RNA-based condensates assembled from genetically encoded RNA nanostructures, which can be potentially expressed in living cells [37, 38].

## Author contributions

D.A.T. performed all experiments and data analysis, aided by L.M. and R.R.S. D.O. performed all analytical and numerical work, supported by E.F. L.D.M. designed and supervised the research. D.A.T., D.O. and L.D.M. wrote the paper. All authors contributed to result interpretation and presentation, and reviewed the manuscript.

## Acknowledgments

We thank the following for contributions to this work: L. Rovigatti (Sapienza University of Rome) and F. Sciortino (Sapienza University of Rome) for helpful discussions, G. Fabrini (The Francis Crick Institute) for the Python script used to extract melting temperatures from UV-Vis melting curves, E. J. Will (EPFL) for advice in the well-plate selection, the Facility for Imaging by Light Microscopy (FILM) at Imperial College London and Stephen Rothery for advice on confocal microscopy imaging.

## Funding

L.D.M., D.A.T. and L.M. acknowledge support from the European Research Council (ERC) under the Horizon 2020 research and innovation programme (ERC-STG No 851667 - NANOCELL). L.D.M. acknowledges support from a Royal Society University Research Fellowship (UF160152, URF\R\221009). R.R.S. acknowledges funding from the Biotechnology and Biological Sciences Research Council through a BBSRC Discovery Fellowship (BB/X010228/1). E.F. and D.O. acknowledge support from the U.S. National Science Foundation (NSF) grant FMRG: Bio award 2134772, by the Alfred Sloan Foundation through awards G-2021-16831 and G-2024-22575.

## Conflict of interest

The authors declare that they have no competing interests.

## Data and materials availability

All data are available from the corresponding author upon reasonable request. A permanent and freely accessible data repository will be generated upon acceptance for publication.

## Notes

### Competing Interest Statement

The authors have declared no competing interest.

### Summary of Updates

Improved discussion of results and moved some content from the SI to the main text.

