## Supporting Information for "Internal Phase Separation in Synthetic DNA Condensates"

### Experimental Section

#### Design of DNA Building Blocks

DNA nanostars (A and B) and linkers (aa, bb, ab) were designed using NUPACK [1] and their correct folding was analyzed using the NUPACK analysis module. All sequences, including the fluorescently-labeled versions, are provided in Table S1. Each nanostar (A or B) is composed of 4 dsDNA arms terminating in a 6 nt long 5' ssDNA overhang (or sticky end). Each linker (aa, bb, or ab) is composed of a dsDNA domain terminating in a 6 nt long 5' ssDNA sticky end complementary to those on nanostar arms.

#### Oligonucleotide Preparation

All oligonucleotides were purchased from Integrated DNA Technologies (IDT). Strands were purified by the suppliers. Non-functionalized strands were purified using a standard desalting procedure and fluorophore-functionalized strands were purified using high-performance liquid chromatography (HPLC).
DNA strands were received freeze-dried and were reconstituted in 1× TE buffer (10 mM Tris, 1 mM EDTA, pH 8.0, Sigma-Aldrich). All buffers were diluted using ultrapure water and syringe-filtered through 0.22 µm pore filters (Hydrophilic 33 mm polyethersulfone (PES) membrane). The concentration of reconstituted DNA strands was determined using the Beer-Lambert Law. Extinction coefficients were provided by the manufacturer and the absorbance of reconstituted strands was measured at 260 nm using a Thermo Scientific NanoDrop One UV-Vis spectrophotometer. Reconstituted strands were stored in the fridge at 4°C for short periods of time (< 2 weeks) or in the freezer at -20°C for longer periods. Before each use, reconstituted strands were vortexed and centrifuged.
Stocks of free nanostars (A and B) and linkers (aa, bb, ab) were prepared using the required oligonucleotides and diluted in 300 mM NaCl in 1× TE to give a final nanostar concentration of 2 µM and final linker concentration of 4 µM. The mixtures were annealed in 0.2 mL 8-tube strips using a thermal cycler (Bio-Rad C1000 Touch Thermal Cycler), by incubating at 95°C for 15 minutes, and then cooling from 90°C to 25°C at a cooling rate of -0.1°C $min^{-1}$. After annealing, stocks of pre-annealed nanostars and linkers were stored in the fridge at +4°C for up to 1 month.

#### UV-Vis Melting Curves

The melting temperatures of nanostars (A and B) and linkers (aa, bb, ab) were determined using UV-Vis melting curves and are listed in Table S2. All melting temperatures for the five building blocks are above 65°C, confirming that heating up mixtures of pre-annealed nanostars and linkers to 55°C for the annealing of condensates would not disrupt the already formed constructs.
UV-Vis melting curves were determined by monitoring absorbance at 260 nm as a function of temperature using an Agilent Cary 3500 UV-vis spectrophotometer. A volume of 150 µL DNA samples in 300 mM NaCl in 1× TE was loaded into low volume quartz cuvettes (HellmaAnalytics, 10 mm light path) and topped with 200 µL mineral oil (Sigma-Aldrich). DNA concentration was chosen such that absorbance values would be below 1. Nanostars concentration was kept at 300 nM (total strands concentration 1.2 µM) and linkers concentration 600 nM (total strands concentration 1.2 µM). Absorbance was monitored over a cooling ramp (90°C - 25°C) and a heating ramp (25°C - 90°C). The heating/cooling rate was set to ±0.4°C $min^{-1}$ and data was acquired every 0.1 °C. Nitrogen purging was set to a flow rate of 10 LPM to prevent condensation on the cuvettes upon cooling.
Melting temperatures ($T_m$) were extracted using a baseline approach [2] implemented in Python. The low-temperature plateau (55°C - 60°C) and the high temperature plateau (85°C - 90°C) of the sigmoid absorbance traces were fitted with straight lines. $T_m$ was then determined as the temperature at which the median between the two linear fits intersects the melting curve.

#### Agarose Gel Electrophoresis

The correct folding of newly designed nanostars and linkers was confirmed using agarose gel electrophoresis. Samples were annealed as described in *Oligonucleotide Preparation.*

Agarose gels were prepared at 1.5% (w/v) in 1× TBE buffer (89mM Tris, 89mM boric acid, 2mM EDTA, pH of 10× TBE = 8.3; Thermo Scientific) and stained with 0.01% (v/v) SYBR Safe DNA gel stain (Invitrogen). Gels were cast to a thickness of 5 mm and left to set for 90 minutes before covering with 1× TBE running buffer. Each well was loaded with a sample composed of ≈ 500 ng DNA and 1.3× BlueJuice Gel Loading Buffer (Invitrogen) diluted in 1× TBE to a final volume of 15 µL. Equivalent samples were made with a 100 bp DNA ladder (Invitrogen). Gels were run at a voltage of 7 V $cm^{-1}$ for 90 minutes and the electrophoresis cell was kept in an ice bath. Gels were imaged using a Syngene G:Box chemi XX6 system.

#### Condensates Annealed in 384-Well Plates

Condensates used for the study of the phase diagram (Figures 1**b**, 2 and 3) were annealed in 384-well plates (IBIDI, µ-Plate 384 Well Glass Bottom # 1.5 Coverslip, sterilized), enabling a high throughput screening of 63 different compositions in triplicate.

The 63 different compositions were made using pre-annealed stocks of nanostars (2 µM) and linkers (4 µM) and mixed in the defined ratios of nanostars A/B such that the final nanostars and linkers concentrations were 0.5 µM and 1 µM, respectively. 80 µL of each sample were prepared with an Opentrons OT-2 pipetting robot in 96-well plates (NEST 0.1 mL 96-Well PCR Plate, Full Skirt), mixed thoroughly, and finally loaded in the flat 384-well plates using 20 µL per well, three wells per sample. The well plate was then covered with adhesive aluminium foil (Thermo Scientific Adhesive PCR Plate Foils, stable between -40°C to + 120°C) and annealed on the heating block of a thermal cycler (Bio-Rad C1000 Touch Thermal Cycler). To improve thermal contact, the well plate was placed on a custom-made 2 mm thick aluminum sheet and covered with an ethylene propylene diene monomer (EPDM) rubber seal (Opentrons Thermocycler GEN2 Seals) sandwiched in between two aluminium foils. The annealing protocol included an incubation step at 55°C for 30 minutes to ensure the 6-nt long sticky ends were melted while preserving hybridization in nanostars and linkers, followed by a slow cooling ramp from 55°C to 25°C at a rate of -0.1°C $h^{-1}$. Samples were stored at 4°C until imaging.

#### Condensates Annealed in Glass Capillaries

Condensates used to study the effect of annealing protocols (Figures 5 and 6) were annealed in glass capillaries. Pre-annealed stocks of nanostars (2 µM) and linkers (4 µM) were mixed such that the final nanostars and linkers concentrations were 1 µM and 2 µM, respectively. A volume of 70 µL was then loaded into borosilicate glass capillaries (inner dimensions: 0.4 mm × 4 mm × 50 mm, CM Scientific) with a micropipette. Prior to loading, capillaries were cleaned by sonication in 1% Hellmanex III (HellmaAnalytics) at 40°C for 30 minutes. The surfactant was removed through a minimum of five rounds of rinsing with deionized (DI) water, followed by a round of sonication in ultrapure water (Milli-Q), rinsing with ethanol, and finally drying with nitrogen. After loading, the capillary ends were capped with mineral oil (Sigma Aldrich). Capillaries were placed on a cover slip (24 mm × 60 mm, Menzel-Gläser) and sealed with epoxy glue (Araldite) and left to dry for a minimum of 2 hours. To improve thermal contact with the heating block of thermal cycler (Bio-Rad C1000 Touch Thermal Cycler), the coverslips with sealed capillary tubes were wrapped in aluminium foil.

The annealing protocol included an incubation step at 55°C for 30 minutes to ensure melting of the 6 nt sticky ends without disrupting nanostars and linkers, followed by a direct quench to the holding temperature (35°C, 37°C or 39°C) where the samples were left to equilibrate for 72 hours. In one protocol, samples were left to equilibrate at 39°C for 3 weeks. In another protocol, the sample were cooled down slowly between 45°C - 35 °C at a cooling rate of -0.01°C $min^{-1}$. In all protocols, the samples were finally cooled down to 25°C at a cooling rate of -0.05°C $min^{-1}$.

#### Imaging Methods

**Confocal Laser Scanning Microscopy for Sample Characterization.**

Confocal images were acquired with a Leica TCS SP5 microscope using a 20×/0.50 NA HC PL FLUOTAR dry objective (Leica) for samples annealed in capillaries (Figures 5 and 6) and a 40×/0.85 NA HCX PLAN APO dry objective (Leica) for samples annealed in 384-well plates (Figures 1, 2, 3). Samples were imaged either directly in the 384-well plates with # 1.5 coverslip glass bottom in which they were annealed, or in the sealed glass capillaries in which they were annealed.

For ATTO 488, a 488 Argon ion laser was used and emission was measured between 493 and 543 nm. For Alexa 647, the He/Ne 633 laser was used and emission was measured between 638 nm and 698 nm. Images were acquired in a frame-sequential mode. The pinhole was set to 1 AU. Images were acquired with a scan speed of 400 Hz, in a 2048 × 2048 format (pixel size 387.5 nm × 387.5 nm) for the phase diagram data and 4096 × 4096 format (pixel size 189.3 nm × 189.3 nm) for the data set in which annealing was varied (apart from the 39°C $F_{ab}$ = 0.05 and 0.10 data sets which are in a 2048 × 2048 format).

**Epifluorescence Microscopy for the Characterization of Melting and Annealing of DNA Condensates.**

Timelapses of the melting and formation of DNA condensates were performed using a Nikon Eclipse Ti2-E inverted microscope with a Perfect Focusing System (PFS) equipped with a Plan Apo $\lambda$ 20×/0.75 NA, WD 1000 µm dry objective (Nikon), a Lumencor SPECTRA X LED engine and a Hamamatsu Orca-Flash4.0v3 camera.
To control the temperature while imaging, a custom-made microscopy stage was used that enabled samples in glass capillary tubes to be mounted on a Peltier element (Temikra) and imaged in real-time.

#### Segmentation and Image Analysis

Custom MATLAB[3] scripts were written to enable the segmentation and image analysis of the condensates. All segmentation and analysis were performed on raw grayscale 8-bit images exported in .tif format. All confocal data were acquired in 8-bit format. Epifluorescence data were acquired in 16-bit format and converted to 8-bit format for analysis. Analysis to determine partition coefficients and contact angles was performed on a condensate-by-condensate basis. Specific MATLAB functions are provided in brackets.

##### Identification and Isolation of Condensates

To identify each condensates as a distinct domain, the entire merged field of view (FOV) was binarized (IMBINARIZE). Domains below a size threshold and on the edge (*i.e.* not fully in the FOV) were excluded. An example of condensates selected for further analysis is shown in Figure S18**a**. Then, each domain was found by identifying boundaries (BWBOUNDARIES) and labeled (LABEL2RGB) as regions of interest (ROIs). A bounding box (STATS.BOUNDINGBOX) and the centroid coordinates (STATS.CENTROID) were used to identify the region to be cropped as an enlarged bounding box with double the width and height of that of the condensate. Then, each channel (BF, ATTO488, and Alexa 647) was cropped using the coordinates of the enlarged bounding box.

##### Mask Identification for each Fluorescent Channel

For each FOV cropped around a condensate and each of the two fluorescent channels (ATTO 488, Alexa 647), a Gaussian filter (IMGAUSSFILT) was applied to reduce pixel level noise and to help with the identification of the condensate boundaries. Gaussian filters are linear filters that smooth images by averaging pixel values with a Gaussian kernel. Then, a flat structuring element was defined (STREL) to perform a morphological opening (IMOPEN) operation that can help with removing small domains and smooth the image. Finally, images with enhanced contrast where 1% of the data is saturated at low and high intensities were obtained by adjusting (IMADJUST) the result obtained by subtracting the opened image from the Gaussian filtered image. To identify a mask in each of the two channels, the corresponding flattened image was used. A thresholding level was found for each channel (MULTITHRESH) and then used to binarize the image (IMQUANTIZE). An example of the masks identified along with key image segmentation steps is shown in Figure SS15.
Finally, the correlation between the two masks was calculated (CORR2) to assess whether samples are mixed ($\geq 0.91$) or de-mixed ($< 0.91$). For mixed samples, the masks described above were used. For de-mixed samples, the intersection of the two masks was subtracted from each mask. Three examples are shown in Figure S17 for various degrees of mixing and mask overlap.
It is important to note the masks identified through image processing were applied to raw images to extract quantitative measurements.

##### Calculation of Partition Coefficients

For each of the two fluorescent channels, we calculate the two partition coefficients, $\rho_A$ and $\rho_B$, as defined in Equation S1 and Equation S2, respectively:

$$\rho_A = \frac{I_A^{\text{B-rich}}}{I_A^{\text{A-rich}}} \tag{S1}$$

$$\rho_B = \frac{I_B^{\text{A-rich}}}{I_B^{\text{B-rich}}} \tag{S2}$$

The masks for A-rich and B-rich domains were identified as described above and applied to the raw images of each channel. For example, the partition coefficient $\rho_A$ (corresponding to the A-rich phase labeled with ATTO488), is calculated as the ratio between the ATTO 488 fluorescence intensity in the B-rich domain and the ATTO 488 fluorescence intensity in the A-rich domain. The value of fluorescence intensity $I_A^{\text{B-rich}}$ is determined by the sum of pixel values in the ATTO 488 channel within the B-rich mask and divided by the number of pixels in the mask. Similarly, $I_A^{\text{A-rich}}$ is determined by the sum of pixel values in the ATTO 488 channel within the A-rich mask divided by the number of pixels in the mask. For samples prepared to study the effects of annealing conditions (Figure 5), we also subtract background signal from both

the numerator and denominator in the partition coefficient calculation to ensure the increasing background in samples re-annealed multiple times does not affect the measurements. The background signal intensity is determined as the median pixel value in the background mask of the entire FOV. In samples prepared for the phase diagram (Figures 1, 2, 3), the background contribution was close to 0 and not subtracted.
It should be noted that for de-mixed samples where the 2D-correlation coefficient of the two masks is < 0.91 (CORR2), the fluorescent contribution in the area belonging to the two masks was removed to avoid including regions assigned in both the A-rich and B-rich domains as exemplified in Figure S17.

##### Calculation of Contact Angles

A custom function was written to identify contact angles in each de-mixed condensate. The key steps are highlighted for representative condensates with one interface in Figure S15 or multiple interfaces in Figure S16.
Based on the masks identified in each channel, we find the boundaries (BWBOUNDARIES) for each segmented domain (*e.g.*, for a Janus-like droplet there would be two domains and one interface as shown in Figure S15; for a network, there would be $n$ domains and $n - 1$ interfaces as shown in Figure S16). Masks are determined as described above, and used to extract contours of the three relevant interfaces (A-water, B-water and A-B). We then fit the contours to circles using the Pratt method [4] with a modification that enforces the circle fitted to the interface to pass through the two points of intersection of the circles fitted to each of the neighbouring phases.
At each interface, the three circles fitted to the A-rich, B-rich, and interface intersect in two points. At each intersection point, tangents to these circles are plotted and used to determine contact angles as highlighted in Figure 3**A**. Contact angles are then reported as the average of the two angles identified at each intersection point.
Finally, before saving the contact angle measurement, the user was prompted to choose whether to keep or discard the measurement based on how well the circles were fitted to the two phases and the interface. In Figure S17 the examples in panels **a** and **b** were kept, while the example in **c** was discarded. In the discarded condensates, the two phases were not well de-mixed, leading the two initial masks (before subtracting the intersection of the masks) to almost coincide. As a result, the circles fitted to the two domains almost coincide and lead to an incorrect contact angle measurement. To prevent bias, the value of the contact angle was not shown and the decision was made only based on the fitting of the circles and tangents. Figure S18 shows an example of condensates selected for partition coefficients measurements in panel **a** and the sub-set of condensates selected for contact angle measurements in panel **b**.
FIJI (ImageJ)[5, 6] was used for cropping and scaling of confocal micrographs (where applicable and specified in figure captions).

#### Transition Temperature Determination through Epifluorescence Microscopy

The eight samples with $F_{\mathrm{A}} = F_{\mathrm{B}} = 0.5$ and varying $F_{\mathrm{ab}}$ highlighted in Figure 4 were mounted on a single coverslip and attached to the Peltier element such that all different compositions were subjected to the same heating and cooling cycles. The samples were made from pre-annealed stocks as previously described with a total NS concentration of 1 µm and linker concentration of 2 µm.
Samples, mounted next to each other on the Peltier element, were exposed to two consecutive heating and cooling cycles from 30°C to 60°C and back to 30°C for a total of four ramps, at a rate of ±0.5°C every 15 minutes.
Temperature was changed in steps of 0.5°C and values of transition temperatures extracted for two cooling and heating ramps were averaged.
At each time point, the standard deviation of all pixel values within a FOV was calculated and normalized by the mean pixel value of the FOV and the maximum value, ensuring that the y-axis is scaled from 0 to 1. Normalized standard deviation values were plotted against time points (and corresponding temperatures). The transition temperature was found by identifying abrupt changes in the variance of the normalized standard deviation values.

#### Statistical Analysis

Below is a summary of the statistical analysis approach used. Additional details are provided in the relevant methods sections, figure captions, or table captions.

1. **Pre-processing of data**
   All analyses on micrographs were performed using raw files. For visualization purposes, some micrographs were contrast-enhanced by scaling intensity values. In such cases, the specific scaling method is described in the corresponding figure captions.

2. **Data presentation**
   Given the diversity of sample types and experimental conditions, median values are typically presented in the main text, with full distributions shown as box plots in the Supplementary Information.

3. **Sample sizes**
   The number of measurements for each analysis is reported in the Supplementary Tables (Tables S3, S4, S6, S7) or in figure captions.

4. **Software used for analysis**
   Image segmentation and agarose gel analysis were performed in MATLAB[3] using custom scripts. UV-Vis spectroscopy data were analyzed using custom scripts written in Python. Box plots were plotted in Python.

### Supplementary Note I: Flory-Huggins Modeling

In the following, we will describe the Flory-Huggins model, from both a theoretical and a numerical perspective. To begin, we will discuss the issues associated with fields where two components are very attractive towards one another. Subsequently, we will introduce a six-component theory, describing all nanostars and linkers. Empirical results suggest the development of a two-component theory to describe the combined linker-nanostar model used in the main text. Finally, we shall present numerical details for the solution of the final model using a numerical scheme.

##### Attractive fluid

For illustrative purpose, we will begin with the Flory-Huggins theory[7] applied to a ternary system and describing the interaction of two components, A and B, immersed in a solvent. For now, we ignore contributions to the free energy arising from surfaces. The free energy of such a system is given by:

$$F = \int \mathrm{d}\mathbf{x} f(\mathbf{x}) \tag{S3}$$

$$\begin{aligned} f(\mathbf{x}) = &\phi_{\mathrm{A}}(\mathbf{x})\log(\phi_{\mathrm{A}}(\mathbf{x})) + \phi_{\mathrm{B}}(\mathbf{x})\log(\phi_{\mathrm{B}}(\mathbf{x})) + (1-\phi_{\mathrm{A}}(\mathbf{x})-\phi_{\mathrm{B}}(\mathbf{x}))\log(1-\phi_{\mathrm{A}}(\mathbf{x})-\phi_{\mathrm{B}}(\mathbf{x})) \\ &+ \chi_{\mathrm{AB}}\phi_{\mathrm{A}}(\mathbf{x})\phi_{\mathrm{B}}(\mathbf{x}) \end{aligned} \tag{S4}$$

where $f$ is a free energy density, $\phi_i(\mathbf{x})$ is the packing fraction of component $i$, the interaction between components A and B is calculated via the parameter $\chi_{\mathrm{AB}}$ and we have used the incompressibility condition on the combination of A, B, and the solvent.

The free energy in Equation S4 corresponds to a theory in two components, with the solvent implicitly included through the incompressibility condition. The relations of phase separation theory[7] can be used to determine the presence of coexisting phases, by calculating the chemical potentials and osmotic pressures *i.e.*, the chemical potentials and osmotic pressures must be matched between the different phases. The chemical potentials and osmotic pressures are given by, respectively:

$$\mu_i = \nu_i \frac{\partial f}{\partial \phi_i} \tag{S5}$$

$$\Pi = -f + \boldsymbol{\phi} \cdot \nabla f \tag{S6}$$

where $\nu_i$ is the volume of component $i$ (assumed to be equal to unity from now on), and $\boldsymbol{\phi}$ is a vector containing all the packing fractions in the problem. The composition of differing phases, I and II, can be found through the solutions of equations:

$$\mu_i^{\mathrm{I}}\big|_{\phi_{\mathrm{A}}^{\mathrm{I}},\phi_{\mathrm{B}}^{\mathrm{I}}} = \mu_i^{\mathrm{II}}\big|_{\phi_{\mathrm{A}}^{\mathrm{II}},\phi_{\mathrm{B}}^{\mathrm{II}}} \tag{S7}$$

$$\Pi^{\mathrm{I}}\big|_{\phi_{\mathrm{A}}^{\mathrm{I}},\phi_{\mathrm{B}}^{\mathrm{I}}} = \Pi^{\mathrm{II}}\big|_{\phi_{\mathrm{A}}^{\mathrm{II}},\phi_{\mathrm{B}}^{\mathrm{II}}} \tag{S8}$$

Intuitively, one would expect that in the limit that $\chi_{\mathrm{AB}}$ (the coupling between A and B) becomes very large and negative, and that the abundances of A and B are similar, the free energy outlined in Equation S4 need not be described using two components anymore, but rather using a one-component theory. We may thus try to understand to what extent this intuition is true, depending on the value of $\chi_{\mathrm{AB}}$, and what would be a new order parameter that can be used to describe the simplified model. One path forward is to compute the phase diagram of a full two-component theory (which is not too difficult in this case), and compare it with that of a standard one-component model to determine the validity of the approximation.

The phase boundaries for the two component theory, shown in Figure SN1**a** and **b**, look similar to a standard one-component phase separating system, with some differences depending on whether there is an equal amount of $\phi_{\mathrm{A}}$ and $\phi_{\mathrm{B}}$ or not (panels **a** and **b**, respectively). Of perhaps more interest is whether the analysis of the compositions of each phase gives rise to a new order parameter for describing this system as an effective one-component system. One may either reason as to its form or find it from the compositions. In either case, one finds that the following parameter is roughly equal in both phases:

$$\psi = \frac{(\phi_{\mathrm{A}} - \langle\phi_{\mathrm{A}}\rangle) - (\phi_{\mathrm{B}} - \langle\phi_{\mathrm{B}}\rangle)}{(\phi_{\mathrm{A}} - \langle\phi_{\mathrm{A}}\rangle) + (\phi_{\mathrm{B}} - \langle\phi_{\mathrm{B}}\rangle)} \tag{S9}$$

This expression is somewhat more involved than what one might have simply posited, which would be the difference in the packing fractions over their total, but it accounts for the cases where we do not have equal amounts of A and B. The value of $\psi$ (calculated numerically) in two different phases is shown in Figure SN1**c**, showing that the two phases have the same value of $\psi$. The two new fields are defined as:

$$\rho = (\phi_{\mathrm{A}} - \langle\phi_{\mathrm{A}}\rangle) + (\phi_{\mathrm{B}} - \langle\phi_{\mathrm{B}}\rangle) \tag{S10}$$

$$\delta = (\phi_{\mathrm{A}} - \langle\phi_{\mathrm{A}}\rangle) - (\phi_{\mathrm{B}} - \langle\phi_{\mathrm{B}}\rangle) \tag{S11}$$

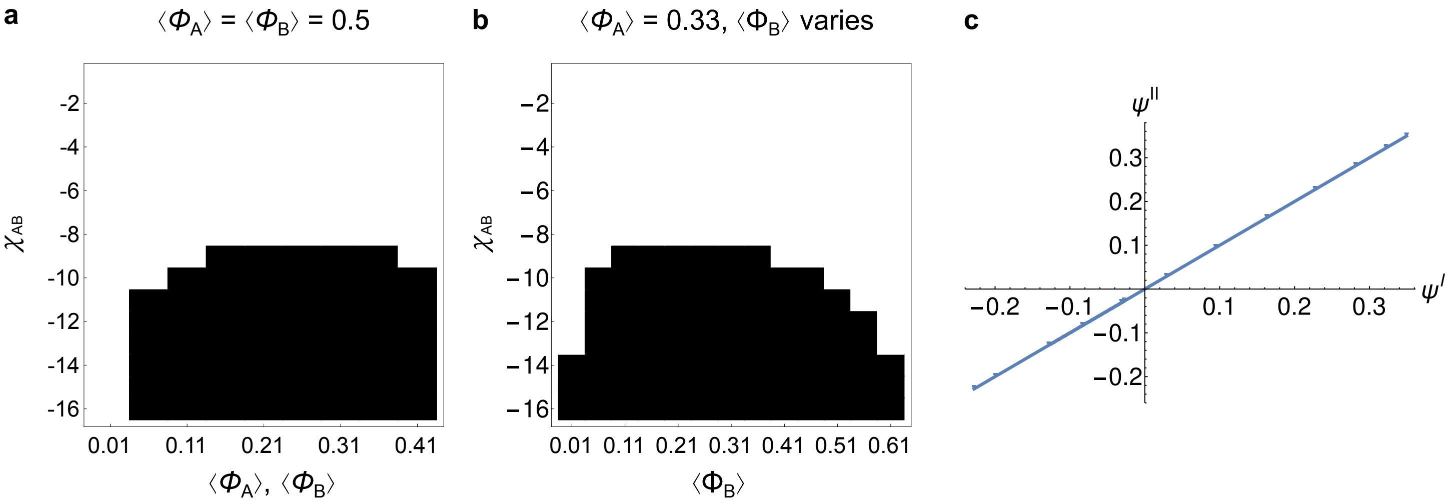


Figure SN1: Phase behavior of the two-component system with attractive interactions (calculated by solving Equation S7 and S8). (**a**) Phase diagram (black = phase separation) for the two component attractive system as a function of composition and interaction $\chi_{\mathrm{AB}}$, where the compositions of the A-rich phase and the B-rich phase are equal to one another. Phase separation can occur when attractive interactions exist between A and B nanostars. (**b**) Similar to panel **a**, but the total amount of A is held fixed, and the amount of B is varied. (**c**) Numerical calculation of the order parameter $\psi$ calculated in both the dense $\psi^{I}$ and dilute $\psi^{II}$ phases for the phase separated phases in **a** and **b**. The straight line shows that this parameter takes the same value in the two phases, indicating that it is a constant.

where it can be seen that $\delta = \psi\rho$. We can perform an expansion in $\psi$ of the original free energy in terms of these fields, leaving:

$$\begin{aligned} f = &\frac{1}{4}\psi^2\left(\frac{\rho^2}{\rho+2\langle\phi_{\mathrm{A}}\rangle} + \frac{\rho^3(-\chi) - 2\rho^2\chi\langle\phi_{\mathrm{B}}\rangle + \rho^2}{\rho+2\langle\phi_{\mathrm{B}}\rangle}\right) \\ &+ \frac{1}{2}\psi\left(-\rho\chi\langle\phi_{\mathrm{A}}\rangle + \rho\log(\rho+2\langle\phi_{\mathrm{A}}\rangle) + \rho\chi\langle\phi_{\mathrm{B}}\rangle - \rho\log(\rho+2\langle\phi_{\mathrm{B}}\rangle)\right) \\ &+ \frac{1}{4}\Big((\rho+2\langle\phi_{\mathrm{B}}\rangle)\left(\rho\chi + 2\log(\rho+2\langle\phi_{\mathrm{B}}\rangle) + 2\chi\langle\phi_{\mathrm{A}}\rangle - 2\log(2)\right) \\ &- 4(\rho+\langle\phi_{\mathrm{A}}\rangle+\langle\phi_{\mathrm{B}}\rangle-1)\log(1-\rho-\langle\phi_{\mathrm{A}}\rangle-\langle\phi_{\mathrm{B}}\rangle) \\ &+ 2(\rho+2\langle\phi_{\mathrm{A}}\rangle)\left(\log(\rho+2\langle\phi_{\mathrm{A}}\rangle) - \log(2)\right)\Big) \end{aligned} \tag{S12}$$

It can be shown that when $\langle\phi_{\mathrm{A}}\rangle = \langle\phi_{\mathrm{B}}\rangle$, the term linear in $\psi$ cancels out. Assuming that $\psi$ is small, we can obtain the following expression for the free energy:

$$f = \frac{1}{4}\theta(\theta\chi - 4\log(2)) + \theta\log(\theta) + (1-\theta)\log(1-\theta) \tag{S13}$$

where $\theta = \rho + 2\langle\phi_{\mathrm{A}}\rangle$. Comparing to the standard phase separation free energy $f = \theta\log(\theta) + (1-\theta)\log(1-\theta) + \chi\theta(1-\theta)$, we can see that the two component field will yield identical results in this case to the one component field, with a rescaled $\chi \to -\chi/4$ (the linear terms in $\theta$ contributing to the free energy will not change the results).

#### Six Component Theory

Having observed the process by which one can reduce the number of components in a problem, we proceed to propose a function for the full six-component theory corresponding to nanostars, linkers, and the solvent solvent implicitly included through the incompressibility condition. We propose the following phenomenological bulk free energy density for the system consisting of nanostars and linkers:

$$\begin{aligned} f = &\phi_{\mathrm{bb}}\phi_{\mathrm{B}}\chi_{\mathrm{Bbb}} + \phi_{\mathrm{aa}}\phi_{\mathrm{A}}\chi_{\mathrm{Aaa}} + \chi_{\mathrm{ABab}}\phi_{\mathrm{A}}\phi_{\mathrm{B}}\phi_{\mathrm{ab}} \\ &+ (-\phi_{\mathrm{aa}} - \phi_{\mathrm{A}} - \phi_{\mathrm{bb}} - \phi_{\mathrm{B}} - \phi_{\mathrm{ab}} + 1)\log(-\phi_{\mathrm{aa}} - \phi_{\mathrm{A}} - \phi_{\mathrm{bb}} - \phi_{\mathrm{B}} - \phi_{\mathrm{ab}} + 1) \\ &+ \phi_{\mathrm{A}}\log(\phi_{\mathrm{A}}) + \phi_{\mathrm{bb}}\log(\phi_{\mathrm{bb}}) + \phi_{\mathrm{aa}}\log(\phi_{\mathrm{aa}}) + \phi_{\mathrm{B}}\log(\phi_{\mathrm{B}}) + \phi_{\mathrm{ab}}\log(\phi_{\mathrm{ab}}) \end{aligned} \tag{S14}$$

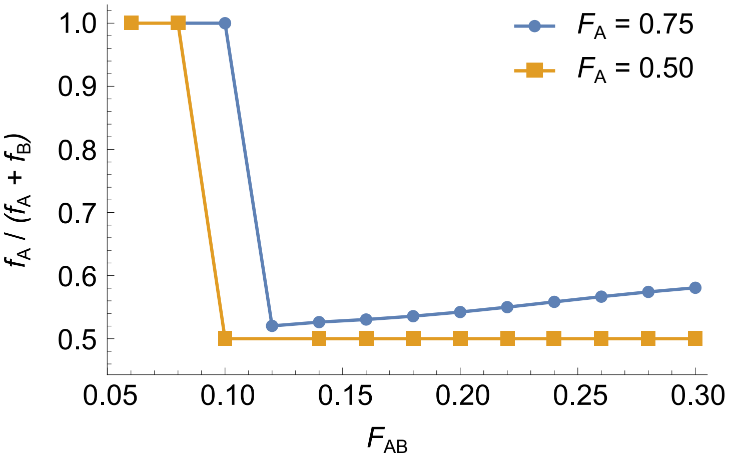


Figure SN2: Fraction of nanostars A inside the dense phase as a function of the ab linker fraction $F_{\mathrm{ab}}$ for different fractions of nanostars A ($F_{\mathrm{A}}$) and nanostars B ($F_{\mathrm{B}}$). While there are similarities to the experimental results, some qualitative differences remain. Simulations parameters in the six-component model are $\chi_{\mathrm{A}\alpha} = -20, \chi_{\mathrm{B}\beta} = -20, \chi_{\mathrm{AB}\gamma} = -200$, and $\langle \phi_{\mathrm{A}} \rangle = \langle \phi_{\mathrm{B}} \rangle = 0.02$. Note these are different from the parameters used for snapshots highlighted in the main text which were generated with the three-component model.

One can see from the expression above that the theory becomes somewhat more unwieldy when each identifiable component is introduced. In the expression of the free energy, the subscripts A and B refer to nanostars, the subscripts aa and bb correspond to the linkers that can either bind to A or B nanostars, and the subscript ab refers to linkers that can bind both A to B nanostars. The entropic contributions to the free energy arise naturally from the Flory-Huggins approach, but we have slightly more complicated interaction terms than usual. The interaction of aa with A (or bb with B) is captured through a simple binary interaction. However, when we consider linking between A and B, we model this through a ternary interaction. The reason for this choice over others is the ab linker can only link A to B. So, for example, binary interactions between A and ab linker would promote large agglomerations of A and ab linker, which would not occur in the experiment in the absence of B. The presence of ternary interactions makes this theory somewhat more involved than usual Flory-Huggins approaches, which usually truncate after binary interactions, though is not unprecedented[8]. Additionally, since this is a mean field model, factors such as the stoichiometry of A and B nanostars binding through the action of linkers are ignored. Therefore, the composition of phases might display some different features compared to the ones observed in the experiment. Nevertheless, the model presented in Equation S14 can suffice as a simple theory that includes the relevant variables in the problem.

We have three energy scales in this problem, $\chi_{\mathrm{A}\alpha}$, $\chi_{\mathrm{B}\beta}$ and $\chi_{\mathrm{AB}\gamma}$. The first two correspond to the interaction of the nanostars with their accompanying linkers, and the last one corresponds to the interaction between A, B and an ab linker. We omit a detail in this model, namely the differing sizes of the different components in the system. This could be theoretically introduced, though would double the effective parameter space of the problem, making drawing conclusions more difficult.

Evaluation of Equation S14 can be performed along similar lines to the previous section (finding phases with the same chemical pressure and osmotic pressures). These solutions can be found numerically. We do this as an example to find if there are any ways we can treat the additional variables in our problem (building on the previous section). This also allows us to probe whether the transitions observed in the main text can be observed in the model (Figure SN2).

As might have been anticipated, the free energy model included in Equation S14 produces a transition from a dispersed phase to a dense phase. The dense phase can either be constituted of separate A or B rich droplets, or, for a high amount of linker, one dense AB-rich droplet. This model does display qualitative differences to the experimental results which we rationalize based on the fact that it is challenging to achieve a relaxed system. In contrast to the experimental model, it is observed that the transition to mixed droplets occurs for smaller $F_{\mathrm{ab}}$ fractions of the ab linker in symmetric systems when $F_{\mathrm{A}} = F_{\mathrm{B}} = 0.5$ rather than asymmetric systems $F_A \neq 0.5$. Similarly, as this is a mean field model, the stoichiometry of the phases is not balanced in the mixed phase when $F_{\mathrm{A}} \neq F_{\mathrm{B}}$.

Similarly to before, in the case that the *ab* linker is present, the following quantity is roughly constant (found empirically):

$$\omega := \frac{(\phi_\gamma - \langle \phi_\gamma \rangle) - (\phi_{\mathrm{A}} - \langle \phi_{\mathrm{A}} \rangle) - (\phi_{\mathrm{B}} - \langle \phi_{\mathrm{B}} \rangle)}{(\phi_\gamma - \langle \phi_\gamma \rangle) + (\phi_{\mathrm{A}} - \langle \phi_{\mathrm{A}} \rangle) + (\phi_{\mathrm{B}} - \langle \phi_{\mathrm{B}} \rangle)} \tag{S15}$$

**Effective Model**

The observations made in previous sections can be used to constrain the form of an effective model that dispenses with many of the details we have developed for the six-field model. The model presented in Equation S14 can be extended to account for the presence of linkers that allow phase separation from the solvent, and which can be accounted for with the presence of additional terms, asserting a free energy density of the form:

$$\begin{aligned} f = {} & \phi_A \log(\phi_A) + \phi_B \log(\phi_B) + (1 - \phi_A - \phi_B)\log(1 - \phi_A - \phi_B) \\ & + \chi_{AB}\phi_A\phi_B + \chi_{AA}\phi_A(1 - \phi_A - \phi_B) \\ & + \chi_{BB}\phi_B(1 - \phi_A - \phi_B) \end{aligned} \tag{S16}$$

A Taylor expansion of Equation S14 where $\psi$ and $\omega$ are constants allows us to express the full six-component theory in terms of the theory given in Equation S16. This allows us to write down the effective $\chi$s in terms of parameters of the more comprehensive model:

$$\begin{aligned} \chi_{AA} = {} & \frac{(\psi_A + 1)\left((\omega - 1)(\psi_A - 1)\chi_{A\alpha} + (\omega + 1)\langle\phi_B\rangle(\psi_A + 1)\chi_{AB\gamma}\right)}{4(\omega - 1)} \\ \chi_{AB} = {} & \frac{1}{4}\left(\left(\psi_A^2 - 1\right)\chi_{A\alpha} + \left(\psi_B^2 - 1\right)\chi_{B\beta}\right) \\ & + \frac{1}{4((\omega - 1))}\chi_{AB\gamma}\Big( - \psi_B\left(\psi_A(\omega(\langle\phi_A\rangle + \langle\phi_B\rangle - \langle\phi_\gamma\rangle) + \langle\phi_A\rangle + \langle\phi_B\rangle + \langle\phi_\gamma\rangle\right) \\ & -\omega(\langle\phi_A\rangle - \langle\phi_B\rangle + \langle\phi_\gamma\rangle) - \langle\phi_A\rangle + \langle\phi_B\rangle + \langle\phi_\gamma\rangle) \\ & + \psi_A\left((\omega + 1)\langle\phi_B\rangle\psi_A - (\omega + 1)\langle\phi_A\rangle\right. \\ & +\omega(\langle\phi_B\rangle + \langle\phi_\gamma\rangle) + \langle\phi_B\rangle - \langle\phi_\gamma\rangle) + (\omega + 1)\langle\phi_A\rangle\psi_B^2 + (\omega - 1)\langle\phi_\gamma\rangle\Big) \\ \chi_{BB} = {} & \frac{(\psi_B + 1)\left((\omega + 1)\langle\phi_A\rangle\chi_{AB\gamma}(\psi_B + 1) + (\omega - 1)(\psi_B - 1)\chi_{B\beta}\right)}{4(\omega - 1)} \end{aligned} \tag{S17}$$

The expressions above are rather complex and involve a large number of constants. Moreover, some of the constants can only be determined through the solution of the field model. Matching to experiment is thus not of great utility, as the abundance of parameters makes conclusion tenuous. Nevertheless, we can ask whether the properties of the effective $\chi$ in the models are related to properties such as the linker concentration $\langle\phi_\gamma\rangle$.
The general trend for $\chi$s as a function of $\langle\phi_\gamma\rangle$ is given by the approximate formulas:

$$\chi_{AA}(\langle\phi_\gamma\rangle) \approx d_0 \tag{S18}$$

$$\chi_{BB}(\langle\phi_\gamma\rangle) \approx d_1 \tag{S19}$$

$$\chi_{AB}(\langle\phi_\gamma\rangle) \approx c_0 - c_1\langle\phi_\gamma\rangle \tag{S20}$$

where $c_i, d_i$ are constants. One can observe that with increasing ab linker concentration, the effective $\chi$ between the AB components decreases, as one would expect.
Additional complexity is induced by the fact the six-component model contains a sharp transition, as can be observed in Figure SN2. This means that the effective $\chi_{AB}$ could contain a sharp discontinuity as linker concentration ratios are varied (due to the fact that the different equilibrium phases may have different values of $\psi_A$,$\psi_B$ and $\omega$). Thus we expect the following qualitative picture to hold for the effective $\chi_{AB}$ as a function of linker concentration, shown in Figure SN3. Unfortunately, the transition in effective theories as a function of linker concentrations can only be determined from the full six-field model, given the empirical relationships underlying $\psi_A, \psi_B$ and $\omega$.

**Numerical Details**

In order to generate density profiles that can be compared to the results of experiment, we take the free energy model of the previous section with additional terms corresponding to the free energy of the surfaces:

$$\begin{aligned} f = {} & \phi_A \log(\phi_A) + \phi_B \log(\phi_B) + (1 - \phi_A - \phi_B)\log(1 - \phi_A - \phi_B) + \chi_{AB}\phi_A\phi_B \\ & + \chi_{AA}\phi_A(1 - \phi_A - \phi_B) + \chi_{BB}\phi_B(1 - \phi_A - \phi_B) + \frac{1}{2}\lambda^2\chi_{AA}|\nabla\phi_A|^2 + \frac{1}{2}\lambda^2\chi_{BB}|\nabla\phi_B|^2 \end{aligned} \tag{S21}$$

where we ignore the free energy contributions due to the surface between concentrations of A and B. The parameter $\lambda$ defines the interface width between A and B and the solvents. We set this parameter to be equal to 1 in our simulations.

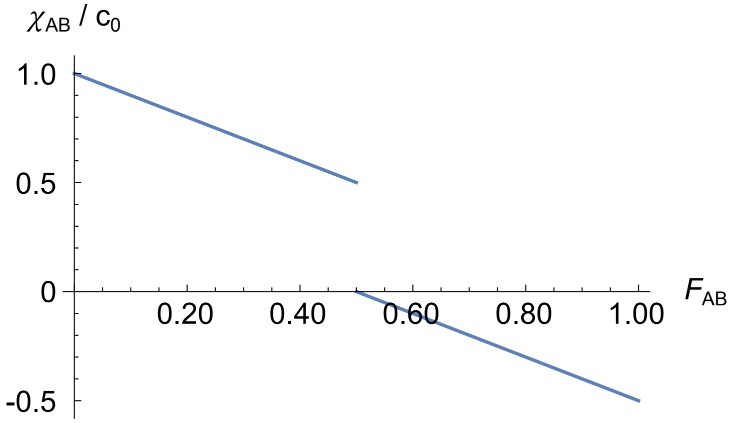


Figure SN3: Qualitative picture of the behavior of $\chi_{\mathrm{AB}}$ as a function of ab linker fraction $F_{\mathrm{ab}}$. The transition point occurs at a point which can only be determined from the full six-field theory.

The time evolution of this field theory can be performed using the Cahn-Hilliard equation[9].

$$\frac{\partial \phi_{\mathrm{A}}}{\partial t} = D_{\mathrm{A}} \nabla^2 \left( \frac{\delta f}{\delta \phi_{\mathrm{A}}} \right) \tag{S22}$$

$$\frac{\partial \phi_{\mathrm{B}}}{\partial t} = D_{\mathrm{B}} \nabla^2 \left( \frac{\delta f}{\delta \phi_{\mathrm{B}}} \right) \tag{S23}$$

where we have introduced the diffusion coefficients $D_{\mathrm{A}}, D_{\mathrm{B}}$ which we set equal to unity for both A and B. Numerically, the free energy $f$ is impacted due to logarithmic divergences when the concentrations of A and B are close to zero. For numerical computations we therefore do a Taylor expansion of free energy $f$ to fourth order around a point $a, b$ in the packing fractions of A and B. This can lead to slight differences in the results as the form of the free energy is slightly modified. The extent to which differences are important can be understood through the difference in the free energy surface as a function of $\phi_{\mathrm{A}}, \phi_{\mathrm{B}}$. We choose $(a, b) = (1/3, 1/3)$ such that this difference is minimized. The equations are solved numerically in Fourier space [see [10] for details].

#### Supplementary Note II: Derivations of Equation (5) and (6)

Recalling the definitions from the main text, we have

$$
\begin{aligned}
F_{\mathrm{A}} &= \frac{[\mathrm{A}]}{[\mathrm{NS}]} \\
F_{\mathrm{B}} &= \frac{[\mathrm{B}]}{[\mathrm{NS}]} \\
F_{\mathrm{ab}} &= \frac{[\mathrm{ab}]}{[\mathrm{aa}]+[\mathrm{bb}]+[\mathrm{ab}]} = \frac{[\mathrm{ab}]}{[\mathrm{linker}]} \\
F_{\mathrm{aa}} &= \frac{[\mathrm{aa}]}{[\mathrm{aa}]+[\mathrm{bb}]+[\mathrm{ab}]} = \frac{[\mathrm{aa}]}{[\mathrm{linker}]} \\
F_{\mathrm{bb}} &= \frac{[\mathrm{bb}]}{[\mathrm{aa}]+[\mathrm{bb}]+[\mathrm{ab}]} = \frac{[\mathrm{bb}]}{[\mathrm{linker}]},
\end{aligned}
$$

where $F_{\mathrm{ab}}$ is the fraction of the ab linker, $F_{\mathrm{aa}}$ the fraction of the aa linker, and $F_{\mathrm{bb}}$ the fraction of the bb linker.
As discussed in the main text, in a stoichiometric regime [linker] : [NS] = 2 : 1 or [linker] = 2[NS]. Recalling the constrain in terms of concentrations of individual components in Equation 2 and Equation 3, we have:

$$
\begin{aligned}
4[\mathrm{A}] &= 2[\mathrm{aa}] + [\mathrm{ab}] \\
4[\mathrm{B}] &= 2[\mathrm{bb}] + [\mathrm{ab}]
\end{aligned}
$$

which can be rearranged to:

$$[\mathrm{aa}] = 2[\mathrm{A}] - \frac{[\mathrm{ab}]}{2} \tag{S24}$$

$$[\mathrm{bb}] = 2[\mathrm{B}] - \frac{[\mathrm{ab}]}{2} \tag{S25}$$

Dividing Equation S24 by [linker] leads to the alternative expression for $F_{\mathrm{aa}}$ (Equation 5):

$$
\begin{aligned}
\Longrightarrow\ \frac{[\mathrm{aa}]}{[\mathrm{linker}]} &= \left(2[\mathrm{A}] - \frac{[\mathrm{ab}]}{2}\right) \times \frac{1}{[\mathrm{linker}]} \\
F_{\mathrm{aa}} &= \frac{2[\mathrm{A}]}{[\mathrm{linker}]} - \frac{[\mathrm{ab}]}{2[\mathrm{linker}]} \\
F_{\mathrm{aa}} &= \frac{2[\mathrm{A}]}{2[\mathrm{NS}]} - \frac{F_{\mathrm{ab}}}{2} \\
F_{\mathrm{aa}} &= \frac{2[\mathrm{A}]}{2[\mathrm{NS}]} - \frac{F_{\mathrm{ab}}}{2} \\
F_{\mathrm{aa}} &= F_{\mathrm{A}} - \frac{F_{\mathrm{ab}}}{2}
\end{aligned}
$$

In the same way, dividing Equation S25 by [linker] leads to the alternative expression for $F_{\mathrm{bb}}$ (Equation 6):

$$
\begin{aligned}
F_{\mathrm{bb}} &= F_{\mathrm{B}} - \frac{F_{\mathrm{ab}}}{2} \\
F_{\mathrm{bb}} &= (1 - F_{\mathrm{A}}) - \frac{F_{\mathrm{ab}}}{2}
\end{aligned}
$$

### Supplementary Note III:
### Nanostar Interaction Combinatorics

Let us consider a sample with equal number of A and B NSs, $N_{\mathrm{A}} = N_{\mathrm{B}} = N$, and a stoichiometrically matched number of linkers $N_{\mathrm{L}} = 4N$. We then allow the fraction of ab linkers, $F_{\mathrm{ab}}$ to vary.
We consider the simplified picture in which linkers act as mediators of NS-NS interactions, rather than describing them as particles in their own right. In this scenario, we note that there are $4N$ $\alpha$ SEs on A-type NSs. Of these $N^{\mathrm{SE}}_{\mathrm{A}\to\mathrm{A}}$ will be, on average, hybridized to aa linkers, and thus mediating A-A interactions, while $N^{\mathrm{SE}}_{\mathrm{A}\to\mathrm{B}}$ will be hybridized to ab linkers, and thus mediating A-B interactions. These quantities can be expressed as

$$N^{\mathrm{SE}}_{\mathrm{A}\to\mathrm{A}} = 4N\frac{2N_{\mathrm{aa}}}{2N_{\mathrm{aa}}+N_{\mathrm{ab}}} = 4N\frac{2N_{\mathrm{aa}}}{N_{\mathrm{aa}}+N_{\mathrm{bb}}+N_{\mathrm{ab}}} = 8N\,F_{\mathrm{aa}} = 4N\,(1-F_{\mathrm{ab}}) \tag{S26}$$

$$N^{\mathrm{SE}}_{\mathrm{A}\to\mathrm{B}} = 4N\frac{N_{\mathrm{ab}}}{2N_{\mathrm{aa}}+N_{\mathrm{ab}}} = 4N\frac{N_{\mathrm{ab}}}{N_{\mathrm{aa}}+N_{\mathrm{bb}}+N_{\mathrm{ab}}} = 4N\,F_{\mathrm{ab}}, \tag{S27}$$

where $N_{\mathrm{aa}}$, $N_{\mathrm{ab}}$, and $N_{\mathrm{bb}}$ are the numbers of aa, ab, and bb linkers, respectively, such that $N_{\mathrm{L}} = N_{\mathrm{aa}}+N_{\mathrm{bb}}+N_{\mathrm{ab}}$, and where we used $N_{\mathrm{aa}} = N_{\mathrm{bb}}$, valid for this symmetric system with $N_{\mathrm{A}} = N_{\mathrm{B}}$. Similarly, for the B-type NSs, we can express

$$N^{\mathrm{SE}}_{\mathrm{B}\to\mathrm{B}} = N^{\mathrm{SE}}_{\mathrm{A}\to\mathrm{A}} = 4N\,(1-F_{\mathrm{ab}}) \tag{S28}$$

$$N^{\mathrm{SE}}_{\mathrm{B}\to\mathrm{A}} = = N^{\mathrm{SE}}_{\mathrm{A}\to\mathrm{B}} = 4N\,F_{\mathrm{ab}}. \tag{S29}$$

We can therefore describe the population of SEs as 4 sets of particles, two sets consisting of $N^{\mathrm{SE}}_{\mathrm{A}\to\mathrm{A}} = N^{\mathrm{SE}}_{\mathrm{B}\to\mathrm{B}}$ particles that can interact only within the same set, and two sets consisting of $N^{\mathrm{SE}}_{\mathrm{A}\to\mathrm{B}} = N^{\mathrm{SE}}_{\mathrm{B}\to\mathrm{A}}$ particles that can interact only with each other. We can thus compute the total number of bonds possible for each set as

$$D_{\mathrm{A}\to\mathrm{A}} = D_{\mathrm{B}\to\mathrm{B}} = \frac{N^{\mathrm{SE}}_{\mathrm{A}\to\mathrm{A}}\left(N^{\mathrm{SE}}_{\mathrm{A}\to\mathrm{A}}-1\right)}{2} \approx \frac{{N^{\mathrm{SE}}_{\mathrm{A}\to\mathrm{A}}}^{2}}{2} = 8N^2\,(1-F_{\mathrm{ab}})^2 \tag{S30}$$

$$D_{\mathrm{A}\rightleftharpoons\mathrm{B}} = \frac{N^{\mathrm{SE}}_{\mathrm{A}\to\mathrm{B}}\,N^{\mathrm{SE}}_{\mathrm{A}\to\mathrm{B}}}{2} = 8N^2\,F^2_{\mathrm{ab}}, \tag{S31}$$

where we used $N_{\mathrm{A}\to\mathrm{A}} \gg 1$. The total number of possible bonds in the system is thus

$$D_{\mathrm{tot}} = D_{\mathrm{A}\to\mathrm{A}} + D_{\mathrm{B}\to\mathrm{B}} + D_{\mathrm{A}\rightleftharpoons\mathrm{B}} = 8N^2\left(F^2_{\mathrm{ab}} + 2\,(1-F_{\mathrm{ab}})^2\right). \tag{S32}$$

Figure SN4 shows that Equation S32 is maximized at $F_{\mathrm{ab}} = 1$ and has a minimum at $F_{\mathrm{ab}} = 2/3$, consistent with the trends in $T_{\mathrm{m}}$ measured in Figure 4.

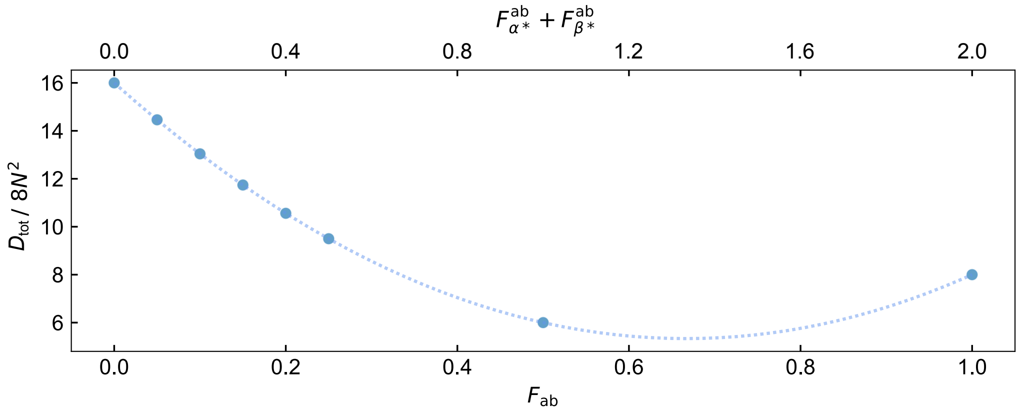


Figure SN4: **Estimated total number of bonds $D_{\mathrm{tot}}$ in a system of $N$ nanostars as a function of $F_{\mathrm{ab}}$ when $F_{\mathrm{A}} = F_{\mathrm{B}} = 0.5$, as derivied in Equation S32.** Markers correspond to values of $F_{\mathrm{ab}}$ tested experimentally. The trend in the number of combinations resembles the trend in transition temperatures measured experimentally, and shown in Figure 4**C**.

### Supplementary Note IV:
### Estimation of Diffusion Constants

To assess whether kinetic effects observed in our system could arise from diffusion-limited equilibration within condensed phases, we estimated characteristic diffusion constants ($D$) using published Fluorescence Recovery After Photobleaching (FRAP) data on related DNA nanostar (NS) systems.
These rough estimates provide insight into the timescales for molecular rearrangements and equilibration in NS-based condensates.
Diffusion constants were approximated using the standard relation:

$$D \approx \frac{w^2}{4\tau},$$

where $w$ is the bleaching radius and $\tau$ is the characteristic recovery time. The following estimates were derived from the literature:

- **Sato et al.** (2020) [11]: $D \approx 1.3 \times 10^{-14}\,\mathrm{m^2 s^{-1}}$ ($T = 60\,^\circ\mathrm{C}$, liquid-like; $w \approx 5\,\mu\mathrm{m}$; $\tau \approx 8\,\mathrm{min}$; $5\,\mu\mathrm{M}$; 8-nt sticky end, 75% GC; trivalent NS; Note: no recovery at $30\,^\circ\mathrm{C}$, gel-like;)
- **Lee et al.** (2021) [12]: $D \approx 4.2 \times 10^{-14}\,\mathrm{m^2 s^{-1}}$ ($T = 30\,^\circ\mathrm{C}$, liquid-like; $w \approx 10\,\mu\mathrm{m}$; $\tau \approx 10\,\mathrm{min}$; $20\,\mu\mathrm{M}$; 6-nt sticky end, 67% GC; trivalent NS)
- **Kengmana et al.** (2024) [13]: $D \approx 2.1 \times 10^{-13}\,\mathrm{m^2 s^{-1}}$ ($T = 37\,^\circ\mathrm{C}$, liquid-like; $w \approx 1\,\mu\mathrm{m}$; $\tau \approx 2\,\mathrm{min}$; $5\,\mu\mathrm{M}$; 6-nt sticky end, 67% GC; trivalent NS)
- **Skipper and Wickham** (2024) [14]: $D \approx 8.3 \times 10^{-14}\,\mathrm{m^2 s^{-1}}$ ($T = 28\,^\circ\mathrm{C}$, liquid-like; $w \approx 1\,\mu\mathrm{m}$; $\tau \approx 5\,\mathrm{min}$; $10\,\mu\mathrm{M}$; 6-nt sticky end, 67% GC; trivalent NS)

#### Supplementary Figures

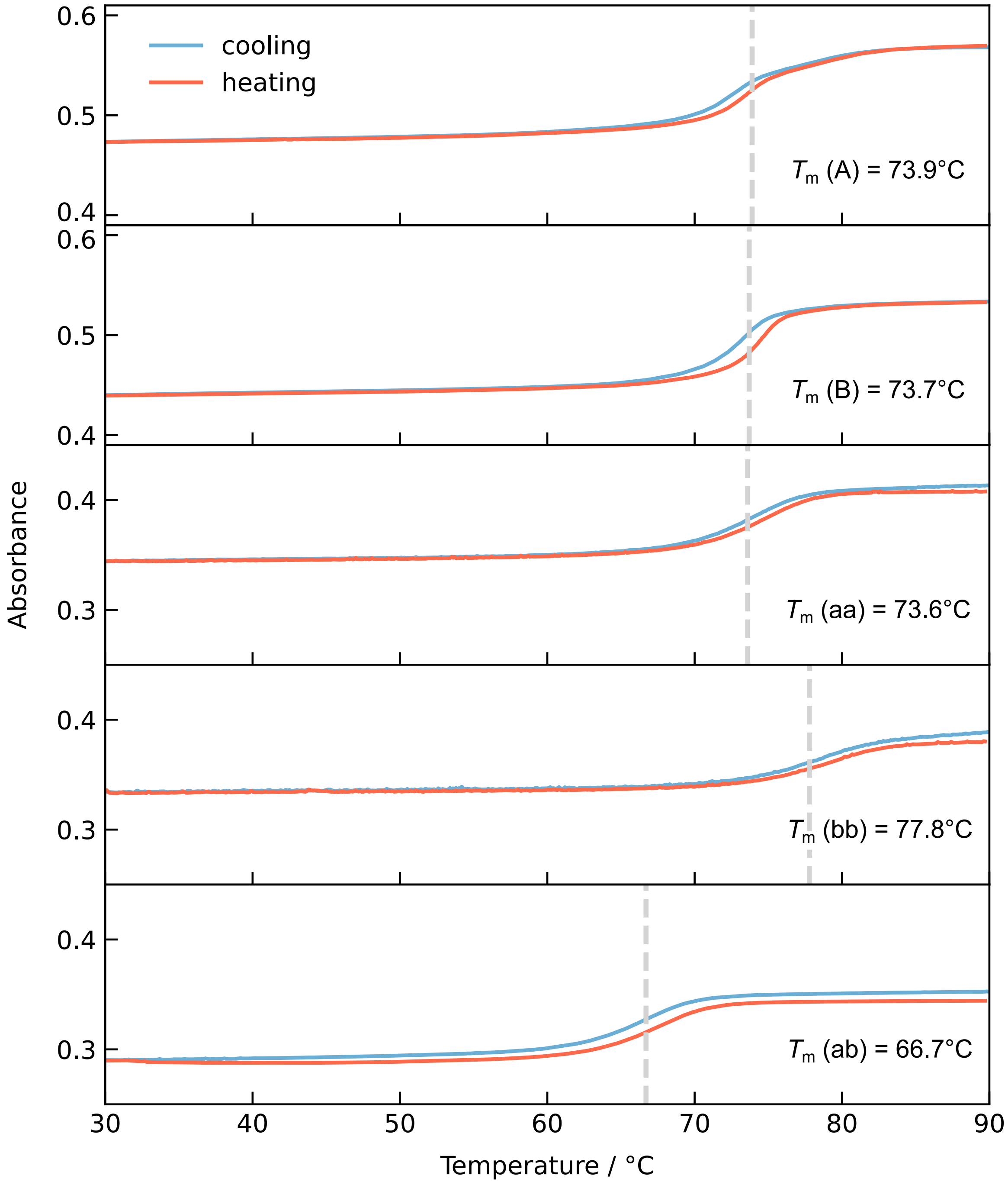


Figure S1: **Representative UV-Vis melting curves on cooling (blue) and heating (red) ramps for the five building blocks.** The grey dotted line shows the average $T_m$ as reported in Table S2.

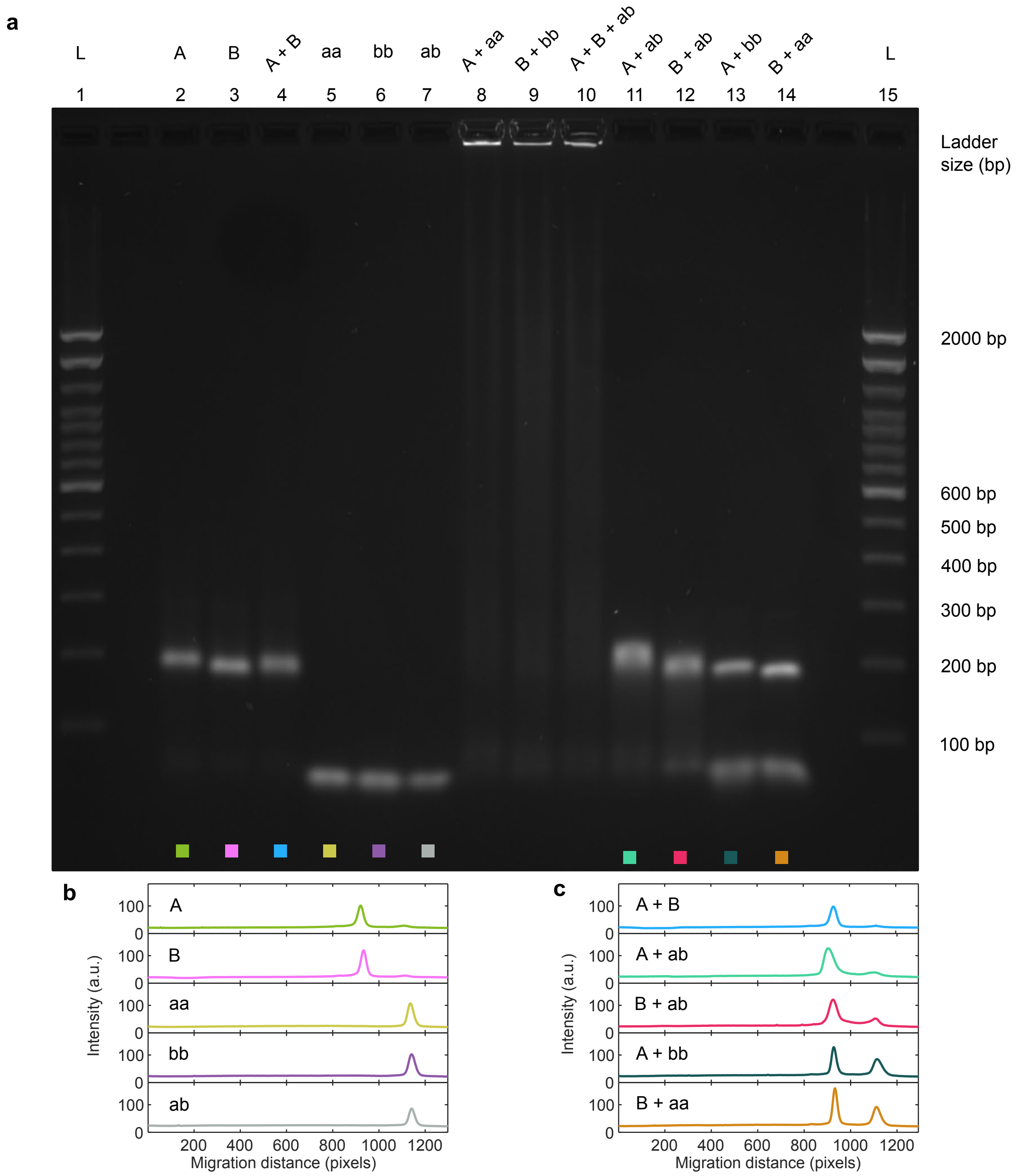


Figure S2: **Agarose gel electrophoresis confirms correct folding of nanostars (NS) and linkers, and interactions between constructs.** (**a**) Image of the agarose gel with a 100 bp ladder (L) on each side. Samples in lanes 8, 9, and 10 correspond to stoichiometric mixtures (not annealed) of NS and complementary linkers. The lack of migration from the wells in lanes 8, 9, and 10 confirms the intended binding between A/aa, B/bb, and A/B/ab. Lanes 11 and 12 show interactions between the ab linker and NS A and NS B, respectively. Lanes 13 and 14 show the lack of interactions between NS and orthogonal linkers. The faint band at a large migration distance present in all lanes containing nanostars is likely to correspond to excess core-forming strands or partially formed nanostars. (**b**) Lane intensity profiles for NS A and B (lanes 2 and 3) and linkers (lanes 5, 6, 7). (**c**) Lane intensity profiles for combinations (not annealed) of NS and linkers.

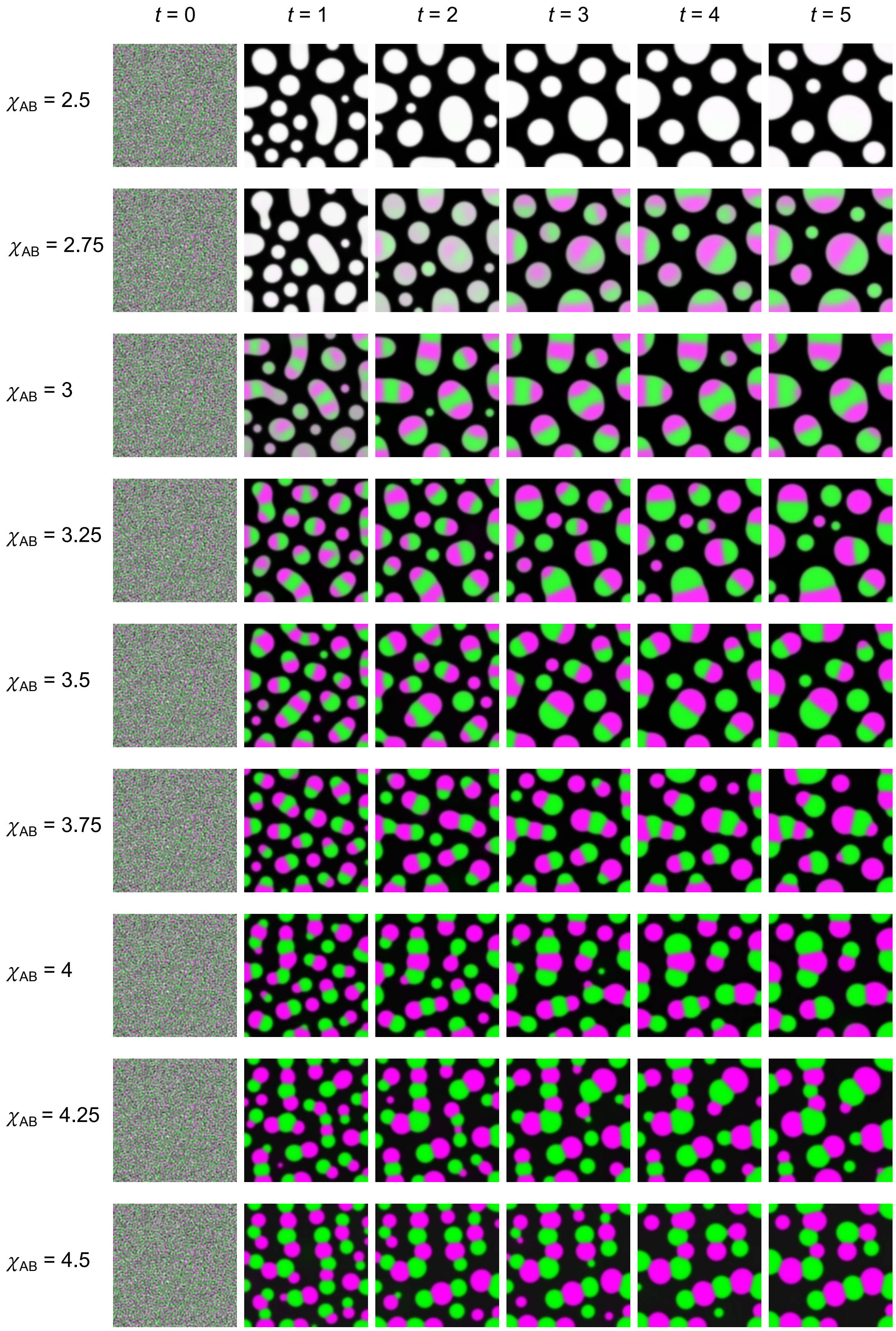


Figure S3: **Simulations snapshots**, corresponding to five different time points and nine different $\chi_{\mathrm{AB}}$ values producing fully mixed condensates (top) and partially mixed constructs with various degrees of de-mixing. In all cases, we kept $\chi_{\mathrm{AA}} = \chi_{\mathrm{BB}} = 4$ and the concentrations of the two components were kept constant and in a ratio of 1:1. The packing fractions $\phi_{\mathrm{A}}$ and $\phi_{\mathrm{B}}$ were set to 0.23. The residual interfaces at $\chi_{\mathrm{AB}} = 4.5$, forming despite the unfavorable A-B interactions, are likely a result of incomplete equilibration.

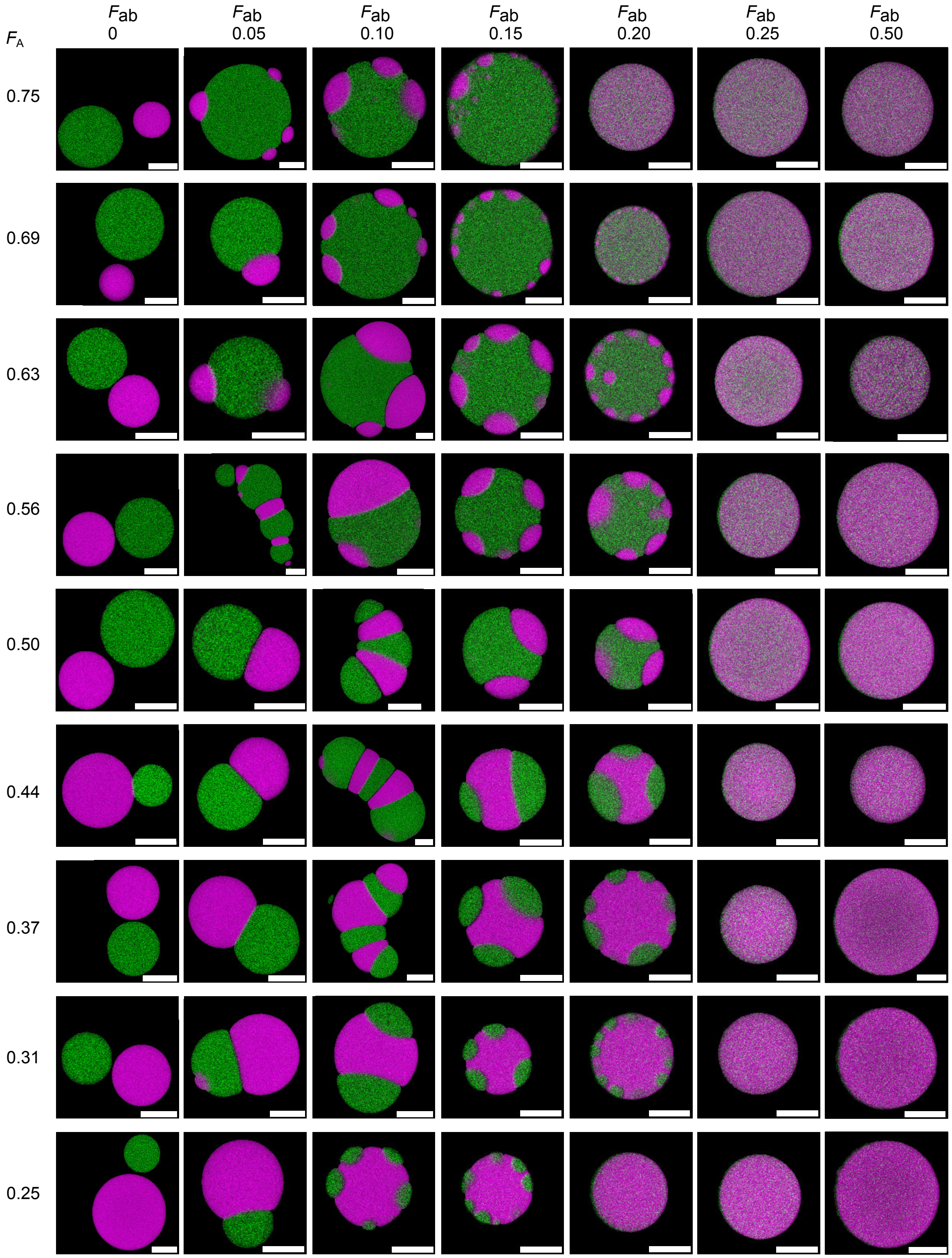


Figure S4: **Confocal micrographs used in the phase diagram in Figure 2.** Each row corresponds to samples at a given $F_\mathrm{A}$, while columns have constant $F_\mathrm{ab}$. DNA condensates were cropped out of larger fields of view and a mask was applied using the image segmentation process outlined in the Experimental Section to remove the background. The two channels were merged, with the A-rich phase shown in green (labeled with ATTO 488) and the B-rich phase shown in magenta (labeled with Alexa 647). All scale bars are 10 µm.

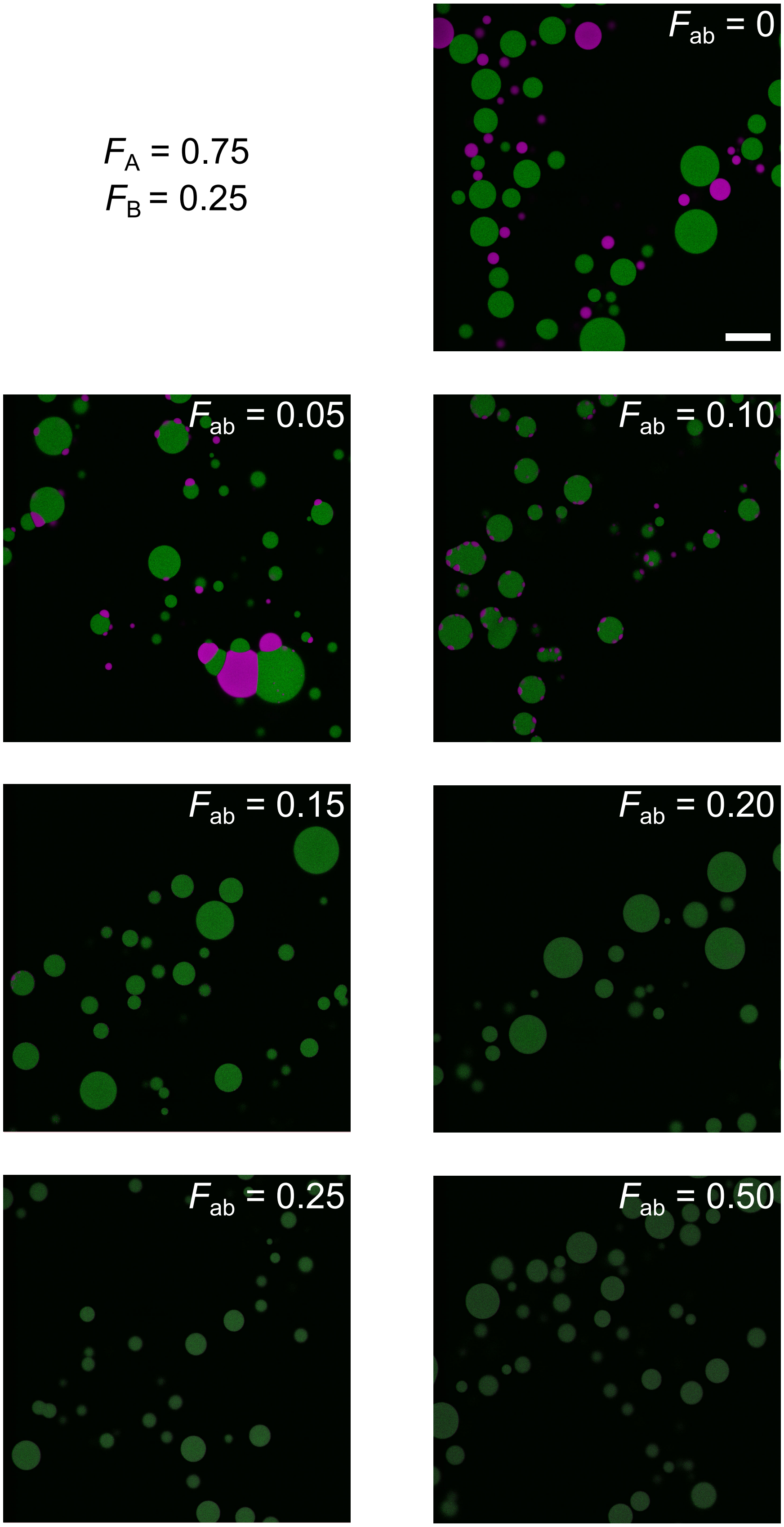


Figure S5: **Representative full fields of view for samples highlighted in Figure 2 and analyzed in Figure 3 with $F_A = 0.75$.** Images are pristine and uncropped with no background removed. Scale bar, 50 μm.

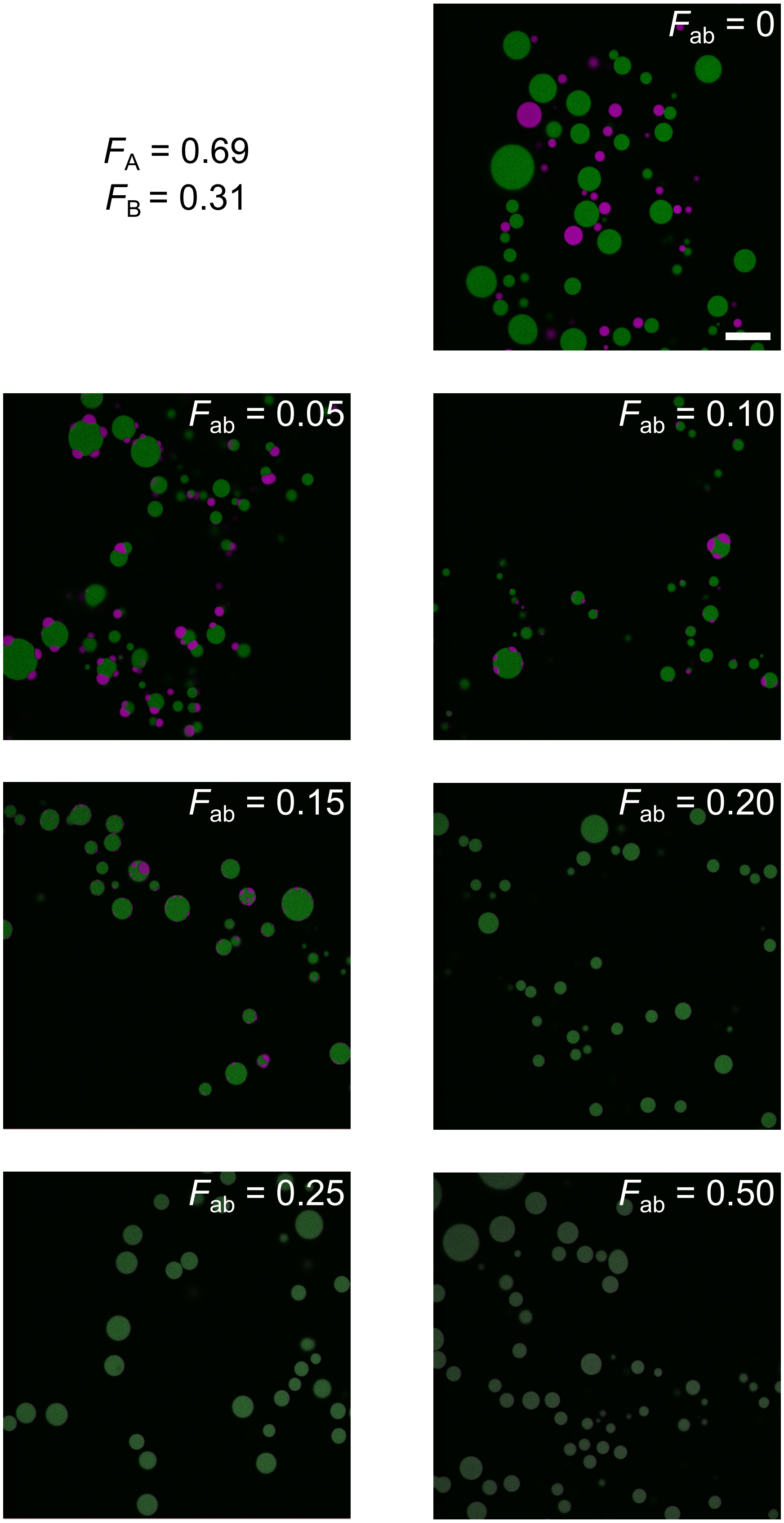


Figure S6: **Representative full fields of view for samples highlighted in Figure 2 and analyzed in Figure 3 with $F_A$** $= 0.69$. Images are pristine and uncropped with no background removed. Scale bar, 50 µm.

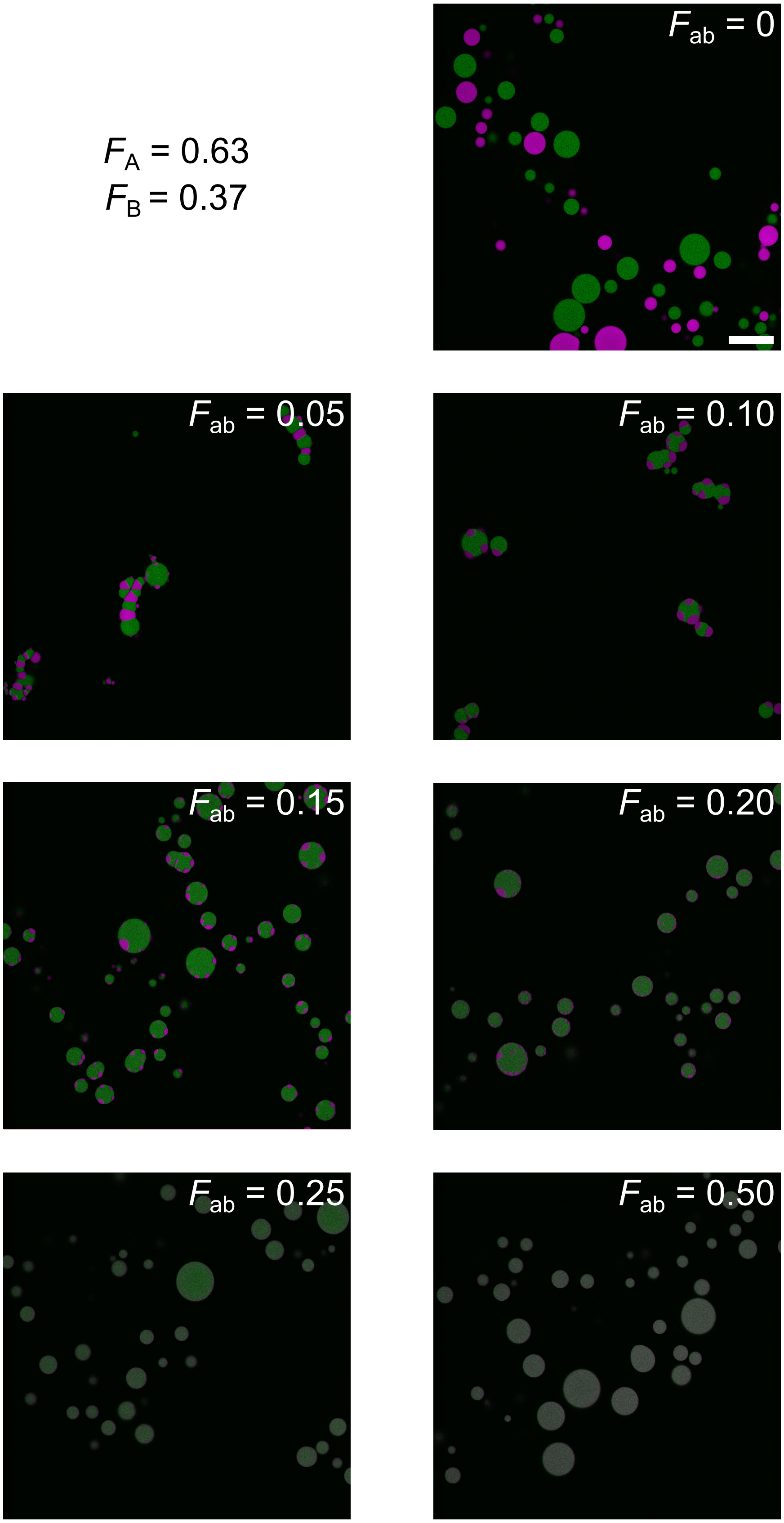


Figure S7: **Representative full fields of view for samples highlighted in Figure 2 and analyzed in Figure 3 with** $F_A = 0.63$. Images are pristine and uncropped with no background removed. Scale bar, 50 µm.

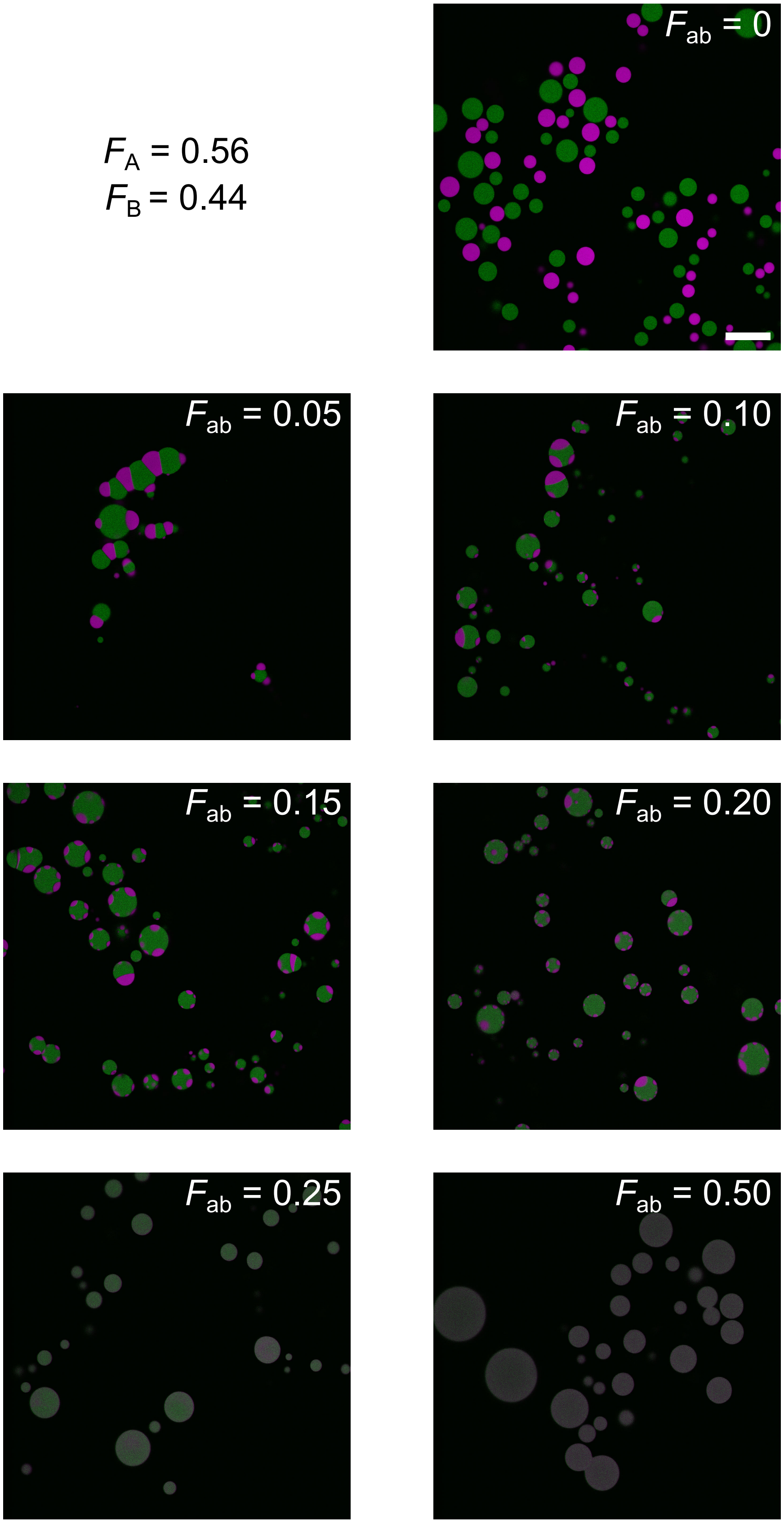


Figure S8: **Representative full fields of view for samples highlighted in Figure 2 and analyzed in Figure 3 with $F_A = 0.56$.** Images are pristine and uncropped with no background removed. Scale bar, 50 µm.

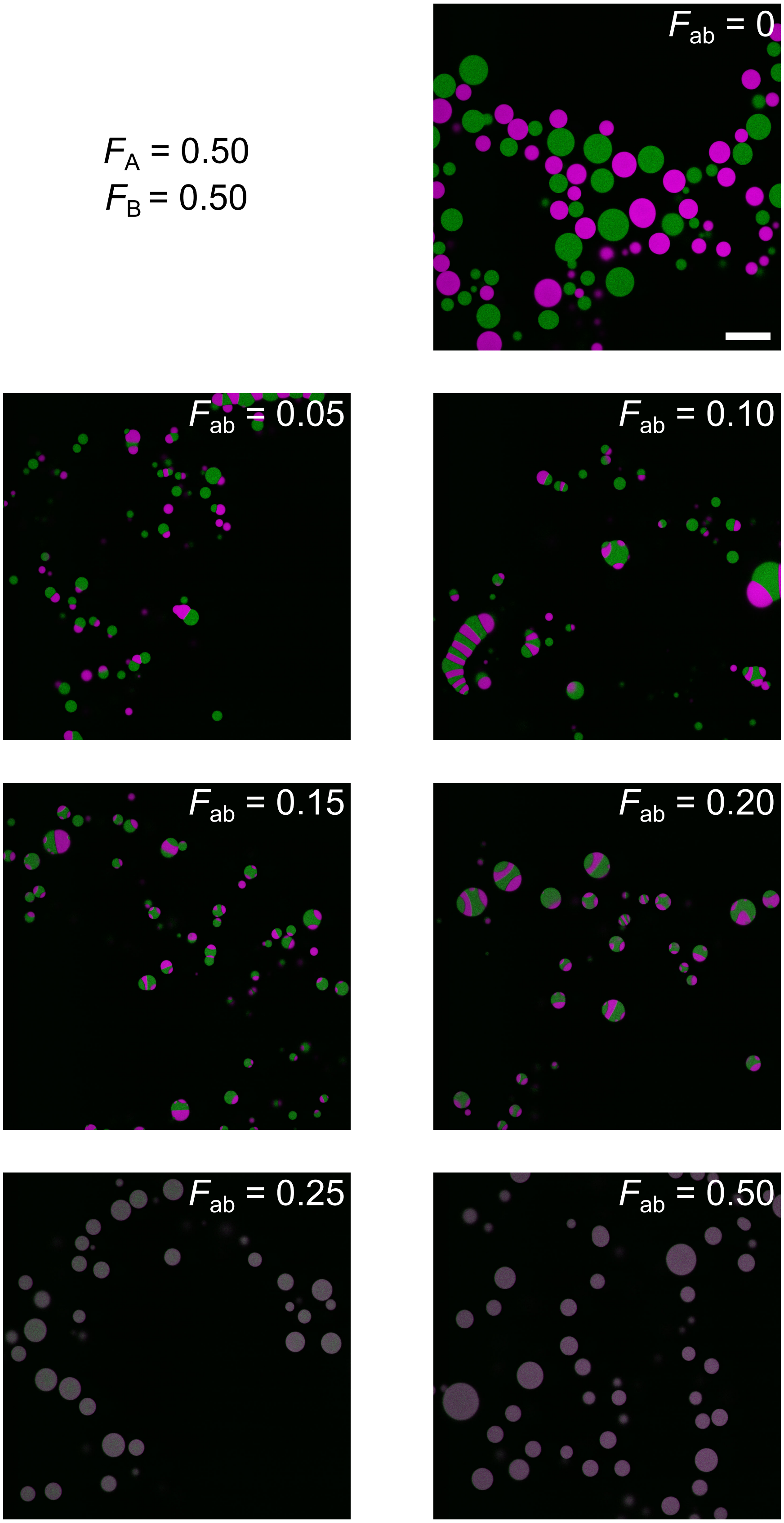


Figure S9: **Representative full fields of view for samples highlighted in Figure 2 and analyzed in Figure 3 with $F_A$ = 0.50.** Images are pristine and uncropped with no background removed. Scale bar, 50 µm.

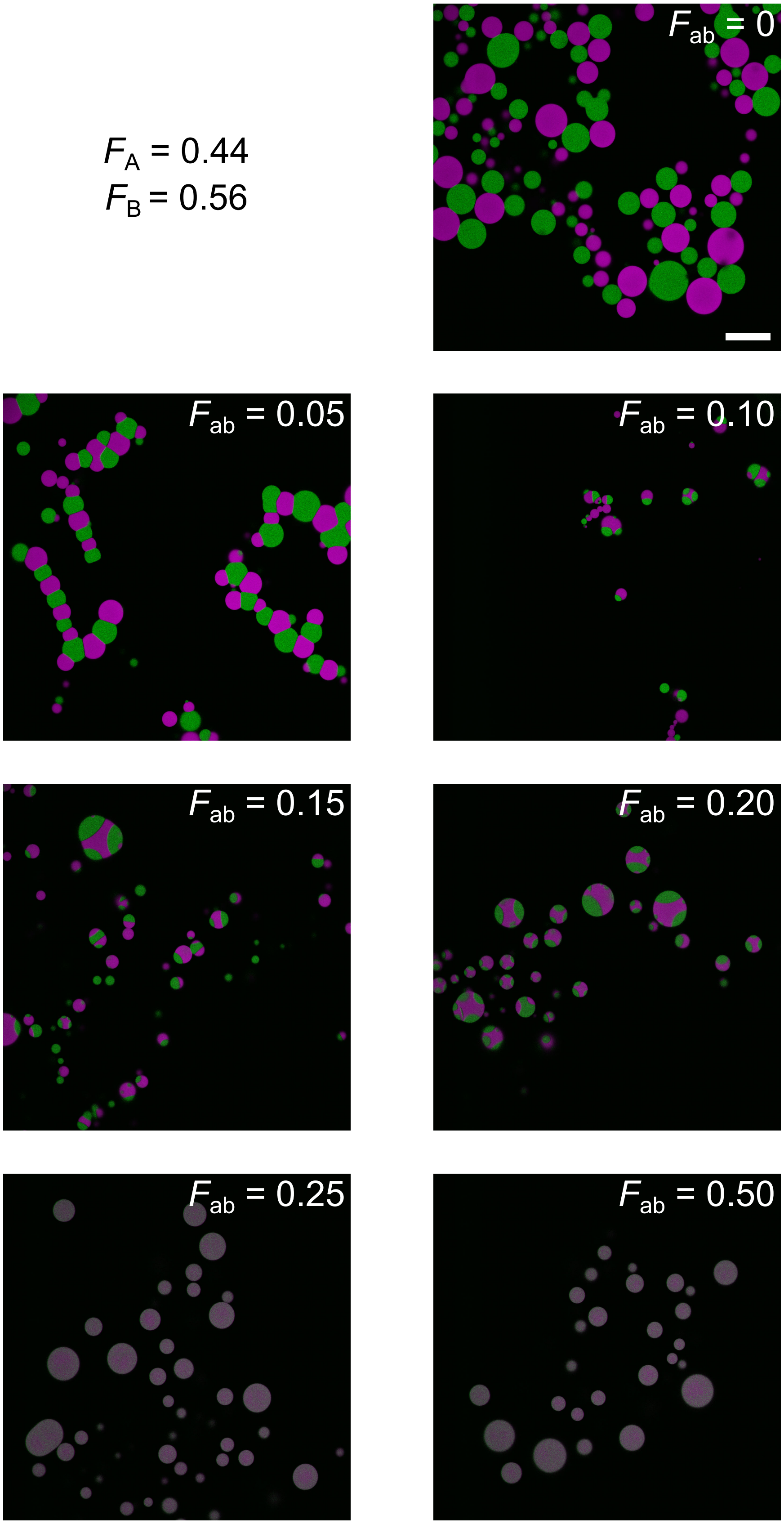


Figure S10: **Representative full fields of view for samples highlighted in Figure 2 and analyzed in Figure 3 with $F_A = 0.44$.** Images are pristine and uncropped with no background removed. Scale bar, 50 µm.

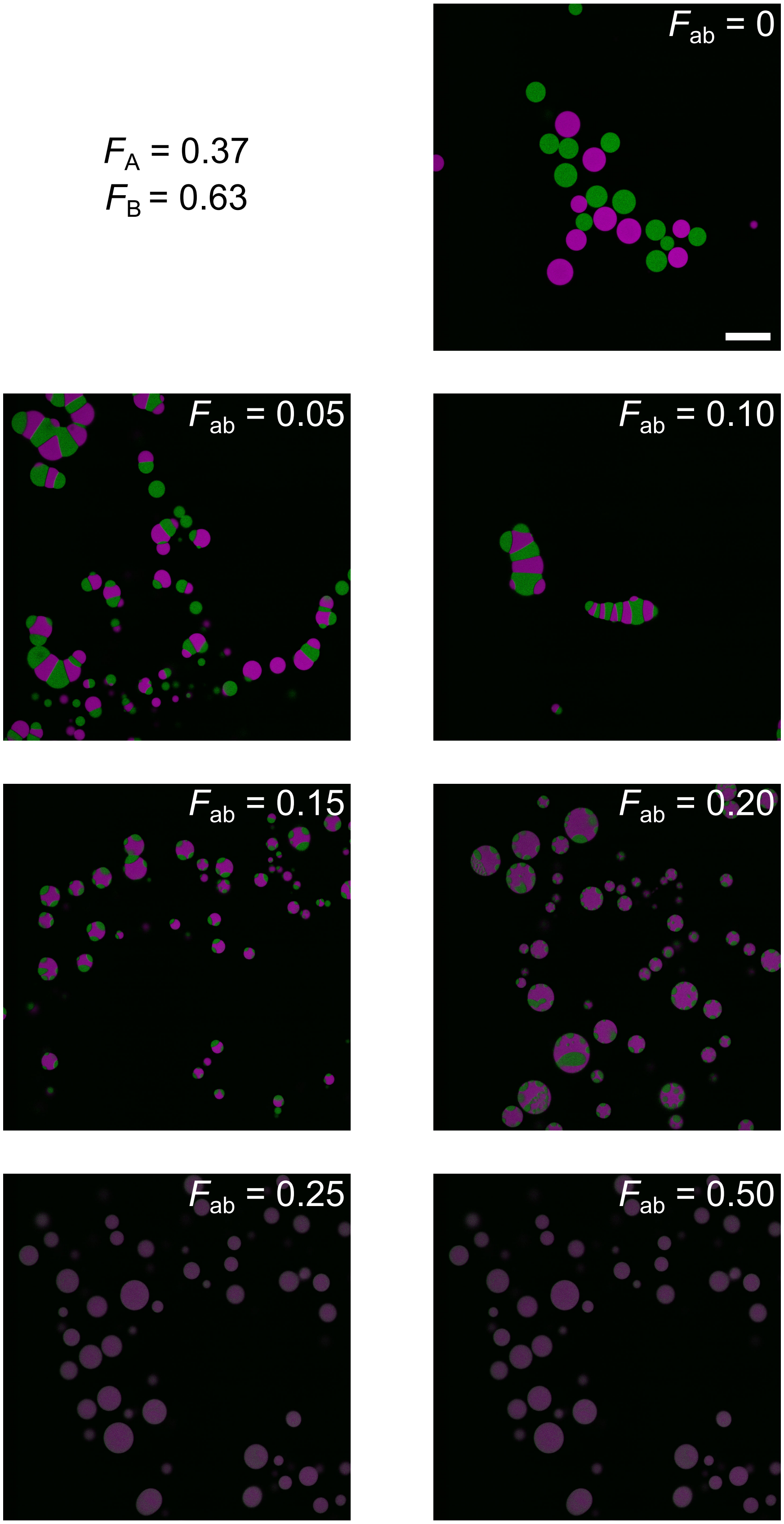


Figure S11: **Representative full fields of view for samples highlighted in Figure 2 and analyzed in Figure 3 with** $F_A = 0.37$. Images are pristine and uncropped with no background removed. Scale bar, 50 µm.

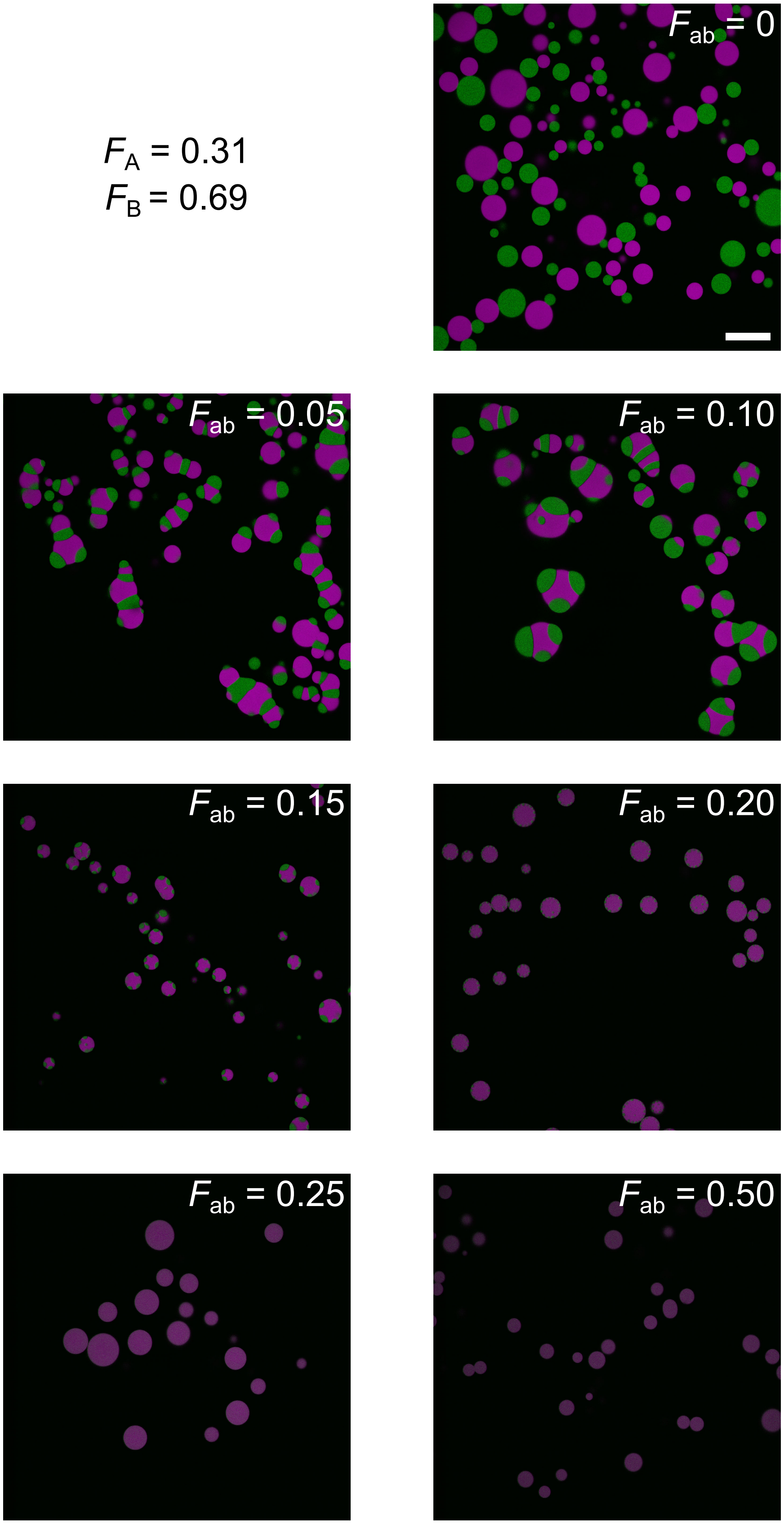


Figure S12: **Representative full fields of view for samples highlighted in Figure 2 and analyzed in Figure 3 with $F_A = 0.31$.** Images are pristine and uncropped with no background removed. Scale bar, 50 µm.

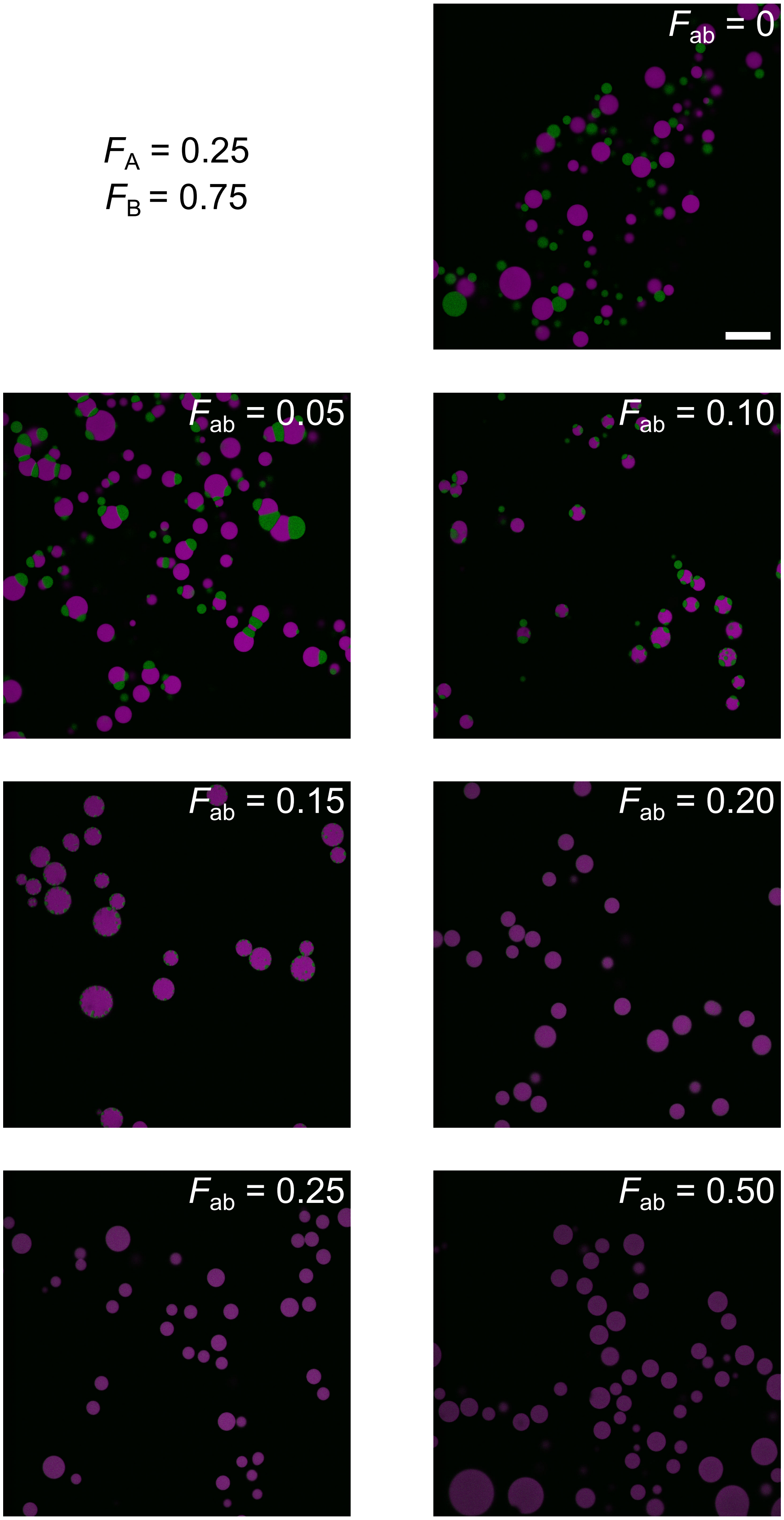


Figure S13: **Representative full fields of view for samples highlighted in Figure 2 and analyzed in Figure 3 with $F_A = 0.25$.** Images are pristine and uncropped with no background removed. Scale bar, 50 µm.

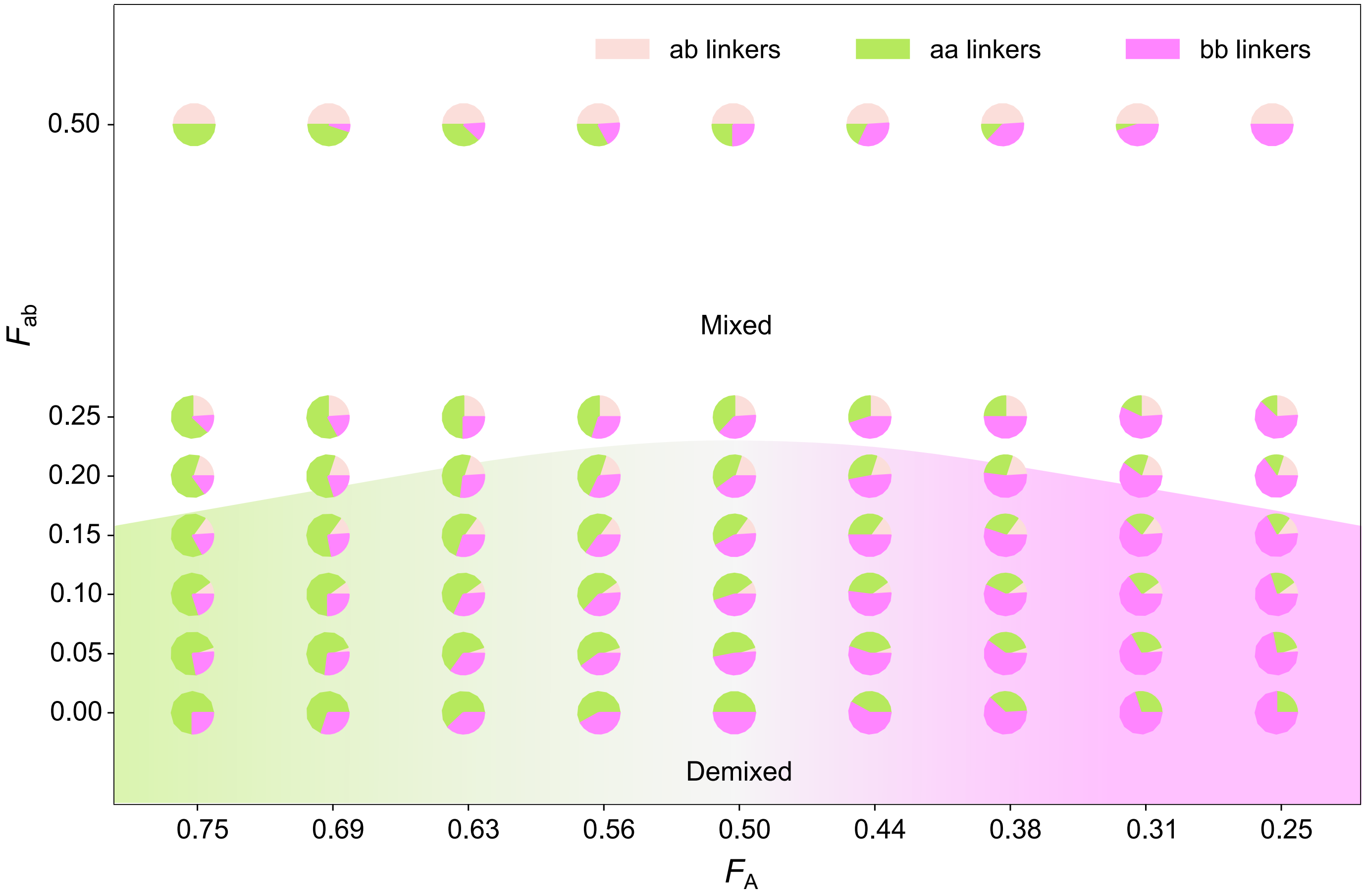


Figure S14: **Distribution of linkers in each sample tested and shown in Figure 2a**. Each pie chart marker represents the distribution of aa (green), bb (pink), and ab (cream) linkers for any given sample in the $F_A \times F_{ab}$ parameter space. Across each row, $F_{ab}$ stays constant, but the ratio between $F_{aa}$ and $F_{bb}$ changes proportionally with the fractions of nanostars $F_A$ and $F_B = 1 - F_A$ as specified on the x-axis.

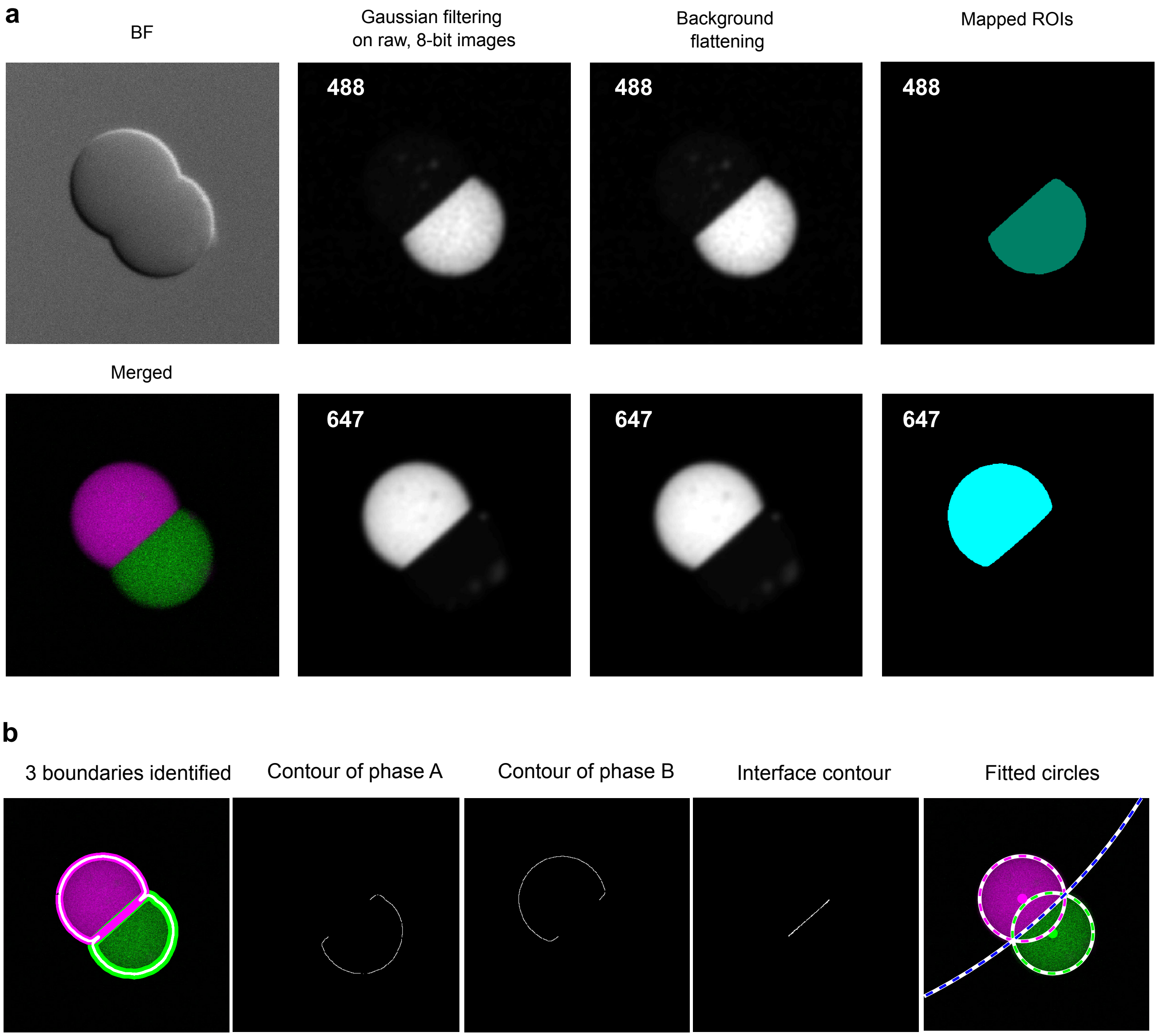


Figure S15: **Image segmentation steps undertaken for the determination of masks used to calculate partition coefficients and contact angles in condensates with a single interface.** Exemplified for DNA condensates made with $F_{\mathrm{ab}} = 0.05$ and annealed at constant 35°C [see Experimental Section for details]. The labels 488 and 647 refer to the fluorophores used to label NS A (ATTO 488) and NS B (Alexa 647). (**a**) Image processing steps for determining masks. (**b**) Boundary identification and circle fitting used for calculating contact angles.

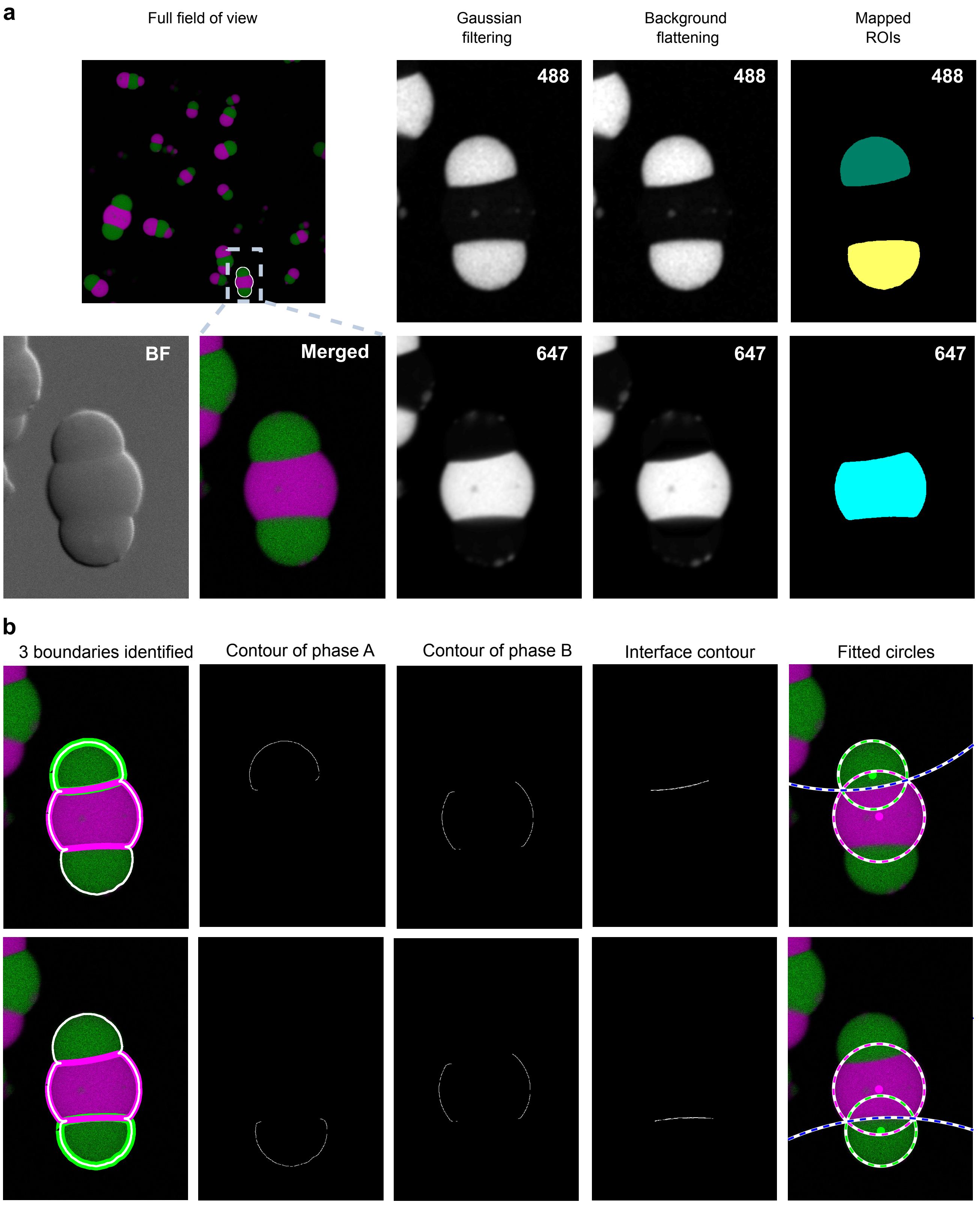


Figure S16: **Image segmentation steps undertaken for determining the masks used to calculate partition coefficients and contact angles in condensates with multiple interfaces.** In the example, DNA condensates are annealed in capillaries with $F_{ab} = 0.05$ and incubated at constant 35°C [see Experimental Section]. Condensates on the edge of the field of view are removed when determining segmentation masks.

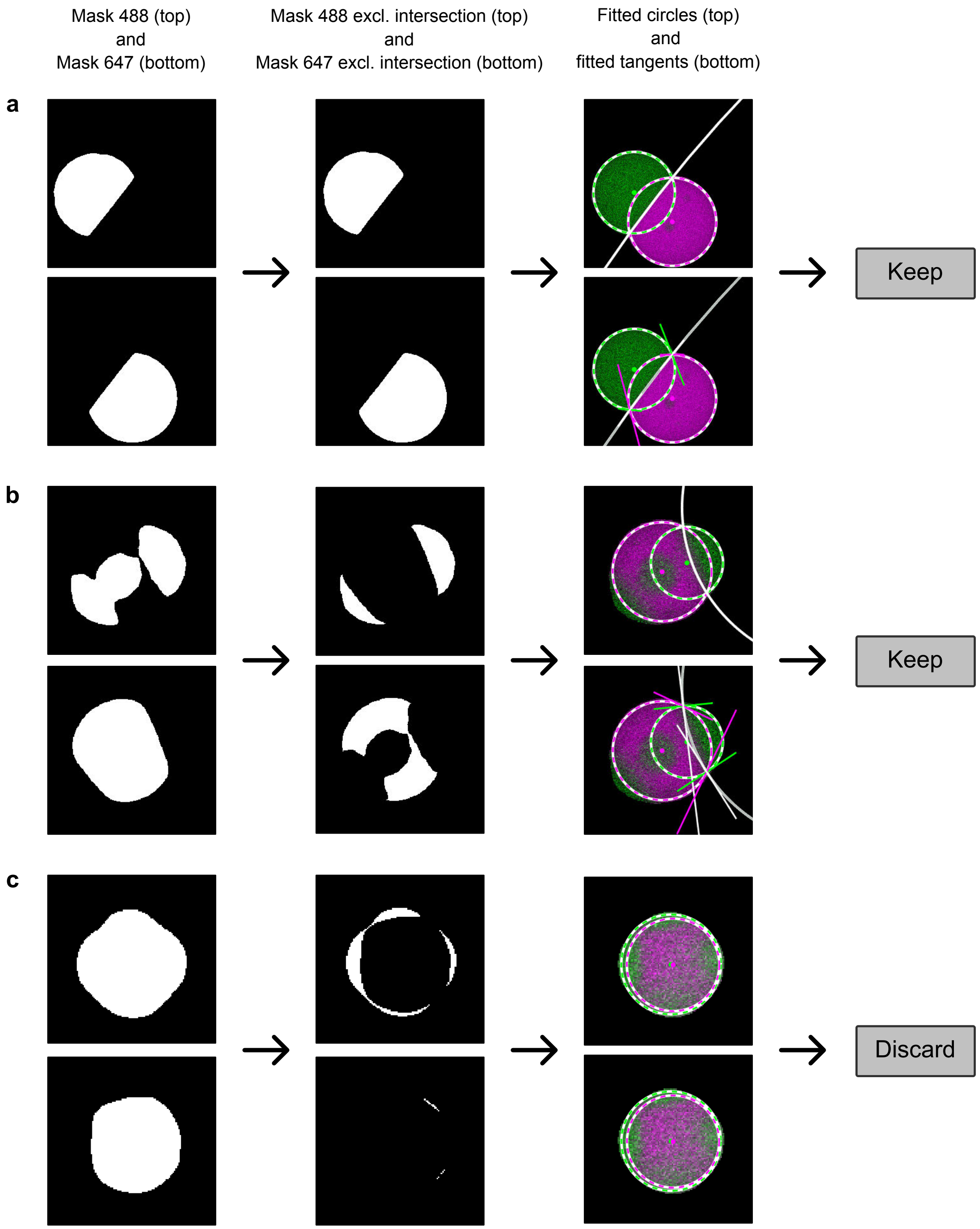


Figure S17: **Examples of the supervised segmentation approach used for biphasic condensates.** Following the segmentation routine described in the Experimental Section, the outcomes for biphasic samples were checked by the user. Contact angle measurements were only kept if the fitting of the circles and tangents was deemed to be accurate. Partition coefficients were recorded for all condensates. The three examples were selected from samples isothermally annealed in capillaries at 35°C. (**a**) Example of an easily segmented condensate for which the measurements were kept ($F_{\mathrm{ab}} = 0.05$). The initial masks for the two channels (left column) do not overlap significantly so the masks excluding the intersection area (middle column) are very similar to the initial masks.

(**b**) Example of a second, more challenging, condensate for which the measurements were kept ($F_{\mathrm{ab}}$ = 0.10). The initial masks for the two channels overlap in the region corresponding to an out-of-focus A-rich domain (green). The mask produced after removing this out-of-focus domain enables a good identification of the interfaces contact angles. (**c**) Example of a condensate which was discarded ($F_{\mathrm{ab}}$ = 0.15). Here the segmentation fails to identify domain contours due to significant interphase mixing.

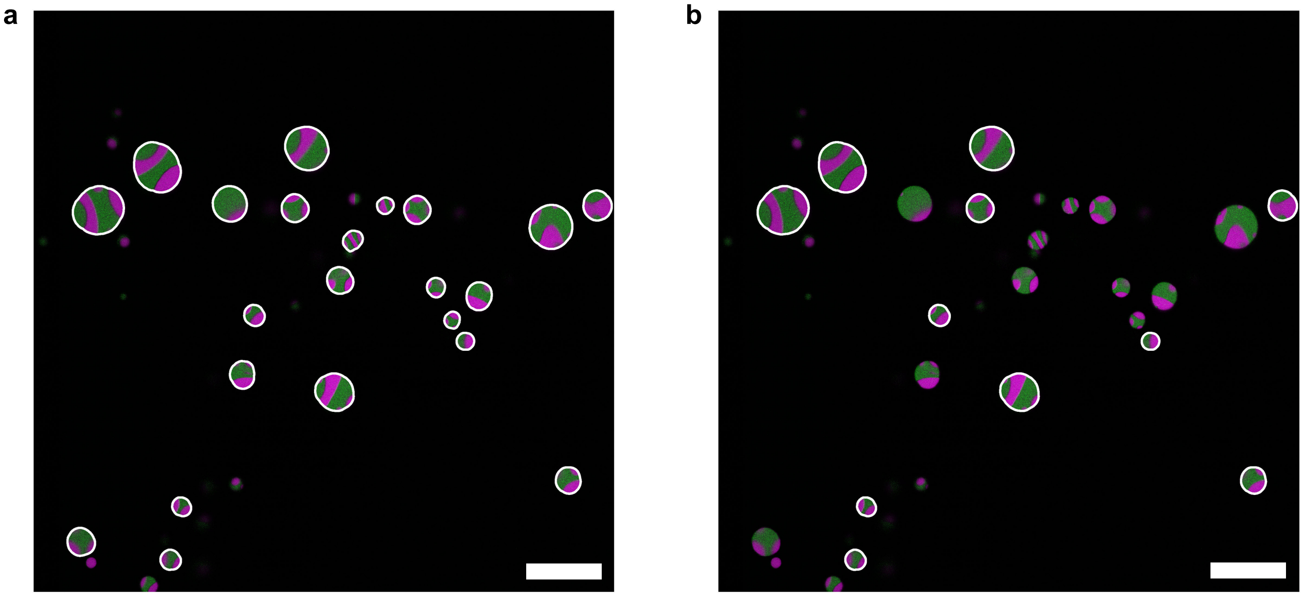


Figure S18: **Example of condensates selection for a sample with $F_A$ = 0.5 and $F_{ab}$ = 0.20.** (**a**) Condensates selected for partition coefficient measurements. Discarded condensates were below the size threshold or touching the edge of the field of view. (**b**) Condensates selected by the user for contact angle measurements based on the quality of the boundary identification as discussed in Figure S17. Scale bars, 50 µm.

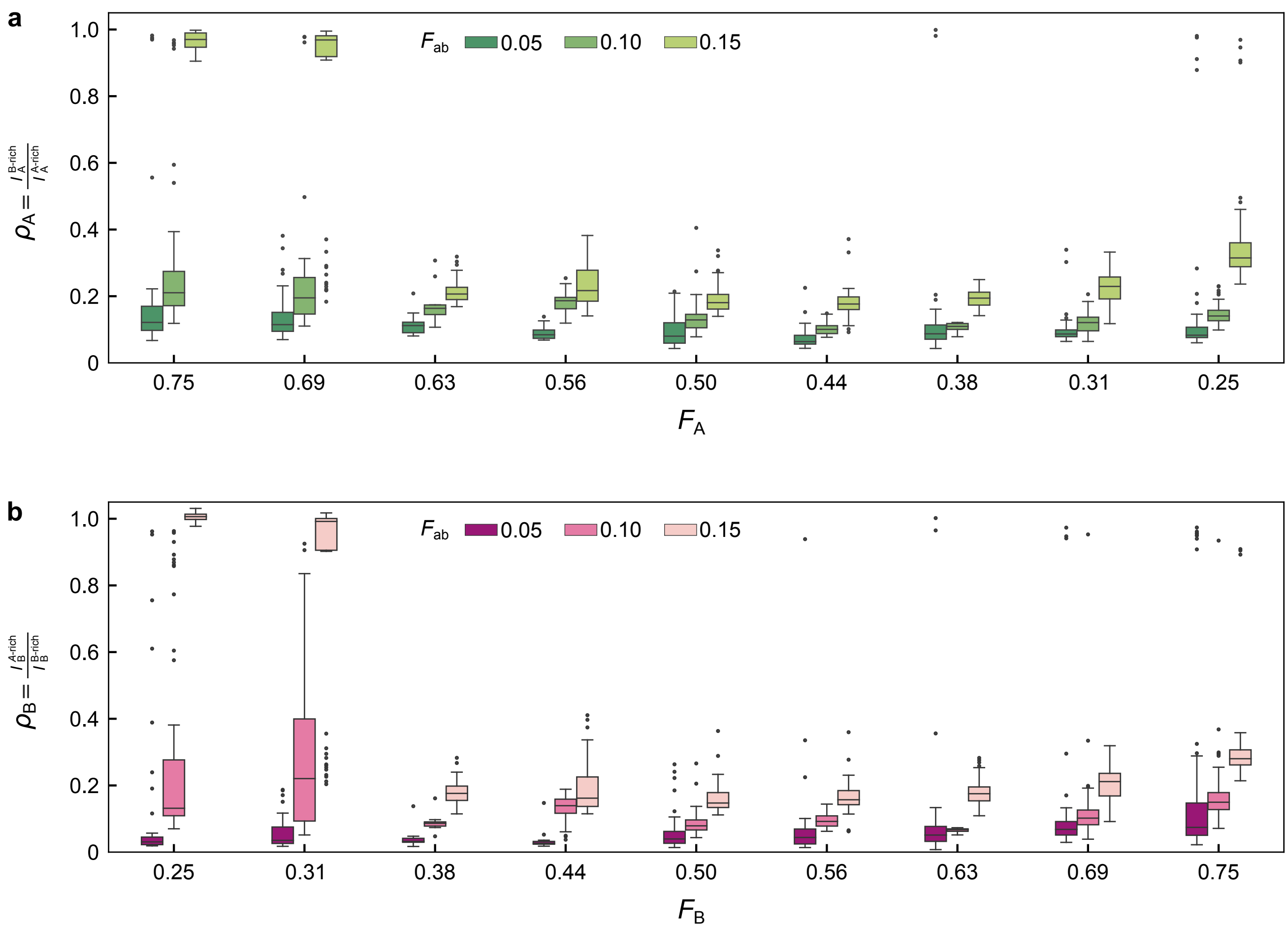


Figure S19: **Box plots of the partition coefficients as a function of NS molar fractions ($F_A$ and $F_B$) and fraction of ab linkers ($F_{ab}$).** (**a**) Partition coefficients for the A-rich phase, $\rho_A$. (**b**) Partition coefficients for the B-rich phase, $\rho_B$. See main text for definition of the partition coefficients. The median value is highlighted in each box and whiskers show the interquartile range, with outliers shown outside the whiskers. See corresponding micrographs in Figure 2 and Figures S5-S13. The number of measurements for each type of sample is provided in Table S3.

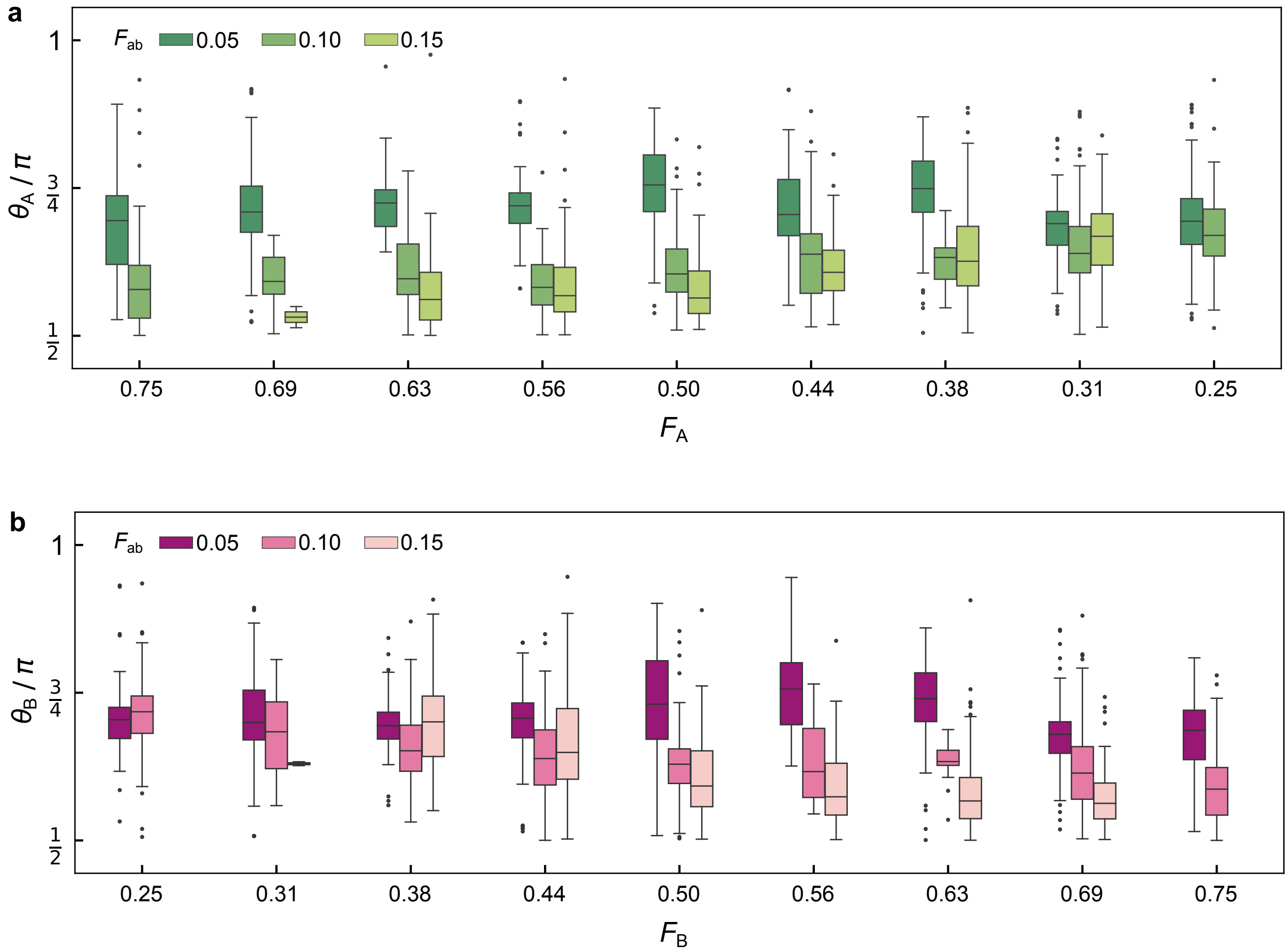


Figure S20: **Box plots of the contact angles as a function of NS molar fractions ($F_A$ and $F_B$) and fraction of ab linkers ($F_{ab}$).** (**a**) Contact angle for the A-rich phase, $\theta_A$. (**b**) Contact angle for the B-rich phase, $\theta_B$. See main text for definition of the contact angles. The median value is highlighted in each box and whiskers show the interquartile range, with outliers shown outside the whiskers. See corresponding micrographs in Figure 2 and Figures S5-S13. The number of measurements for each type of sample is provided in Table S4.

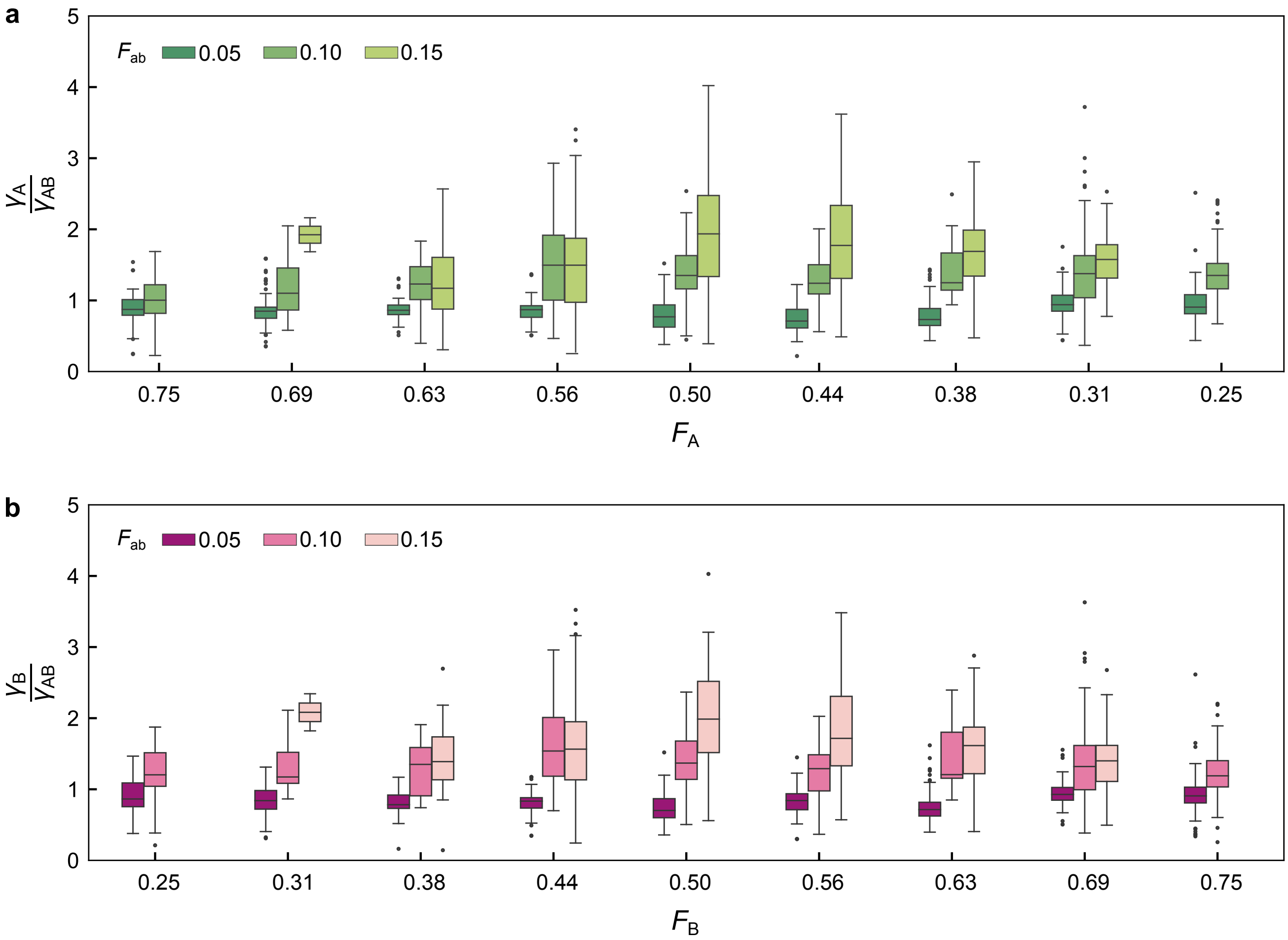


Figure S21: **Box plots of the interfacial tension ratios as a function of NS molar fractions ($F_A$ and $F_B$) and fraction of ab linkers ($F_{ab}$).** (**a**) Ratio between the surface tension for the A-rich phase and the A-B interfacial tension, $\gamma_A/\gamma_{AB}$. (**b**) Ratio between the surface tension of the B-rich phase and the A-B interfacial tension, $\gamma_B/\gamma_{AB}$. The median value is highlighted in each box and whiskers show the interquartile range, with outliers shown outside the whiskers. See corresponding micrographs in Figure 2 and Figures S5-S13. The number of measurements for each type of sample is provided in Table S4.

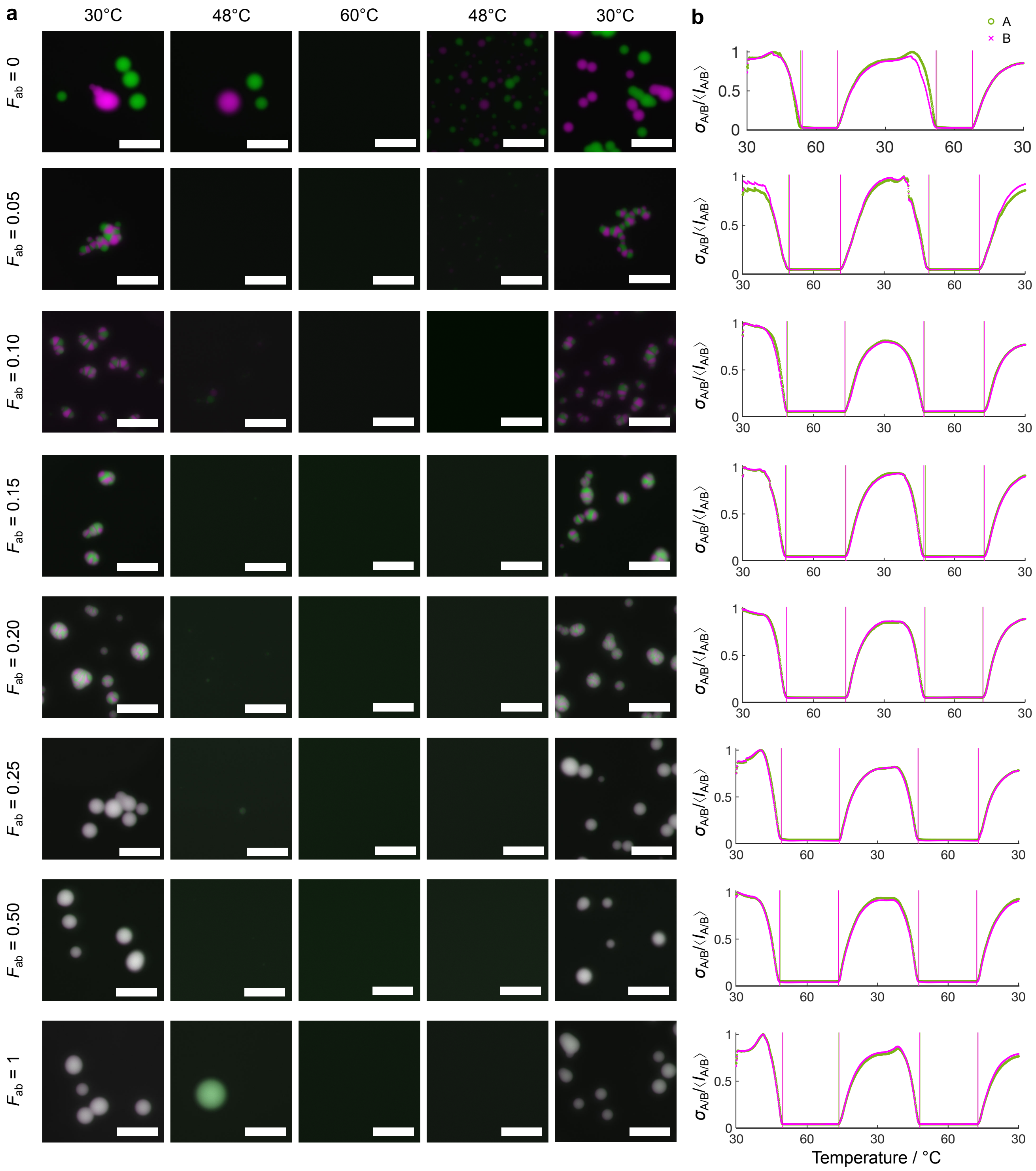


Figure S22: **Determination of condensate melting/assembly temperature ($T_m$).** (**a**) Selected epi-fluorescence snapshots for $F_{ab}$ between 0 and 1 as specified on the left, and $F_A = F_B = 0.5$. Images are shown at three representative temperatures on a heating ramp between 30°C - 60°C followed by a cooling ramp between 60°C - 30°C (± 0.5°C 15 min$^{-1}$). To facilitate visualization, micrographs were scaled from 0 to the maximum pixel intensity identified in the first frame before melting. Fluorescent channels shown in green and magenta are relative to A and B NSs, respectively. Scale bars 50 µm. (**b**) Plots of standard deviation of each field of view (FOV) normalized by the FOV mean as a function of temperature for samples in panel **a**.

Green and magenta curves correspond to data extracted from the A and B fluorescent channels, respectively. Transition temperatures on cooling and heating were determined from sharp changes as highlighted by vertical lines. Average $T_\mathrm{m}$ values as a function of $F_\mathrm{ab}$ are provided in Figure 4**b** and Table S5.

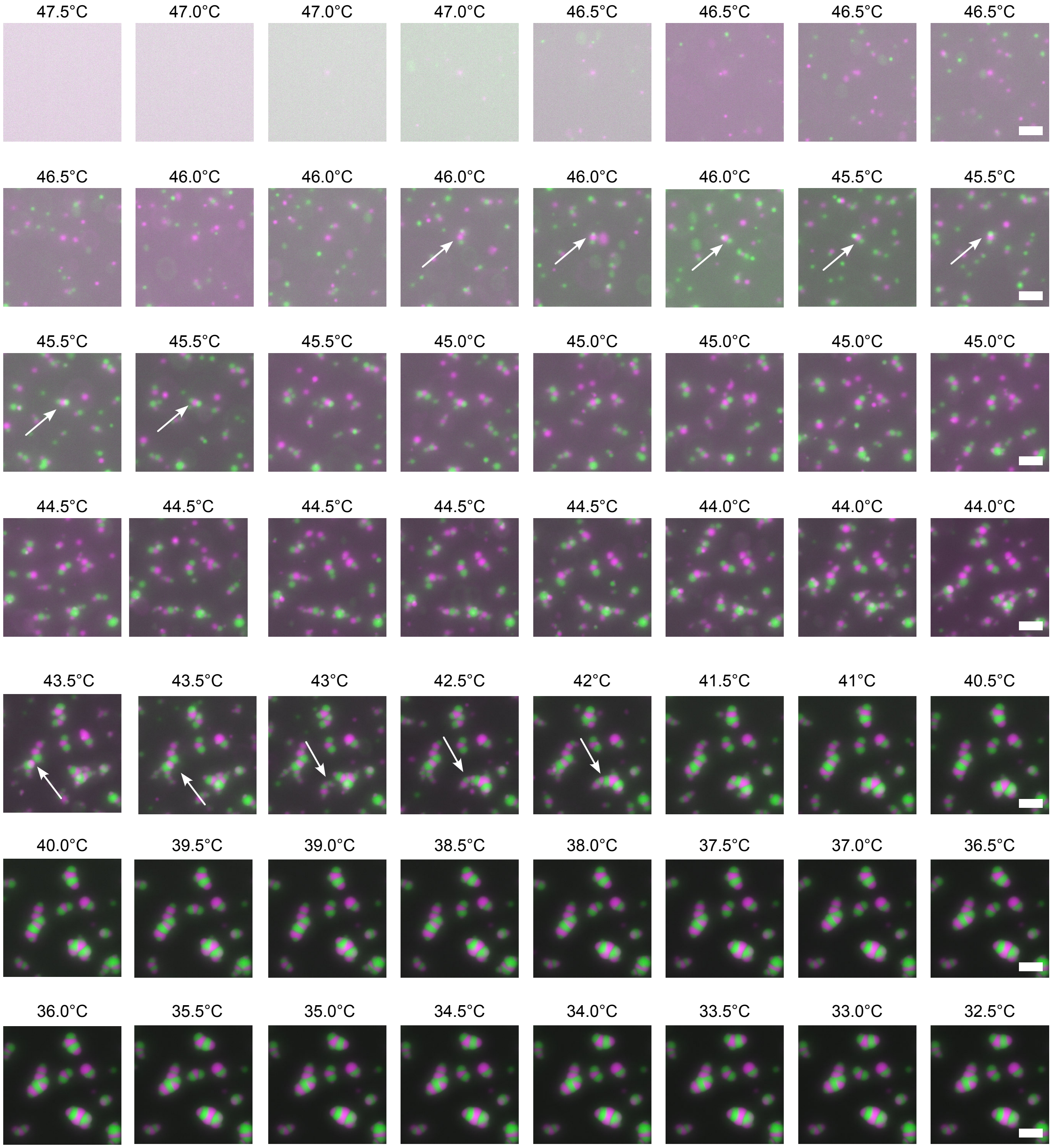


Figure S23: **Representative image sequences showing condensate formation and coalescence during a cooling ramp.** Some coalescence events are highlighted with white arrows. At early stages, many droplets appear to be monophasic, either A-rich or B-rich, later coalescing to form biphasic domains. The snapshots are selected from the first annealing cycle (60°C - 30°C) as described in the Experimental Section. Briefly, the cooling rate was set to -0.5°C for each 15 minutes, and the sample was imaged every 3 min resulting in 5 images at any one given temperature. Each image was scaled from 0 to maximum pixel intensity in each channel to aid visualization. Data are shown for $F_{\mathrm{ab}}$ = 0.10 and $F_{\mathrm{A}}$ = $F_{\mathrm{B}}$ = 0.5. Close to the temperature at which condensates start to form, multiple snapshots are shown at the same temperature for images acquired sequentially over time. All scale bars are 20 µm.

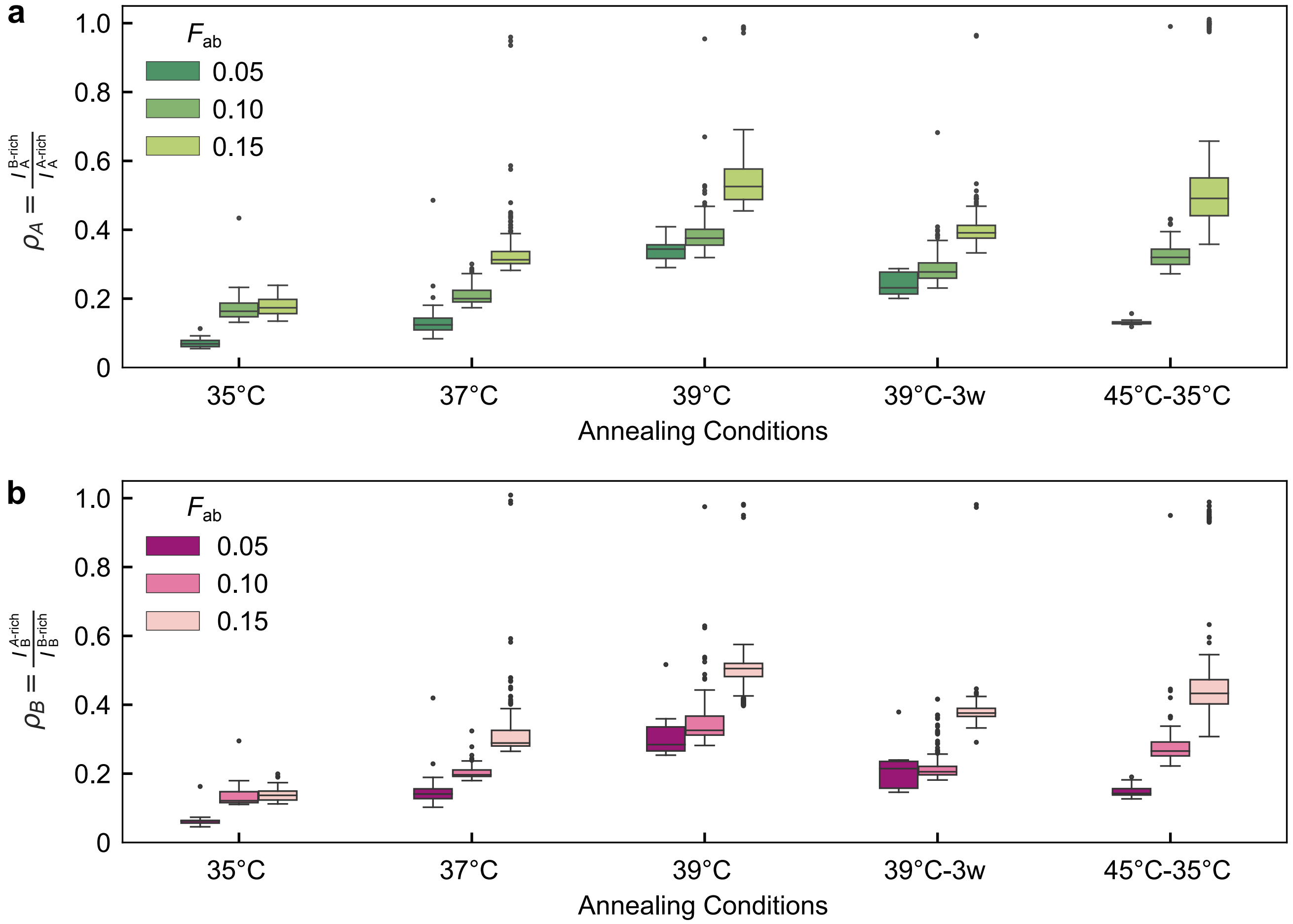


Figure S24: **Box plots of partition coefficients for samples annealed under five different conditions and shown in Figure 5.** For each annealing condition, measurements are shown for $F_{\text{ab}}$ = 0.05, 0.10, and 0.15. In samples with $F_{\text{ab}}$ = 0.20, 0.25, and 0.5, condensates were fully mixed and partition coefficients approaching 1 are not shown. (**a**) Partition coefficients $\rho_{\text{A}}$ for the A-rich phase. (**b**) Partition coefficients $\rho_{\text{B}}$ for the B-rich phase. The median value is highlighted in each box and whiskers show the interquartile range, with outliers shown outside the whiskers. The number of measurements for each type of sample is provided in Table S6.

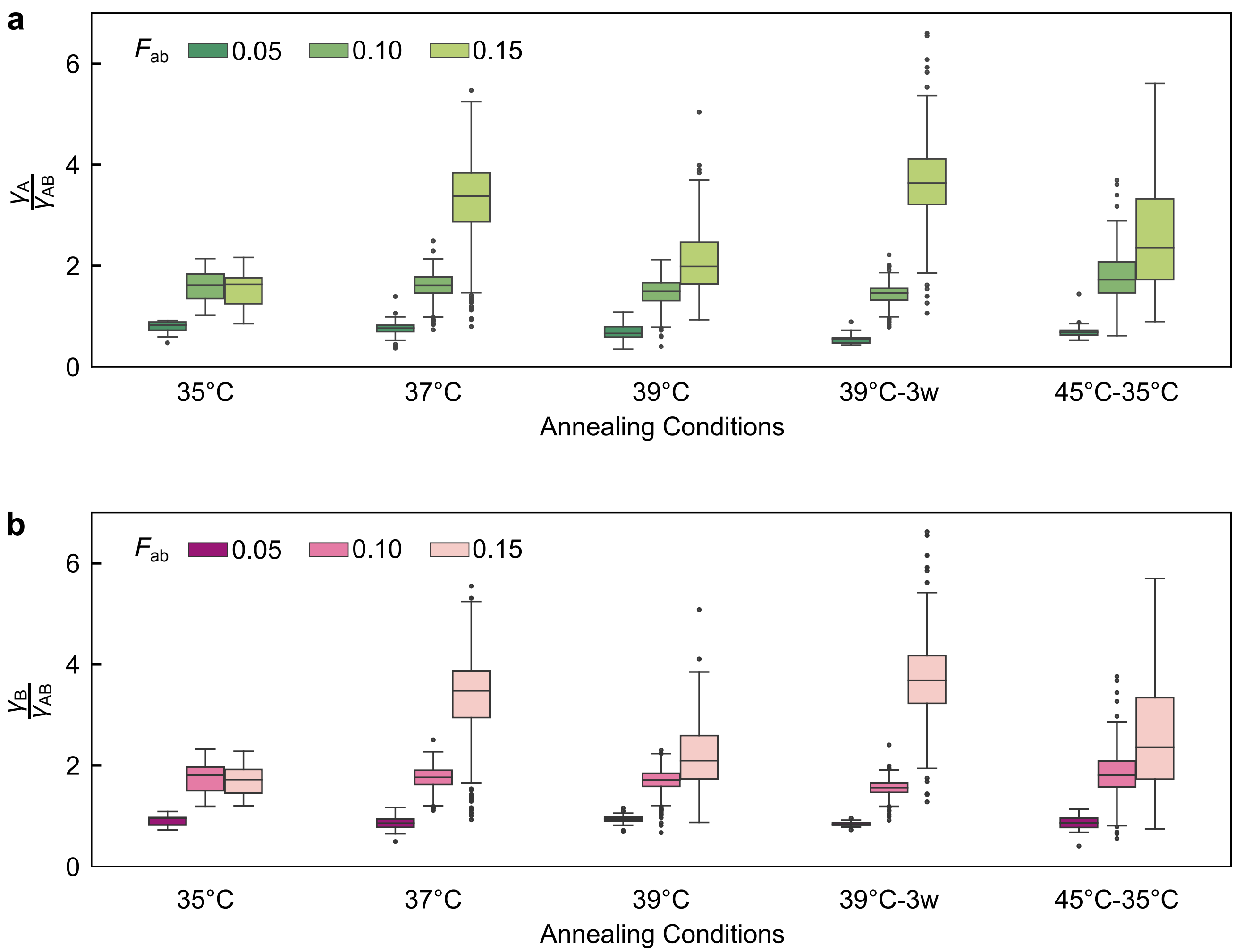


Figure S25: **Box plots of interfacial tension ratios for the five samples annealed under different conditions and shown in Figure 5.** (**a**) Ratio between the surface tension for the A-rich phase and the A-B interfacial tension, $\gamma_A/\gamma_{AB}$. (**b**) Ratio between the surface tension of the B-rich phase and the A-B interfacial tension, $\gamma_B/\gamma_{AB}$. The median value is highlighted in each box and whiskers show the interquartile range, with outliers shown outside the whiskers. The number of measurements for each type of sample is provided in Table S7.

### Supplementary Tables

Table S1: **Oligonucleotide sequences used for the preparation of nanostars and linkers.** Sticky ends (SEs) are emphasized in bold.

| Strand name | Sequence (5' → 3') |
|---|---|
| A4_core1 | GATCGCCGCCGCAATCACGCGCGTGCTCGGCGCCAGCAGTCCTGGCG |
| A4_core1_ATTO488 | GATCGCCGCCGCAATCACGCGCGTGCTCGGCGCCAGCAGTCCTGGCGT/3ATTO488N/ |
| A4_core2 | GATCGCCGCCAGGACTGCTGGCGCCGTCGCTTCTCTTCATAACAACG |
| A4_core3 | GATCGCCGTTGTTATGAAGAGAAGCGTCGCTCTGGCACAGGTGTACG |
| A4_core4 | GATCGCCGTACACCTGTGCCAGAGCGTGCACGCGCGTGATTGCGGCG |
| B4_core1 | GCGTGTGCTGTGCACTGTGAGGAGCGTCGCGTAACGTTCATTTGCCG |
| B4_core1_Alexa647 | GCGTGTGCTGTGCACTGTGAGGAGCGTCGCGTAACGTTCATTTGCCGT/3AlexF647N/ |
| B4_core2 | GCGTGTCGGCAAATGAACGTTACGCGTCGGCGTTGATCGAGTTAACG |
| B4_core3 | GCGTGTCGTTAACTCGATCAACGCCGTGCAGCGCTCGACACACGTCG |
| B4_core4 | GCGTGTCGACGTGTGTCGAGCGCTGCTCGCTCCTCACAGTGCACAGC |
| Linker_aa1 | GCGATCCGCAAACCAGCAAGCTCACG |
| Linker_aa2 | GCGATCCGTGAGCTTGCTGGTTTGCG |
| Linker_bb1 | ACACGCCGACATGCGTGCGGAGCGCG |
| Linker_bb2 | ACACGCCGCGCTCCGCACGCATGTCG |
| Linker_ab_a | GCGATCCGCCTGGTGGAACGAGCCGC |
| Linker_ab_b | ACACGCGCGGCTCGTTCCACCAGGCG |

Table S2: **Melting temperatures (mean) reported in °C ± standard deviation extracted from UV-Vis melting curves.** A and B represent nanostars samples and aa, bb, and ab represent linker samples made as described in Methods with total strands concentration 1.2 µM solubilized in 300 mM NaCl 1× TE. Data was collected for at least three technical replicates.

| | **$T_m$ cooling** | | **$T_m$ heating** | | **$T_m$ average** | |
|---|---|---|---|---|---|---|
| **A** | 73.5 | ± 0.3 | 74.3 | ± 0.4 | 73.9 | ± 0.5 |
| **B** | 73.0 | ± 0.2 | 74.5 | ± 0.4 | 73.7 | ± 0.9 |
| **aa** | 73.1 | ± 0.4 | 74.2 | ± 0.2 | 73.6 | ± 0.7 |
| **bb** | 77.3 | ± 0.3 | 78.3 | ± 0.5 | 77.8 | ± 0.6 |
| **ab** | 66.3 | ± 0.3 | 67.1 | ± 0.6 | 66.7 | ± 0.6 |

Table S3: **Number of partition coefficient measurements for each $F_A$ and $F_{ab}$.** Data is presented in Figure 3 for condensates shown with representative examples in Figure 2 and Figures S5-S13. Data was collected for three technical replicates.

| | | **$F_A$** | | | | | | | | |
|---|---|---|---|---|---|---|---|---|---|---|
| | | **0.75** | **0.69** | **0.63** | **0.56** | **0.5** | **0.44** | **0.38** | **0.31** | **0.25** |
| **$F_{ab}$** | **0.05** | 35 | 40 | 12 | 28 | 48 | 34 | 48 | 56 | 64 |
| | **0.10** | 51 | 44 | 12 | 24 | 29 | 16 | 10 | 66 | 73 |
| | **0.15** | 64 | 56 | 56 | 92 | 60 | 42 | 82 | 76 | 72 |
| | **0.20** | 56 | 66 | 50 | 48 | 56 | 52 | 98 | 58 | 66 |
| | **0.25** | 188 | 224 | 172 | 168 | 170 | 166 | 168 | 126 | 153 |

Table S4: **Number of contact angle measurements and interfacial tension ratio measurements for each $F_A$ and $F_{ab}$.** Note that not all compositions produced sufficiently de-mixed domains to determine contact angles and interfacial tension ratios from segmentation. Data is presented in Figure 3 for condensates shown with representative examples in Figure 2 and Figure S5-S13. Data was collected for three technical replicates.

| | | **$F_A$** | | | | | | | | |
|---|---|---|---|---|---|---|---|---|---|---|
| | | **0.75** | **0.69** | **0.63** | **0.56** | **0.5** | **0.44** | **0.38** | **0.31** | **0.25** |
| **$F_{ab}$** | **0.05** | 92 | 111 | 28 | 96 | 53 | 54 | 105 | 109 | 116 |
| | **0.10** | 97 | 19 | 17 | 28 | 62 | 34 | 13 | 138 | 125 |
| | **0.15** | | 2 | 55 | 84 | 75 | 54 | 130 | 75 | |
| | **0.20** | | | | 9 | 48 | 49 | 5 | | |

Table S5: **Transition temperatures in °C ± standard deviation as determined on cooling and heating ramps.** Transition temperatures for the A-rich phase and the B-rich phase were determined in each corresponding channel, in samples with varying $F_{\mathrm{ab}}$ and constant nanostar concentration. Values are presented as the average transition temperature extracted in three independently prepared samples, for a minimum of six cooling ramps and six heating ramps, respectively. The same median values are plotted in Figure 4**C**.

| $F_{\mathrm{ab}}$ | $T_{\mathrm{m}}$(*A* - cooling | $T_{\mathrm{m}}$(*A* - heating | $T_{\mathrm{m}}$(*B* - cooling | $T_{\mathrm{m}}$(*B* - heating |
|---|---|---|---|---|
| **0** | 49.9 ± 1.2 | 50.9 ± 2.6 | 49.9 ±1.3 | 50.8 ± 1.9 |
| **0.05** | 47.8 ± 1.0 | 49.3 ± 1.9 | 47.6 ± 1.0 | 49.2 ± 1.9 |
| **0.10** | 46.5 ± 1.4 | 48.9 ± 1.7 | 46.5 ± 0.9 | 48.8 ± 1.7 |
| **0.15** | 46.2 ± 1.0 | 48.3 ± 1.1 | 46.1 ± 0.7 | 48.0 ± 0.9 |
| **0.20** | 46.0 ± 1.2 | 48.6 ± 1.1 | 45.7 ± 0.9 | 48.0 ± 1.3 |
| **0.25** | 45.8 ± 1.3 | 48.1 ± 0.9 | 45.8 ± 0.8 | 47.8 ± 1.2 |
| **0.50** | 46.1 ± 1.4 | 47.8 ± 0.8 | 46.1 ± 1.0 | 47.6 ± 0.8 |
| **1.00** | 46.1 ± 1.5 | 48.8 ± 1.6 | 45.8 ± 1.1 | 48.5 ± 1.5 |

Table S6: **Number of condensates analyzed for each annealing condition, as a function of $F_{\mathrm{ab}}$, to determine partition coefficients**. Data are presented in Figure 6 for condensates shown with representative examples in Figure 5.

| | | **Annealing conditions** | | | | |
|---|---|---|---|---|---|---|
| | | **35°C** | **37°C** | **39°C** | **39°C-3w** | **45°C-35°C** |
| $F_{\mathrm{ab}}$ | **0.05** | 18 | 80 | 7 | 15 | 16 |
| | **0.10** | 49 | 203 | 254 | 258 | 140 |
| | **0.15** | 35 | 284 | 312 | 280 | 290 |
| | **0.20** | 147 | 149 | 283 | 255 | 699 |
| | **0.25** | 106 | 422 | 353 | 283 | 618 |
| | **0.50** | 110 | 556 | 300 | 271 | 655 |

Table S7: Number of interfacial tension ratio measurements for each annealing condition, by $F_{\mathrm{ab}}$. Data are presented in Figure S25 and micrographs of condensates are shown in Figure 5.

| | **35°C** | **37°C** | **39°C** | **39°C-3w** | **45°C-35°C** |
|---|---|---|---|---|---|
| **0.05** | 16 | 96 | 12 | 37 | 37 |
| **0.10** | 44 | 230 | 282 | 292 | 279 |
| **0.15** | 30 | 215 | 172 | 173 | 109 |

Table S8: **Sticky ends (SEs) free energies at different key temperatures.** Free energies were calculated with NUPACK [1] (2 µM, 0.3 M $Na^+$, Ensemble: All stacking, DNA Parameters: dna).

| **Temperature / °C** | $G_{\alpha-\alpha^*}$ / **kcal mol**$^{-1}$ | $G_{\beta-\beta^*}$ / **kcal mol**$^{-1}$ | $\Delta G$ / **kcal mol**$^{-1}$ |
|---|---|---|---|
| 20 | -9.57 | -10.85 | 1.28 |
| 25 | -8.98 | -10.25 | 1.27 |
| 30 | -8.4 | -9.65 | 1.25 |
| 35 | -7.81 | -9.06 | 1.25 |
| 37 | -7.58 | -8.83 | 1.25 |
| 39 | -7.34 | -8.6 | 1.26 |
| 40 | -7.23 | -8.48 | 1.25 |

[3] The MathWorks Inc. Matlab version: 24.1.0 (r2024a), 2024.

[4] Nikolai Chernov. Circle fit (pratt method), 2009.

[14] Karuna Skipper and Shelley Wickham. Core-shell coacervates formed from DNA nanostars. 2024. chemrxiv:10.26434.
